# Programmable conjugative CRISPR interference targeting genotoxin in the gut

**DOI:** 10.1101/2025.10.13.682122

**Authors:** Brian Hamp, Hania Timek, Chengyuan Fang, Zongqi Wang, Amber E. Haley, Maria Jennings, Collins I. Chimezie, Noah Hutchinson, Zhihong Dong, Shannon McCollum, Saleh Khawaled, Yatrik M. Shah, Omer Yilmaz, Jiahe Li

**Affiliations:** Department of Biomedical Engineering, College of Engineering and School of Medicine, University of Michigan, Ann Arbor, MI, 48109, United States; Department of Esophageal and Mediastinum, Harbin Medical University Cancer Hospital, Harbin, Heilongjiang, 150001, China; Program in Chemical Biology, Life Sciences Institute, University of Michigan, Ann Arbor, MI, 48109, United States; Cellular and Molecular Biology Program, University of Michigan, Ann Arbor, MI, 48109-5622, United States; Koch Institute for Integrative Cancer Research, Massachusetts Institute of Technology, Cambridge, MA, 02139, United States; Department of Internal Medicine, Division of Gastroenterology, University of Michigan Medical School, Ann Arbor, MI, 48109, United States; Department of Molecular and Integrative Physiology, University of Michigan Medical School, Ann Arbor, MI, 48109, United States; University of Michigan Rogel Cancer Center, University of Michigan Medical School, Ann Arbor, MI, 48109, United States; Department of Biology, Massachusetts Institute of Technology, Cambridge, MA, 02139, United States

**Author notes:** Correspondence: Jiahe Li, Department of Biomedical Engineering, College of Engineering and School of Medicine, University of Michigan, Ann Arbor, MI, 48109-5622.

## Abstract

Among microbially derived metabolites that influence host disease, colibactin garners increasing attention for its roles in the rising incidence of early-onset colorectal cancer. Produced by *pks*⁺ *Escherichia coli*, colibactin is a potent genotoxin, yet no approved therapeutics directly suppress it. Here, we engineered a self-transmissible conjugative plasmid to deliver CRISPR interference (CRISPRi) into multiple *pks⁺* strains. This system silences transcription of colibactin biosynthetic genes and abolishes *pks*⁺ *E. coli* genotoxicity without the resistance mutations associated with wild-type Cas9-mediated bacterial inhibition. In mice, conjugation-mediated CRISPRi reduces DNA damage and *pks*⁺ *E. coli* colonization while preserving commensal diversity. Importantly, the system also lowers tumorigenesis driven by *pks*⁺ *E. coli* and outperforms a pharmacologic inhibitor in a mouse colorectal cancer model. Finally, we extend this platform to silence a second pathogenic metabolite, establishing a translational strategy to neutralize diverse microbial metabolites and expanding the toolkit for programmable live biotherapeutics in the gut.

## Introduction

The human microbiome modulates host physiology through discrete microbial genes and metabolites, yet correlative studies linking taxa to disease rarely establish a mechanism or deliver therapeutic interventions ^1^. An emerging priority is to pinpoint and directly modulate microbial genes and metabolites that causally influence host biology. Classic reductionist approaches (gene knockouts or overexpression in germ-free mice and organoid co-cultures) have been invaluable ^2–5^, but they can prove inadequate when metabolite production depends on microbiome-host interplay or when conserved biosynthetic clusters are shared across species. To address these limitations, a complementary strategy is directly targeting specific microbial genes and metabolites *in situ* without affecting other commensal bacteria. Translationally, this strategy presents opportunities to neutralize pathogenic metabolites associated with various host diseases. Among deleterious microbial metabolites, colibactin from *pks*⁺ *Escherichia coli* has emerged as a driver of colorectal carcinogenesis, particularly in early-onset colorectal cancer (EOCRC), where incidence is rising in individuals under 50 ^3,6–10^. Colibactin is a hybrid polyketide-nonribosomal peptide encoded by the 54-kb polyketide synthase (*pks*) island. It was first reported in 2006 ^10^ and has been detected in pathogenic gut bacteria of over half of CRC patients ^11^. Two recent US prospective cohort studies of 134,775 participants underscored the link between a Western-style diet, *pks*^+^ bacteria, and CRC^12^. Despite its strong evidence for the causal relationship between colibactin and CRC, no approved therapies directly suppress colibactin production within the gut ecosystem.

In parallel, the clustered regularly interspaced short palindromic repeats (CRISPR) technology has revolutionized the landscape of genome editing ^13^, encompassing genomic cleaving ^14^, DNA editing ^15^, and DNA insertion ^16^. Beyond mammalian gene editing, CRISPR can also be repurposed to modulate the microbiome that colonizes the body, opening new avenues to interrogate host–microbe interactions and develop microbiome-targeted therapeutics ^17,18^. Here, we introduce the precision of CRISPR genome editing to directly inhibit colibactin directly in the native gut environment. We initially sought to employ wild-type Cas9 to selectively eliminate *pks*⁺ bacteria through targeted DNA cleavage due to the inefficient nonhomologous repair in bacteria. This strategy, however, proved ineffective as resistant clones emerged due to transposon-mediated inactivation of Cas9. We therefore harnessed a CRISPR interference (CRISPRi) approach, employing catalytically inactive Cas9 (dCas9) to silence colibactin biosynthetic genes without inducing bacterial death and transposon-mediated Cas9 inactivation. CRISPR-dCas9 effectively abolished genotoxicity while avoiding the mutational escape that undermined wild-type Cas9 killing. To achieve the delivery of CRISPR-dCas9 directly in the gut, we selected a self-transmissible conjugative plasmid, which naturally transfers across a broad range of *Enterobacteriaceae,* including *Escherichia*, *Citrobacter*, and *Klebsiella*, taxa frequently linked to genotoxin-mediated pathology ^19,20^. This promiscuous transfer ensures the dissemination of CRISPR-dCas9 modules to diverse *pks*⁺ strains, while specificity is dictated by the guide RNA-dCas9. To demonstrate the versatility of our platform, we extended this approach to silence yersiniabactin, another virulence-associated metabolite, which has been linked to inflammatory bowel disease ^21^. We deployed the CRISPR-dCas9 conjugative system to assess the translational potential in mouse models colonized with *pks*⁺ *E. coli* (**Figure 1**). This system reduces genotoxicity, diminishes *pks*⁺ colonization, suppresses tumorigenesis, and outperforms a benchmark small-molecule inhibitor without detectable systemic toxicity. Together, these data establish conjugative CRISPRi as a new strategy for *in situ* genetic neutralization of disease-driving microbial functions and a generalizable framework for programmable live biotherapeutics.

**Figure 1.**
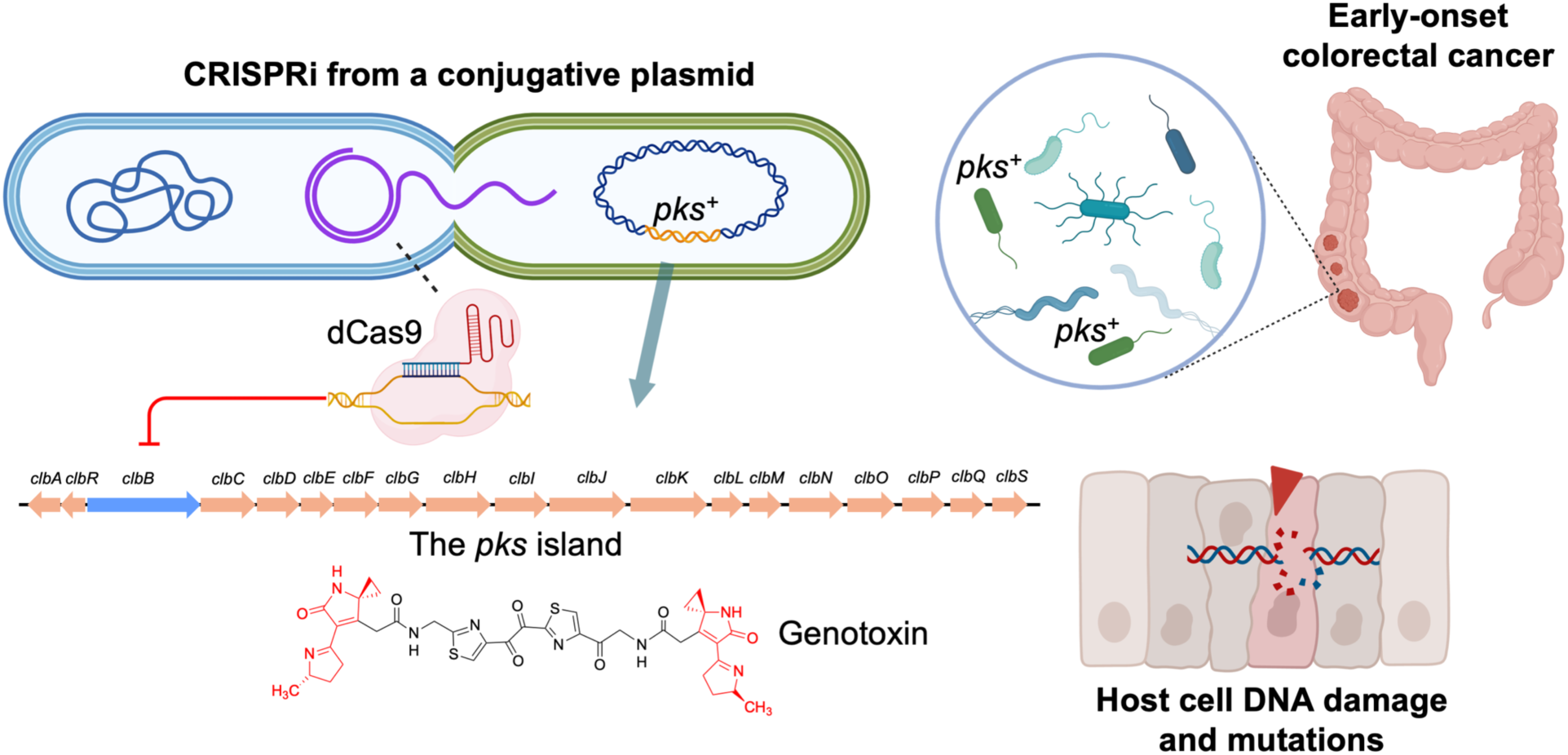
CRISPRi neutralizes colibactin and mitigates colorectal tumor initiation. CRISPRi utilizing catalytically deactivated Cas9 and a broad-host-range self-transmissible conjugative plasmid is capable of transcriptionally silencing the *pks* island in pathogenic bacteria and preventing colibactin-mediated tumorigenesis and colorectal cancer.

## Results

### Wild-type Cas9 fails to eliminate colibactin genotoxicity due to mutational escape

Synthetic biologists have long explored CRISPR-Cas9 for targeted bacterial killing, eliminating antimicrobial resistance (AMR) genes, or introducing kill switches for biocontainment purposes ^22,23^. These strategies leverage the fact that genomic cleavage is generally lethal to bacteria due to their inefficient nonhomologous DNA repair mechanisms ^22,23^. The *pks* island is highly conserved (∼100% identical at the nucleotide level) across different genera of *pks*^+^ bacteria ^19,24^, restricted to colibactin-producing bacteria, yet absent in non-genotoxic commensals. We reason that these attributes are well-suited for CRISPR-Cas9-mediated targeting tools to ensure precision editing of the microbial community by selectively eliminating *pks⁺* bacteria. This represents a paradigm shift from conventional antibiotic or probiotic interventions toward programmable, self-propagating microbial therapeutics.

The *pks* island encodes multiple highly conserved genes essential for colibactin biosynthesis ^19^. To this end, we first computationally designed three unique single guide RNAs (sgRNAs) targeting two genes (*clbA* and *clbB*), as knocking out *clbA* or *clbB* has been shown to eliminate colibactin biosynthesis and genotoxicity of *pks⁺* bacteria. We also selected a previously validated control sgRNA (sgCtl) to ensure the specificity of the CRISPR-Cas9 on the *pks* island ^25^. Each of these sgRNAs along with Cas9 constitutively expressed from its native promoter (denoted as pNative-Cas9) were cloned into a replicative plasmid with a low-copy replication origin (SC101) (**Extended Data Fig. 1a**). Since NC101 represents one of the best characterized adherent invasive *pks*^+^ *E. coli* for studying their pro-carcinogenicity in mice ^26–28^, we transformed the pNative-Cas9 plasmids into NC101 and plated them on selective LB agar for characterization. Introduction of pNative-Cas9-sg*clbA*_287, pNative-Cas9-sg*clbB*_4387, and pNative-Cas9-sg*clbB*_73 in NC101 markedly reduced the transformation efficiency relative to that of the control, pNative-Cas9-sgCtl (**Extended Data Fig. 2a**). In contrast, transformation efficiency in the isogenic mutant strain NC101Δ*pks* (with the *pks* island deleted) was comparable across all sgRNA constructs (**Extended Data Fig. 2b**). These results indicate that Cas9 paired with *pks*-targeting sgRNAs can selectively inhibit *pks⁺* bacteria while sparing the isogenic strains lacking the *pks* island, underscoring the selectivity of CRISPR-Cas9 against *pks⁺ E. coli*.

Although encouraging, pNative-Cas9 with *pks*-targeting sgRNAs achieved only partial elimination of *pks*⁺ NC101 (**Extended Data Fig. 2a**). Incomplete killing raises the concern that these Cas9-resistant *pks*⁺ bacteria could expand and dominate, analogous to antibiotic resistance. We therefore asked whether these putative ‘resistant’ *pks*⁺ bacteria retained genotoxic activity. Because colibactin is chemically unstable, genotoxicity is typically assessed through indirect assays ^29–31^. As such, a phosphorylated variant of the H2AX histone (γH2AX), accumulating at sites of DNA double-strand breaks, is a common indirect biomarker for colibactin-induced DNA damage to host cells ^7,32–34^. As evidenced by γH2AX staining, *pks⁺* NC101 transformed with *pks*-targeting sgRNAs (sg*clbA*_287, sg*clbB*_4387, or sg*clbB*_73) retained genotoxicity comparable to that of the control sgRNA (**Figure 2a**). In contrast, genetic knockout of the *pks* island led to complete neutralization of the genotoxicity by *pks⁺ E. coli*. This surprising finding suggested that partial inhibition of bacteria via CRISPR-Cas9 failed to reduce the genotoxicity of *pks*⁺ *E. coli*. To understand why this strategy failed, prior studies reported that targeted genomic cleavage by Cas9 can lead to inactivation of the *cas9* gene by transposable elements (e.g., insertional sequences) from the *E. coli* genome ^35^. As a result, *pks⁺ E. coli* carrying the mutated *cas9* can dominate the bacterial population over time. To test this, we expanded *pks⁺* NC101 colonies transformed with the CRISPR-Cas module and compared *pks-*specific sgRNAs (sg*clbA*_287, sg*clbB*_4387, sg*clbB*_73) to the control sgCtl. Interestingly, whole plasmid sequencing showed that nearly all clones carrying *pks*-specific sgRNAs had insertional sequences in the *cas9* open reading frame (**Figure 2b, Extended Data Fig. 3, and Table S1**). Since insertional sequences contain their own stop codon and introduce out-of-frame insertions, they can effectively abolish the translation of Cas9 prematurely, explaining the escape from Cas9-mediated killing. Therefore, we conclude that transposable insertional sequences from the *E. coli* genome provide a mutational escape route for *pks⁺ E. coli* from Cas9-mediated killing, resulting in reduced growth inhibition and genotoxicity of *pks⁺ E. coli*.

**Figure 2.**
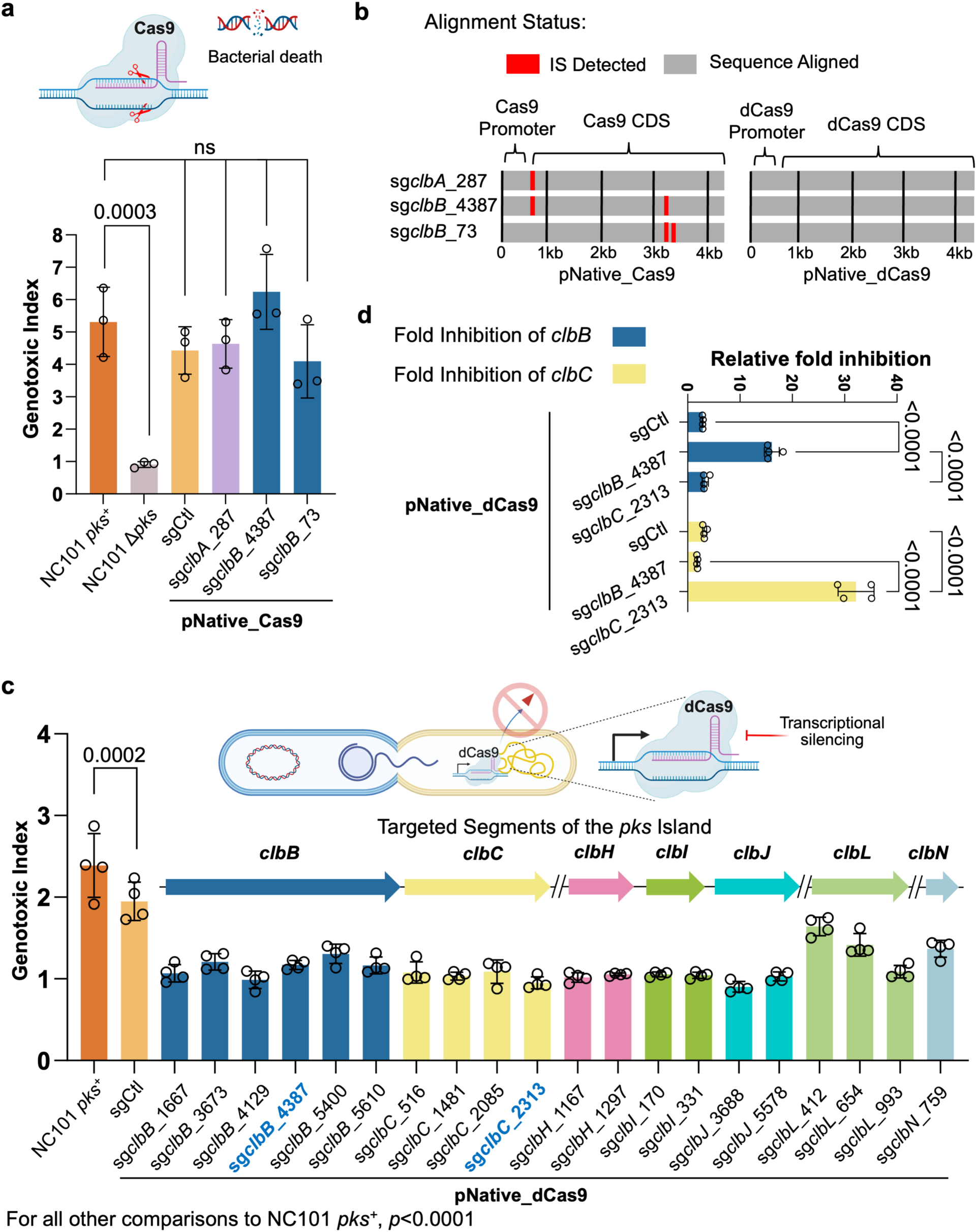
Design and optimization of dCas9 guides for transcriptional silencing of colibactin biosynthetic genes. **a**, Limitations of wild-type Cas9 in targeting the *pks* island. *pks*^+^ NC101 *E. coli* was transformed with a plasmid encoding wtCas9 driven by a native promoter (pNative) and corresponding sgRNAs yet failed to lower the genotoxicity in HeLa cells following 4 h of co-culture as compared to positive controls. The genotoxic index is calculated by normalization against the median fluorescence index of HeLa cells cultured without bacteria present. Multiplicity of infection (MOI) = 100, n = three technical repeats per independent biological replicate, N = two independent biological replicates. **b**, Inactivation of wtCas9 by insertional sequences. Sequence alignment of pKill1 plasmids extracted from transformed *pks*^+^ NC101 *E. coli* shows deleterious insertional sequences (not shown to scale) in the promoter and Cas9 regions, but no insertional sequences for the dCas9 versions of these constructs. **c**, Experimental validation of *in silico* designed sgRNAs targeting different transcriptional units in the *pks* gene cluster. Designed pNative_dCas9 strains transformed into *pks*^+^ NC101 *E. coli* significantly lower genotoxicity in HeLa cells after 4 h co-culture as compared to positive controls; x-axis labels show the *pks* island target of the sgRNA present. Genotoxic index is calculated by normalization against the median fluorescence index of HeLa cells co-cultured with Δ*pks* NC101 *E. coli* cells. MOI = 50, n = four technical repeats per independent biological replicate. N = three independent biological replicates. **d,** 2^-ΔΔCt^ qPCR analysis of *pks*^+^ NC101 *E. coli* transformed with pNative_dCas9 plasmids show highly specific targeting of the *pks* island. n = four technical repeats per independent biological replicate, N = two independent biological replicates. Data are means ± SD (**a, c, d**). Significance values calculated using one-way ANOVA with Dunnett’s multiple comparison test (**a, c**) and two-way ANOVA with Tukey’s multiple comparison test (**d**).

### dCas9 transcriptional silencing abolishes colibactin genotoxicity without resistance

Next, we implemented CRISPR interference (CRISPRi) using catalytically inactive Cas9 (dCas9) to silence transcription of colibactin biosynthetic genes in a sequence-specific manner. Unlike nuclease-active Cas9, dCas9 blocks transcription without inducing double-strand breaks or imposing bactericidal pressure, thereby minimizing selection for escape mutations. This strategy was motivated by our observation that killing bacteria with active Cas9 led to insertion-sequence–mediated *cas9* inactivation (**Figure 2b**). To confirm this, in the original pNative-Cas9, we introduced alanine (D10A) and alanine (H840A), two established mutations that abolish the DNA-cleaving activity of the Cas9 protein, rendering it catalytically inactive (**Extended Data** Fig. 1b). Upon transformation of different CRISPR-dCas9 constructs (sg*clbA*_287, sg*clbB*_4387, sg*clbB*_73 or sgRNA control) into *pks⁺* NC101, it was found that dCas9 did not inhibit the growth of *pks⁺* NC101 when comparing *pks*-specific sgRNAs to the control sgRNA (**Extended Data** Fig. 2c). This agrees with the loss of double-stranded DNA break-induced bacterial death for dCas9. Furthermore, by pooling multiple clones for DNA sequencing of different CRISPR-dCas9 modules, no insertional sequences were detected in NC101 transformed with these *pks*-specific sgRNAs paired with dCas9, indicating the genetic stability of CRISPR-dCas9 modules targeting the *pks* island (**Figure 2b**). As the *pks* island is dispensable for bacterial viability *in vitro* and *in vivo* ^19,24^, our findings indicate that dCas9 does not pose a negative selection against dCas9 as we have seen in the wild-type Cas9.

After validating the genetic stability of the CRISPR-dCas9 system in *pks⁺ E. coli,* we sought to identify lead sgRNA candidates for dCas9-mediated transcriptional silencing of colibactin biosynthetic genes. Based on prior work, effective dCas9-based repression requires guide RNAs targeting the non-template strand close to the 5’ end of the gene ^36^. Using an online design tool, we identified ∼20 computationally optimized sgRNAs targeting essential *pks* biosynthetic genes, including *clbB*, *clbC*, *clbH*, *clbI*, *clbJ*, *clbL,* and *clbN.* These genes were chosen because they were reported to be associated with separate transcriptional units ^37^. Each individual sgRNA along with dCas9 (pNative-dCas9) were cloned into the same low-copy plasmid with the SC101 origin. We then cocultured HeLa cells with *pks⁺* NC101 transformed with different CRISPR-dCas9 constructs. NC101 containing each of the 20 different *pks*-specific sgRNAs exhibited various degrees of reduction in genotoxicity despite no apparent differences in growth rates compared to wild-type NC101 (**Figure 2c**). Nonetheless, we focused on characterizing two sgRNAs, sg*clbB*_4387 and sg*clbC*_2313, given their complete loss of genotoxicity relative to uninfected HeLa cells.

To assess whether the reduction in genotoxicity resulted from the transcriptional silencing of target genes, we quantified gene expression of *clbB* and *clbC* in NC101 transformed with pNative-dCas9 constructs encoding sg*clbB*_4387, sg*clbC*_2313, or sgCtl. Quantitative PCR analysis of RNAs revealed that pNative-dCas9-sg*clbB*_4387 or pNative-dCas9-sg*clbC*_2313 resulted in 15-to 30-fold reductions in the RNA levels for *clbB* and *clbC*, respectively. In comparison, NC101 strains transformed with pNative-dCas9-sgCtl displayed minimal reduction in *clbB* or *clbC* relative to those of non-transformed bacteria (**Figure 2d**). Taken together, our results indicate that our CRISPRi-dCas9 system with optimized sgRNA design can effectively neutralize the genotoxicity of *pks⁺ E. coli* through transcriptional silencing of target gene expression.

**Figure 3.**
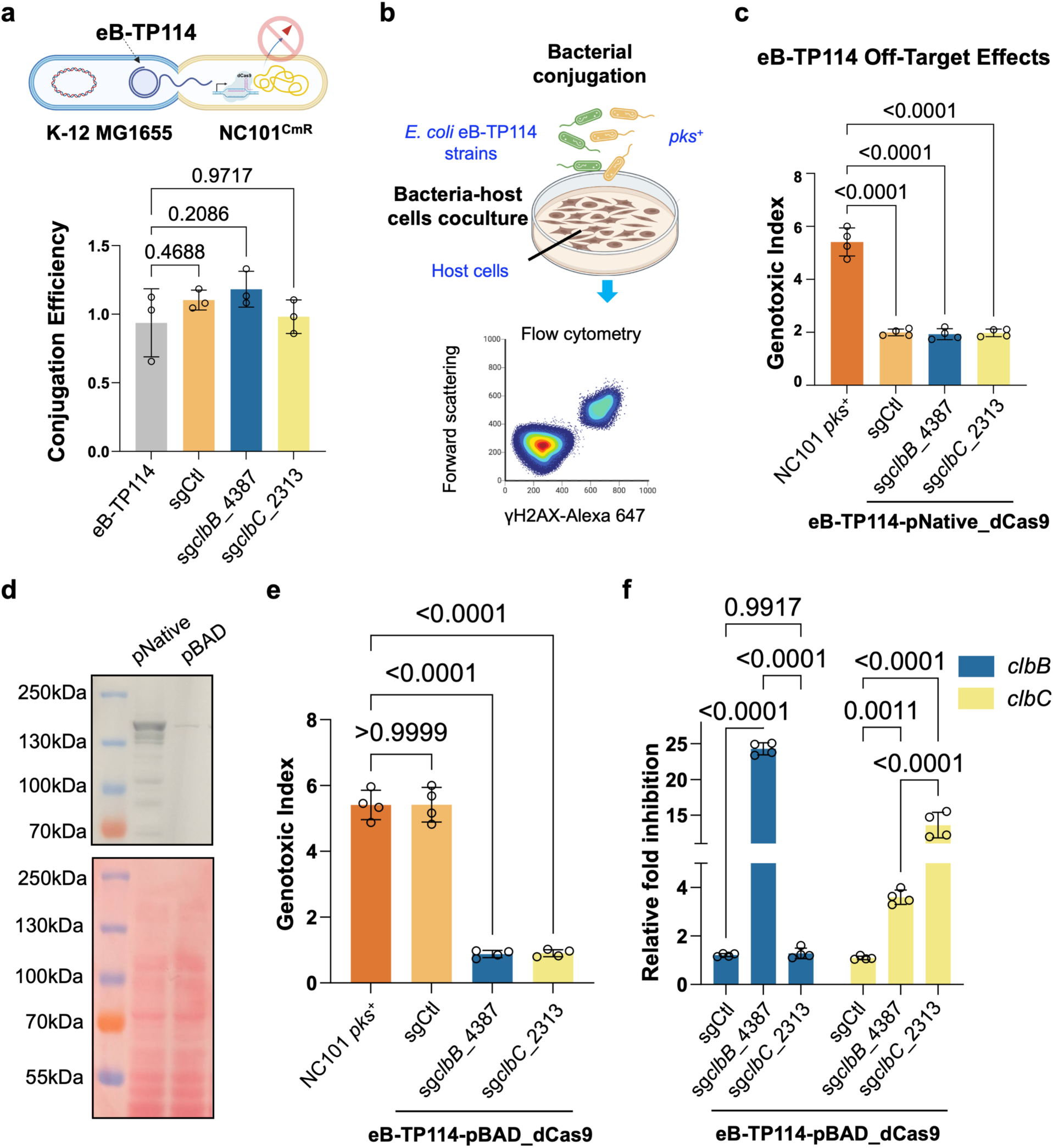
Conjugative plasmids efficiently deliver CRISPRi to silence colibactin. **a**, eB-TP114 encoding dCas9 and sgRNA displayed nearly 100% conjugation efficiency, and no decrease compared to eB-TP114 alone in *pks*^+^ NC101 *E. coli* carrying a chloramphenicol resistance marker (NC101^CmR^). n = three technical repeats per independent biological replicate, N = two independent biological replicates. Conjugation Efficiency = transconjugates/total recipients. **b**, Schematic showing the process of bacterial conjugation, co-culture, and flow cytometry of co-cultured host cells. **c**, The native dCas9 promoter, pNative, exhibited significant off-targeting when paired with a control sgRNA (sgCtl) in the eB-TP114 backbone. MOI = 100, n = four technical repeats per independent biological replicate, N = two independent biological replicates. **d**, Western Blot showing basal expression of dCas9 protein from the pBAD promoter was much lower than that of the native dCas9 promoter, pNative. Results are representative of two independent biological replicates. **e**, dCas9 protein driven by the pBAD promoter exhibited optimal on-target silencing of genotoxicity by *clb*-specific sgRNAs (sg*clbB*_4387 and sg*clbC*_2313) with minimal off-targeting effect from sgCtl. MOI = 100, n = four technical repeats per independent biological replicate, N = two independent biological replicates. **f**, 2^-ΔΔCt^ qPCR analysis of *pks*^+^ NC101 *E. coli* conjugated with eB-TP114-pBAD_dCas9 plasmids show highly specific targeting of the *pks* island, while also showing a downstream polar effect of sgclbB_4387 on the *clbC* gene. n = four technical repeats per independent biological replicate, N = two independent biological replicates. Data are means ± SD (**a, c, e, f**). Significance values calculated using one-way ANOVA with Dunnett’s multiple comparisons test (**a, c, e**), and two-way ANOVA with Tukey’s multiple comparison test (**f**).

### Conjugative plasmids efficiently deliver CRISPRi to silence colibactin

Having designed and characterized the CRISPR-dCas9 constructs via sgRNA screening in a replicative plasmid, we sought to deliver the system into *pks⁺ E. coli* in a more practical method. It is worth noting that the gut microbiota is a hotspot for horizontal gene transfer through bacterial conjugation. Conjugative plasmids can disseminate genes across phylogenetically diverse bacteria, including strains that harbor the *pks* island responsible for colibactin biosynthesis ^19^. We propose to co-opt the conjugation machinery for precision microbiome editing by CRISPRi. This strategy departs from conventional antibiotic or probiotic interventions toward programmable, self-propagating microbial therapeutics. We focus on TP114 as a self-transmissible broad-host-range conjugative plasmid for CRISPRi delivery for three reasons ^25^. First, TP114 achieves high conjugation rates in the mouse intestine, with ∼10-fold greater efficiency under anaerobic conditions that mimic the gut. Second, TP114 transfers across members in *Enterobacteriaceae*, including *Escherichia*, *Citrobacter*, and *Klebsiella*, taxa frequently linked to genotoxin-mediated pathology ^19,20^. Third, directed evolution has produced eB-TP114 with >400-fold higher conjugation efficiency ^25^. Thus, eB-TP114 was chosen in this study as a chassis to deliver the optimized dCas9 system to target the *pks* island. The pNative-dCas9 cassettes carrying sg*clbB*_4387, sg*clbC*_2313, or sgCtl in the original replicative plasmid were first transferred into eB-TP114 via two orthogonal DNA recombinases (FLP and Bxb1), denoted as eB-TP114-dCas9. Specifically, eB-TP114 was modified to include two specific FRT and Bxb1 sequences for FLP and Bxb1-mediated site-specific recombination ^25,38^. The FRT and Bxb1 sequences are present in both the replicative plasmid and eB-TP114, which allows for the transfer of a large DNA construct encoding pNative-dCas9 cassettes (**Extended Data** Fig. 1c). Subsequently, they were introduced into a domesticated nonpathogenic commensal *E. coli* K-12 MG1655 as the donor strains for proof-of-concept conjugation into *pks⁺ E. coli*. We quantified the conjugation efficiency between the K-12 donor with eB-TP114-dCas9 cassettes and a recipient strain, *pks⁺* NC101^CmR^, which encodes a genomically integrated chloramphenicol resistance gene. eB-TP114-dCas9 carrying different sgRNAs achieved ∼100% conjugation efficiency after mating for four hours, which is comparable to the empty eB-TP114 vector (**Figure 3a**). This suggests that the inclusion of the pNative-dCas9 did not interfere with the conjugation process.

We next assessed whether the dCas9 constructs in eB-TP114 remained able to neutralize genotoxicity upon conjugation to *pks⁺ E. coli.* It was noted earlier that dCas9 paired with the control sgRNA (sgCtl) in the original low-copy replicative plasmid exhibited a slight decrease in genotoxicity compared to wild-type NC101 (**Figure 2c**). However, the off-targeting effect was exacerbated when the pNative-dCas9-sgCtl was cloned into the eB-TP114 conjugative plasmid, likely due to the increased copy number of eB-TP114 compared to the low-copy replication origin SC101 (**Figure 2c and 3c**). Prior work by others showed that constitutive expression of dCas9 by its native promoter may contribute to off-targeting effects ^39–41^. To reduce the off-targeting effects while maintaining high specificity, we next employed an arabinose-inducible promoter (pBAD) to reduce the expression of dCas9 (pBAD-dCas9) (**Extended Data Fig. 1d-e**). Western blot analysis confirmed that the pBAD promoter yielded much lower dCas9 expression under the noninduced condition than that of the native promoter (**Figure 3d**). Consequently, NC101 transformed with pBAD-dCas9-sgCtl exhibited minimal off-targeting effects with the genotoxicity level comparable to wild-type NC101 (**Figure 3e**). Notably, despite much reduced expression of dCas9 driven by pBAD under the noninduced conditions (**Figure 3d**), the pBAD-dCas9-sg*clbB*_4387 or -sg*clbC*_2313 remained able to completely neutralize NC101-induced genotoxicity (**Figure 3e**).

The *pks* biosynthetic pathway is adapted to the anoxic intestinal lumen, and genes in the *pks* island are upregulated under anaerobic conditions ^42^, conferring higher genotoxicity than those under aerobic culture conditions. Therefore, it is critical to assess whether the eB-TP114-pBAD-dCas9 system can still effectively neutralize genotoxicity from *pks*^+^ bacteria pre-conditioned to the gut-relevant anaerobic conditions. Despite culturing *pks*^+^ bacteria carrying eB-TP114-pBAD-dCas9 constructs under anaerobic conditions, the genotoxic levels induced in HeLa cells were undetectable compared to the noninfected control (**Extended Data Fig. 4**). We further quantified gene expression of *clbB* and *clbC* in these transconjugates. qPCR analysis revealed that eB-TP114-pBAD-dCas9 carrying sg*clbB*_4387 or sg*clbC*_2313 resulted in 15-to 25-fold reductions in *clbB* and *clbC*, while the non-targeting control sgRNA did not reduce their expression compared to wild-type NC101 (**Figure 3f**). In addition to NC101, probiotics such as *Escherichia coli Nissle* 1917 (*EcN*), a phylogenetic lineage B2 strain, harbor the *pks* island encoding for colibactin biosynthesis ^43,44^. Although *EcN* is a popular chassis for live therapeutics, concerns have been raised about its potential to promote DNA mutations such as SBS88 in mammalian cells ^45^. It was found that the same eB-TP114-pBAD-dCas9 constructs (sg*clbB*_4387 or sg*clbC*_2313) resulted in 20-to 25-fold silencing in *clbB* and *clbC*, respectively, while the non-targeting sgRNA control did not (**Extended Data Fig. 5**).

**Figure 4.**
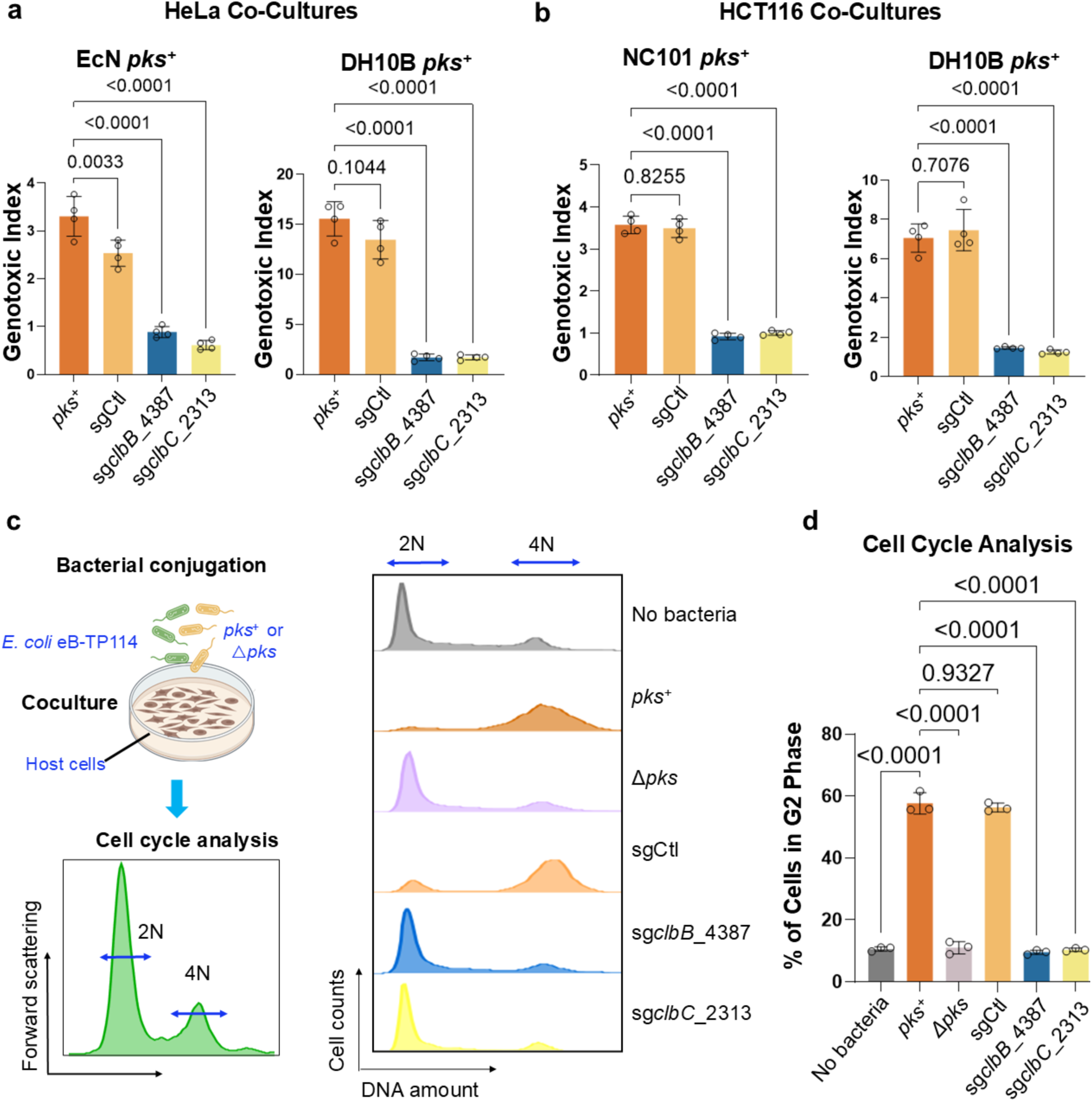
Validation of the conjugative CRISPRi module across different *pks*^+^ strains and cell lines in preventing the genotoxicity and cell cycle arrest in host cells. **a**, eB-TP114-pBAD_dCas9 strains transformed into different *pks*^+^ bacterial species significantly lower genotoxicity in HeLa cells after 4 h co-culture as compared to positive controls. MOI = 100 (*EcN*), MOI = 50 (DH10B), n = four technical repeats per independent biological replicate, N = two independent biological replicates. **b**, eB-TP114-pBAD_dCas9 strains transformed into different *pks*^+^ bacterial species significantly lower genotoxicity in HCT116 cells (a colon cancer cell line) after 4 h co-culture as compared to positive controls. MOI = 100 (NC101), MOI = 50 (DH10B), n = four technical repeats per independent biological replicate, N = two independent biological replicates. **c**, HeLa cells co-cultured for 4 h with eB-TP114-pBAD_dCas9 strains transformed into NC101 *pks*^+^ *E. coli* show a standard phenotype of 2N vs. 4N cells following 24 h incubation with antibiotics as compared to controls. MOI = 100, n = three technical repeats per independent biological replicate, N = two independent biological replicates; histograms are representative samples of the data presented in **d**. **d**, HeLa cells co-cultured for 4 h with eB-TP114-pBAD_dCas9 strains transformed into NC101 *pks*^+^ *E. coli* show a significantly lower percentage of cells in the G2 phase of the cell cycle following 24 h incubation with antibiotics as compared to controls. MOI = 100, n = three technical repeats per independent biological replicate, N = two independent biological replicates. Data are means ± SD (**a, b, d**). Significance values calculated using one-way ANOVA with Dunnett’s multiple comparisons test (**a, b**) and with Tukey’s multiple comparison test (**d**).

### Validation of the conjugative CRISPRi module across different *pks*^+^ strains and cell lines in preventing the genotoxicity and cell cycle arrest in host cells

To assess the generalizability of this strategy, we extended the eB-TP114-pBAD-dCas9 constructs to other *pks^+^* strains, including *EcN*, DH10B pBAC-*pks*, which harbors a bacterial artificial chromosome encoding the *pks* island ^8,10,46^, and three *E. coli* isolates from mice and guinea pigs. Following coculture with HeLa cells, the genotoxicity in different *pks⁺* strains was completely abolished by eB-TP114-pBAD-dCas9-sg*clbB*_4387 or sg*clbC*_2313 when compared to the non-targeting sgRNA control (**Figure 4a and Extended Data Fig. 6**). Most studies have utilized HeLa cells to assess genotoxicity induced by *pks*^+^ bacteria. To further evaluate our approach in the context of colon cancer, we demonstrated that pBAD-dCas9-mediated repression mitigated genotoxic effects in HCT116, a human colorectal cancer cell line (**Figure 4b**). To benchmark the effectiveness of our approach, we further compared it to D-serine, a small-molecule inhibitor previously shown to inhibit *clbB* transcription and thereby suppress colibactin *in vitro* ^47^. While D-serine modestly reduced DNA damage in NC101, it failed to fully protect host cells. By comparison, eB-TP114-pBAD-dCas9-sg*clbB*_4387 or sg*clbC*_2313, completely abolished genotoxicity in NC101 (**Extended Data** Fig. 7). These findings demonstrate that transcriptionally silencing the *pks* island can efficiently protect host cells from colibactin-mediated genotoxicity, outperforming the small molecule inhibitor, D-serine.

In addition to γH2AX staining, cells exposed to colibactin also experience cell-cycle arrest, which can be evaluated by staining nuclear DNA contents with nucleic acid-binding dyes for flow cytometry (**Figure 4c**). Consistent with previous findings ^10,48^, infection of HeLa cells with NC101 alone significantly reduced the percentage of cells characterized by 2N (diploid, G1 phase) DNA content compared to infection with the isogenic mutant NC101Δ*pks*, indicating that NC101 infection induced the cell cycle arrest. However, in NC101 transconjugates that carry eB-TP114-pBAD-dCas9-sg*clbB*_4387 or sg*clbC*_2313, the cell cycle profile was restored to levels seen in negative controls (NC101Δ*pks* or uninfected cells). In contrast, NC101 transconjugates with the non-targeting sgCtl construct failed to rescue this effect (**Figure 4d**). These data demonstrate that pBAD-dCas9 targeting the *pks* island effectively prevents both genotoxicity and cell cycle arrest induced by *pks*^+^ bacteria.

### CRISPR-dCas9 conjugation extends to silence another pathogenic metabolite, yersiniabactin

Having established *in vitro* proof-of-concept targeting colibactin, we next expanded our platform against another microbial metabolite implicated in gut inflammatory diseases.

Importantly, yersiniabactin (ybt), a siderophore also synthesized by *pks*^+^ bacteria, has been found to be enriched in inflammatory bowel disease patients ^21,49^. Interestingly, the biosynthetic gene cluster coding for ybt is in the vicinity of the *pks* island for colibactin. Both colibactin and ybt biosynthetic gene clusters are associated with the High-Pathogenic Island in *pks*^+^ bacteria ^50^, which has been speculated to aid in gut colonization of *pks*^+^ bacteria and promote inflammatory bowel disease and CRC ^51^. Using a similar methodology, we designed four sgRNAs targeting the 5’ end of the longest gene, *irp1*, in the *ybt* gene cluster and generated corresponding eB-TP114-pBAD-dCas9 constructs. After conjugation of eB-TP114-pBAD-dCas9 constructs via the K-12 donor strains into NC101^CmR^, we quantified transcript levels in different transconjugates. qPCR results indicated 5-to 25-fold reduction in the transcript levels of *irp1* mediated by different *irp1*-specific sgRNAs, while the non-targeting control sgRNA (sgCtl) showed no off-targeting effect (**Extended Data Fig. 8**). Unlike colibactin, ybt is relatively stable in the spent medium from bacterial culture. Therefore, we quantified the ybt abundance in bacterial culture by a quadrupole-time of flight-mass spectrometer. Consistent with the transcriptional silencing of *irp1*, the levels of ybt were undetectable in NC101 transconjugates carrying *irp1*-specific sgRNAs, comparable to those of the *irp1* knockout strain (**Extended Data Fig. 9**). Therefore, our results indicate that the same eB-TP114-pBAD-dCas9 system can be repurposed to target a different bacterial metabolite.

### Conjugative CRISPRi achieves *in vivo* conjugation and neutralizes the genotoxicity of *pks^+^ E. coli*

After validating our platform against colibactin and yersiniabactin *in vitro*, we prioritized colibactin for *in vivo* proof-of-concept testing, given its strong implication in the rising incidence of early-onset colorectal cancer ^6^. In contrast, yersiniabactin has primarily been linked to intestinal fibrosis and studied in germ-free, inflammation-prone mice. We therefore asked whether conjugative CRISPRi could transfer into *pks*⁺ *E. coli in vivo* and suppress their genotoxicity in mice under conventional housing. Mice were pretreated with 2 g/L streptomycin in drinking water for three days (**Figure 5a**). Mice were then gavaged twice weekly with NC101 for three weeks to establish chronic infection. In parallel, animals received twice-weekly oral gavage with one of the following *E. coli* donor strains: (1) *Ec^dCas9-sgCtl^*, where the sgRNA is the scramble DNA control, (2) *Ec^dCas9-sgclbB_4387^* targeting *clbB*, (3) *Ec^dCas9-sgclbC_2313^*targeting *clbC*, or (4) PBS vehicle. Across treatment groups, body weights remained stable (**Figure 5b**), cytokine and chemokine profiles (∼30 analytes, Luminex) showed no significant differences (**Figure 5c**), and histology of the colons, spleens, kidneys, and livers revealed no overt toxicity relative to PBS controls (**Extended Data Fig. 10**). These organs were specifically chosen because (1) the colon is the primary site of colibactin production and tumorigenesis, (2) the liver and spleen are sentinel organs for systemic bacterial dissemination and immune activation, and (3) the kidneys are highly sensitive to off-target toxicities and serve as a standard readout for systemic safety in therapeutic development.

To assess *in vivo* conjugation, fecal *E. coli* was recovered on MacConkey agar with and without antibiotic selection on days 2 and 16, post-NC101 gavage. Enumeration of donor, recipient, and transconjugate strains revealed that there was 50-80% conjugation efficiency *in vivo* from *Ec^dCas9-sgCtl^* and *Ec^dCas9-sgclbB_4387^*and slightly lower conjugation efficiency from *Ec^dCas9-sgclbC_2313^*(**Figure 5d**). We next examined whether *in vivo* transconjugates had lost genotoxicity. After recovery of *E. coli* from mouse feces on the MacConkey agar plates, *E. coli* transconjugates were confirmed by PCR for both the *clbA* gene (NC101-specific) and *dcas9* (conjugative plasmid-specific). When cocultured with HeLa cells, transconjugates derived from *Ec^dCas9sg-clbB_4387^*or *Ec^dCas9-sgclbC_2313^* were completely non-genotoxic, whereas those carrying the non-targeting sgRNA retained genotoxicity (**Figure 5e**).

To define transcriptomic consequences of *in vivo* silencing, we performed RNA sequencing on fecal transconjugates recovered from mice receiving *Ec^dCas9-sgCtl^*, *Ec^dCas9-sgclbB_4387^*, or *Ec^dCas9-sgclbC_2313^* (three biological replicates per group). Differentially expressed genes (DEGs) are summarized in volcano plots (**Extended Data Fig. 11a, b**), which highlight changes within the *pks* operon and a few alterations outside of it. Notably, dCas9-sg*clbB*_4387 produced an apparent down-regulation of *clbC*, and dCas9-sg*clbC*_2313 showed the strongest reduction in *clbD* (**Extended Data Fig. 11a, b**). This pattern is consistent with a downstream polar effect within the operon and with the physical location of our guides, where both sg*clbB*_4387 and sg*clbC*_2313 bind ∼2–4 kb downstream of the 5′ ends of *clbB* and *clbC*, respectively. Only RNA fragments downstream of the dCas9 binding site are expected to be downregulated, whereas upstream fragments should be unaffected by dCas9 as previously reported ^36^. As a result, the standard RNA sequencing from the Illumina paired-end 150-bp reads may underestimate silencing at the targeted gene and therefore shift the maximum effect to the downstream gene due to the polar effect of the colibactin biosynthetic genes (i.e., *clbC* for sg*clbB*_4387 and clbD for sg*clbC*_2313). Consistent with the mapping results, by designing qPCR primers to target transcripts downstream of each sgRNA binding site, we confirmed knockdown of the targeted genes in transconjugates (**Figure 5f**). Additionally, principal component analysis indicated minimal global perturbation, with sg*clbB*_4387 clustering closely with sgCtl (**Extended Data Fig. 11c**). These trends agree with the volcano plots that relative to dCas9-sgCtl, dCas9-sg*clbB*_4387 exhibits markedly fewer differentially expressed genes than that of dCas9-*clbC*_2313 (**Extended Data Fig. 11a, b**). Furthermore, Kyoto Encyclopedia of Genes and Genomes (KEGG) analysis of DEGs identifies ‘biosynthesis of secondary metabolites’ as the most affected pathway, likely attributed to the downregulation of the *pks* island (**Extended Data Fig. 11d**). Together, these data indicate that the conjugative CRISPRi system efficiently transfers *in vivo*, stably silences expression of *pks* genes in *pks^+^ E. coli,* and neutralizes colibactin genotoxicity without systemic toxicity in mice.

**Figure 5.**
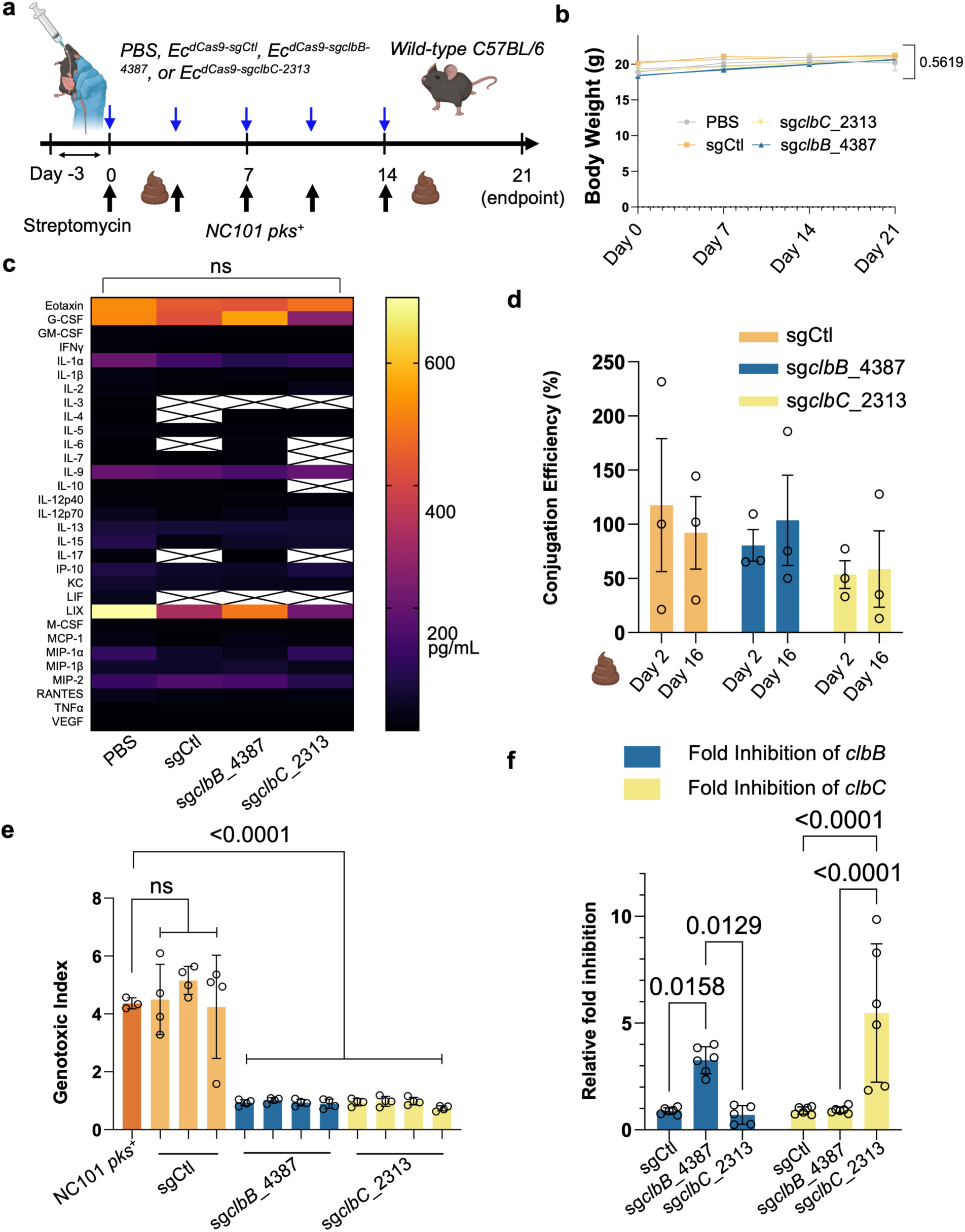
*In vivo* safety and efficacy of the conjugative CRISPRi in wild-type mice. **a**, Experimental method for assessing the safety and efficacy of *Ec^dCas9^*strains *in vivo*. **b**, Body weight records show no statistically significant difference in slope of values over time between experimental groups. n = four or five mice per group. **c**, Luminex analysis of blood samples collected at endpoint shows no significant difference in blood analytes between the treatment groups. White boxes with crosses indicate that values were below the threshold of detection. n = four or five mice per group. **d**, Conjugation efficiency analysis between week one and week three show values between 50% and 100% efficiency for the three donor strain groups. Conjugation Efficiency = transconjugates/total recipients. **e**, *In vivo* transconjugates isolated from fecal samples show significantly lower genotoxicity in HeLa cells after 4 h co-culture as compared to positive controls and sgCtl *in vivo* transconjugates, further validating results from the earlier *in vitro* transconjugates. MOI = 100, n = four technical repeats per independent biological replicate, N = two independent biological replicates. **f**, 2^-ΔΔCt^ qPCR analysis of *in vivo* transconjugates show highly specific targeting of the *pks* island. n = six technical repeats per independent biological replicate, N = two independent biological replicates. Data are means ± SEM (**b, d**), means ± SD (**e, f**), and median values (**c**). Significance values calculated using simple linear regression analysis (**b**), two-way ANOVA with Dunnett’s multiple comparisons test (**c**), one-way ANOVA with Dunnett’s multiple comparisons test (**e**) and two-way ANOVA with Tukey’s multiple comparisons test (**f**).

### Conjugative CRISPRi suppresses tumorigenesis and reduces NC101 colonization in a colitis-associated mouse colorectal cancer model

After validating the efficacy of the conjugative CRISPRi in wild-type mice colonized with *pks*^+^ *E. coli*, we next tested whether this approach could mitigate tumorigenesis induced by *pks*^+^ *E. coli* in *Apc*^Min/+^ mice. Despite certain limitations, this mouse model has been frequently used to study the effects of colibactin due to its ability to accelerate colonic tumor formation when treated with dextran sodium sulfate (DSS) ^26,52,53^. Following DSS pre-treatment, *Apc*^Min/+^ mice were gavaged weekly with NC101 for three weeks to establish chronic infection. Concurrently, mice received one of three donor strains by oral gavage: (1) *Ec^eB-TP114^* (eB-TP114 lacking dCas9 and sgRNA), (2) *Ec^dCas9-sgCtl^*, and 3) an equal mixture of Ec^dCas9-sgclbB_4387^ and Ec^dCas9-sgclbC_2313^, denoted as ‘*Ec^dCas9-clbB/C^*’) (**Extended Data Fig. 12a**). Donors were given four hours after NC101 to enable *in situ* conjugation in the mouse gut and were administered twice weekly for a total of six doses. Twenty-eight days after the first NC101 dose, mice were euthanized for tumor burden analysis. Notably, mice treated with the *Ec^dCas9-clbB/C^* cocktail developed significantly fewer tumors than the sgRNA control group (*Ec^dCas9-sgCtl^*), with one mouse remaining completely tumor-free (**Extended Data** Fig. 12b, d). Additionally, no significant body weight changes were observed over the course of different treatment regimens (**Extended Data** Fig. 12c).

After validating the cocktail approach (*Ec^dCas9-clbB/C^*), we next assessed single *pks*-specific sgRNAs in the DSS/Apc^Min/+^ model (**Figure 6a**). Our earlier qPCR analysis showed that eB-TP114-pBAD-dCas9-sg*clbB*_4387 suppressed expression of both *clbB* and *clbC*, with a stronger effect on *clbB*, whereas sg*clbC_2313* selectively reduced *clbC* alone (**Figure 3f**). Given that *clbC* lies downstream of *clbB* in the *pks* island, the dCas9-sg*clbB*_4387 construct may exert a polar effect that partially silences *clbC*. Therefore, we focused on comparing the two arms, *Ec^dCas9-sgCtl^* and *Ec^dCas9-clbB^*. Using the same treatment schedule, *Ec^dCas9-sgclbB_4387^* markedly lowered colonic tumor burden compared to *Ec^dCas9-sgCtl^*, with results pooled from two independent biological replicates (n=5–7 mice per replicate, mixed sex) (**Figure 6b**). No significant body weight changes were observed during treatment (**Extended Data Fig. 13**). We further selected D-serine as a pharmacologic inhibitor because prior work reports that it suppresses *clbB* transcription and attenuates colibactin-associated genotoxicity ^37^. It is orally deliverable, widely regarded as safe, and has shown benefit in other mouse disease models ^47,54,55^. Therefore, D-serine may offer a fair, target-matched benchmark against our genetic silencing strategy. Continuous D-serine administration (600 mg/L in drinking water), a dose reported in prior studies ^47,54,55^, failed to lower tumor counts in DSS/Apc^Min/+^ mice (**Extended Data Fig. 14**). Therefore, although both small-molecule- and CRISPRi-based interventions converge on *clbB* inhibition, engineered bacteria delivering eB-TP114-pBAD-dCas9 achieved robust and durable tumor suppression where a well-tolerated small molecule did not, underscoring the better *in vivo* efficacy of our conjugative CRISPRi strategy.

Because the abundance of pro-carcinogenic NC101 correlates with tumor burden in DSS/*Apc*^Min/+^ mice ^26^, we also evaluated whether the conjugative CRISPRi system reduced NC101 colonization. To this end, we analyzed the abundance of NC101 longitudinally in three different treatment groups receiving (1) NC101 and PBS, (2) NC101 and *Ec^dCas9-sgCtl^* and (3) NC101 and *Ec^dCas9-sgclbB_4387^*in DSS/*Apc*^Min/+^ mice following the same treatment scheduling in **Figure 6a**. The NC101 abundance was quantified using qPCR by normalizing the copy number of the *clbP* gene to the total bacterial load via 16S primers to derive the delta cycle threshold (Ct) values (**Figure 6c**). The relative abundance of NC101 in the treatment group receiving NC101 and *Ec^dCas9-sgclbB_4387^* decreased by approximately 1000 times compared to the treatment group receiving NC101 and *Ec^dCas9-sgCtl^* at the midpoint (on Day 23), where the Ct values (*clbP* relative to 16S) inversely correlated with the NC101 abundance. Additionally, the non-targeting sgCtl group (*Ec^dCas9-sgCtl^*) showed colonization levels indistinguishable from PBS controls, indicating that the reduction stemmed from targeted CRISPRi rather than nonspecific donor-recipient competition (**Figure 6c**). Notably, this trend still persisted at the endpoint (day 34) when PBS and sgCtl groups required euthanasia due to morbidity, albeit with reduced fold differences (∼30 times) between *Ec^dCas9-sgCtl^* and *Ec^dCas9-sgclbB_4387^*. Colibactin produced by *pks*^+^ bacteria has been shown to suppress the abundance of other gut microbiota members in neonatal mice ^56^, suggesting that colibactin may confer competitive ecological benefits to *pks*^+^ bacteria. Therefore, the markedly reduced abundance of NC101 from *Ec^dCas9-sgclbB_4387^*relative to that of *Ec^dCas9-sgCtl^* may be attributed to the loss of the competitive niche benefit conferred by colibactin. By repressing *clbB*, *Ec^dCas9-sgclbB_4387^* removes a genotoxin-mediated competitive advantage, which may expose NC101 to greater interspecies competition. However, it is also possible that NC101 transconjugates from *Ec^dCas9-sgclbB_4387^* displayed decreased proliferation relative to that of *Ec^dCas9-sgCtl^* despite that the transcriptomics showed no major global RNA expression changes (**Extended Data** Fig. 11). To rule out this latter possibility, we isolated *in vivo* transconjugates and confirmed that NC101 containing dCas9-sgCtl or dCas9-sg*clbB_*4387 proliferated comparably to wild-type NC101, arguing against a direct proliferation defect as the primary driver (**Extended Data** Fig. 15). Together with the observed reduction in colibactin-mediated genotoxicity, our findings indicate that conjugative CRISPRi reduces the ecological persistence of *pks*⁺ NC101, likely by removing a colibactin-dependent niche advantage in *Apc*^Min/+^ mice.

To further evaluate the impact of the conjugative CRISPRi on gut microbiota composition, we performed 16S rRNA gene sequencing on fecal samples from DSS/*Apc*^Min/+^ mice infected with *pks*⁺ NC101. Mice were randomly assigned by sex and age before gavaging with bacteria. Three treatment groups were performed in NC101-colonized mice that received (1) NC101 and PBS vehicle, (2) NC101 and *Ec^dCas9-sgCtl,^* or (3) NC101 and *Ec^dCas9-sgclbB_4387^*. Shannon and Chao1 indices of alpha diversity showed an increase in community evenness and richness in the treatment group *Ec^dCas9-sgclbB_4387^* in comparison to the vehicle and *Ec^dCas9-sgCtl^* control groups (**Figure 6d**). Beta diversity analysis revealed that administration of *Ec^dCas9-sgclbB_4387^* is capable of altering community structure. Permanova analysis and pairwise Adonis testing of unweighted unifrac distances revealed a shift in community compositions at both midpoint (*p* = 0.005) and endpoint (*p* = 0.004) between mice receiving *Ec^dCas9-sgclbB_4387^* and both control groups (**Figure 6e**). While the PBS vehicle and *Ec^dCas9-sgCtl^* controls clustered together, *Ec^dCas9-sgclbB_4387^*treatment led to a separate clustering with statistically significant differences (**Figure 6e**). Consistent with this shift, we observed a progressive decrease in the relative abundance of *Enterobacteriaceae*, a family often expanded during intestinal inflammation and linked to adverse clinical outcomes ^20^. Both *Enterobacteriaceae* and the genus *Escherichia* displayed reduced relative abundance in *pks*⁺ NC101-infected mice co-treated with *Ec^dCas9-sgclbB_4387^* (**Figure 6f**, **Extended Data Fig. 16**). The 16S sequencing, coupled with the qPCR quantitation of NC101, suggests that *Ec^dCas9-sgclbB_4387^* may attenuate the expansion of inflammation-associated *Enterobacteriaceae*.

**Figure 6.**
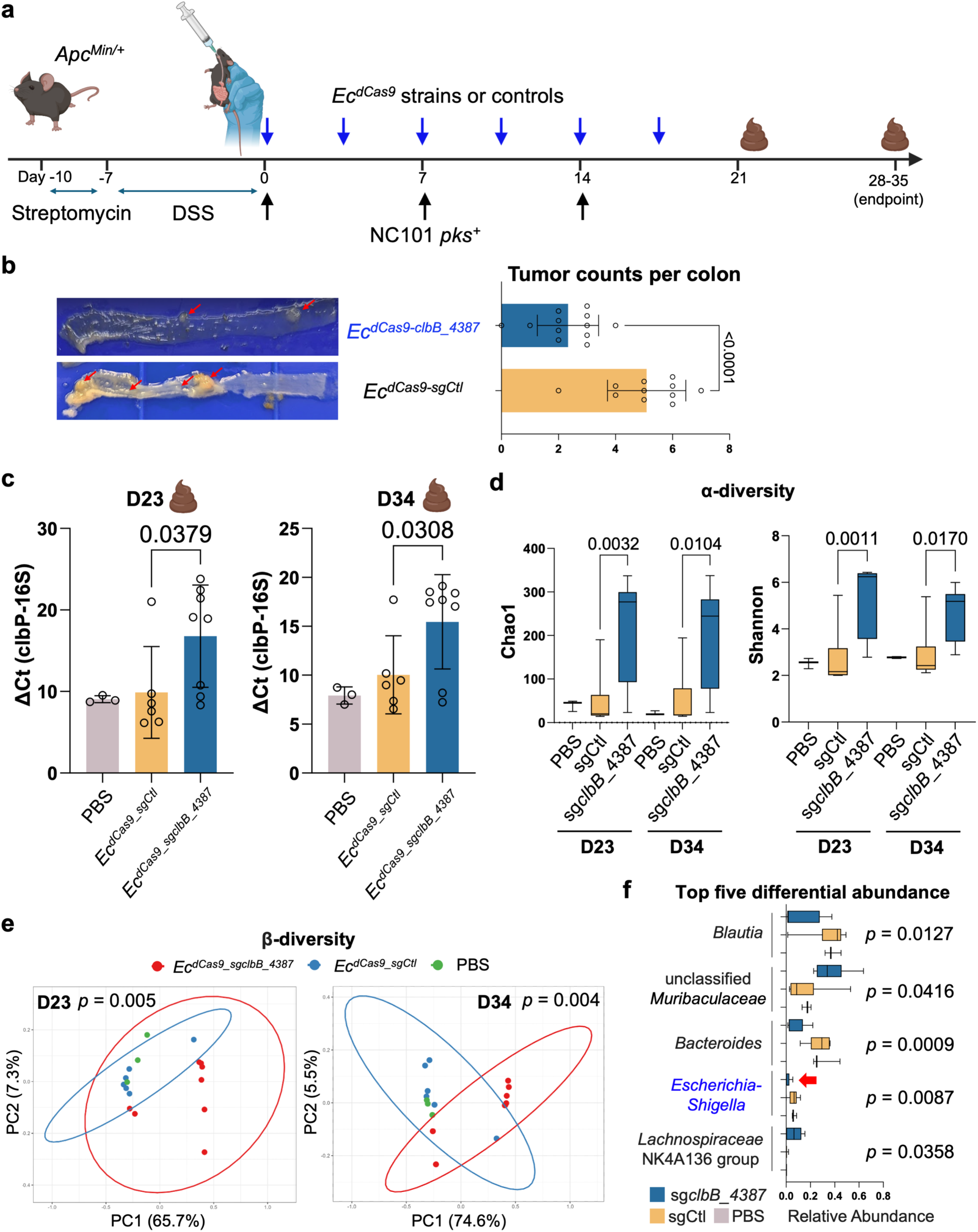
Conjugative CRISPRi reduces *pks*⁺ colonization and tumor burden while preserving commensal diversity. **a**, Experimental method for assessing the efficacy of *Ec^dCas9^* strains *in vivo*. **b**, Data pooled from two independent experiments shows a statistically significant decrease in tumor burden between *Ec^dCas9-sgclbB_4387^* and *Ec^dCas9-sgCtl^* donor strains. n=5–7 mice per replicate with mixed male and female Apc^Min/+^ mice per group. **c**, ΔCt qPCR analysis of fecal gDNA between the *clbP* gene and the 16S bacterial endogenous control shows a statistically significant decrease for the *Ec^dCas9-sgCtl^* treatment group, indicating an increase in the presence of *pks*-associated genes as compared to the *Ec^dCas9-sgclbB_4387^* treatment group (a higher ΔCt value indicates a lower NC101 abundance). **d**, Alpha diversity indices (Chao1 index and Shannon index) at days 23 and 34. In box plots, the bounds of the box indicate the 25th–75th percentiles, the center line indicates the median, and whiskers extend from the minimum to the maximum values. **e**, Principal coordinates analysis (PCoA) plots based on unweighted UniFrac distance showing distinct clustering of fecal microbiota composition among NC101 + PBS, NC101 + *Ec^dCas9-sgCtl^*, and NC101 + *Ec^dCas9-sgclbB_4387^* groups at days 23 and 34. Percent variation explained by each axis is indicated. 95% confidence ellipses are shown for *Ec^dCas9-sgCtl^* and *Ec^dCas9-sgclbB_4387^*, and the *p*-values shown compare these two groups. **f**, Top five differential abundance (%) at the genus level from fecal microbiota in different groups at Day 23. In box plots, the bounds of the box indicate the 25th–75th percentiles, the center line indicates the median, and whiskers extend from the minimum to the maximum values. Red arrow indicates significant reduction of *Escherichia-Shigella* by *Ec^dCas9-sgclbB_4387^* relative to PBS and *Ec^dCas9-sgCtl^* control groups. Data are means ± SD (b, c). Significance values calculated using Welch’s two-sided unpaired t-test (b), one-way ANOVA with Šídák’s multiple comparisons test (c, d), Adonis PERMANOVA with pairwise Adonis testing (e), and one-way ANOVA (f).

## Discussion

The rising incidence of early-onset colorectal cancer has been linked to pathogenic gut bacteria that produce procarcinogenic metabolites, with colibactin representing one of the most intensively studied. Recent evidence indicates that *pks*⁺ bacteria can induce the earliest APC driver mutations during childhood, well before clinical detection of CRC, underscoring a critical early-life window where microbial interventions may provide lasting preventive benefit. Despite this causal link between colibactin and mutagenesis, few therapeutic strategies have been designed to directly suppress colibactin production *in situ*. Here, we present conjugative CRISPRi as a broadly applicable strategy to silence colibactin biosynthesis in the gut. Unlike small molecular inhibitors, CRISPRi suppresses colibactin at its genetic source. By leveraging a self-transmissible plasmid chassis, our approach achieves three key outcomes: (1) efficient horizontal transfer of dCas9-based silencing modules across diverse *pks*⁺ *E. coli*, (2) durable repression of *clb* genes without bacterial killing or mutational escape, and (3) robust suppression of DNA damage, NC101 colonization, and tumor burden in mice. Together, these features establish conjugative CRISPRi as a first-in-class approach to genetically reprogram disease-associated microbes directly in the gut.

From a fundamental perspective, this work expands the microbiome engineering toolkit beyond bacteriophage delivery and synthetic probiotics. Conjugative CRISPRi exploits the evolutionary trait of horizontal gene transfer to disseminate programmable repression modules across complex microbial communities. In addition to neutralizing genotoxicity, silencing *clbB* removed a colibactin-mediated interference-competition advantage, reducing the ecological persistence of *pks*⁺ strains without imposing an intrinsic growth defect on NC101. This competition-driven loss of fitness suggests that programmable repression of the *pks* island may reshape community structure by diminishing colibactin-mediated antagonism, an effect likely to generalize to other pathogenic effectors and siderophores. Beyond colibactin, we also validated the flexibility of our chassis by targeting yersiniabactin, demonstrating the versatility of conjugative CRISPRi for silencing distinct pathogenic metabolites encoded by conserved biosynthetic islands. From a translational perspective, this study demonstrates that conjugative CRISPRi outperforms a well-tolerated small molecule, D-serine, in reducing tumorigenesis. Although both target *clbB*, only CRISPRi achieved durable silencing and tumor suppression *in vivo*, highlighting the limitations of metabolic inhibitors compared to programmable genetic repression. This is likely because the CRISPRi approach not only silences the genotoxicity of *pks*⁺ bacteria, but also abolishes colibactin-mediated antagonism. Although we deploy conjugative plasmids as the delivery vehicle here, we reason that the CRISPRi cassette is inherently portable and could be adapted to alternative platforms such as bacteriophages. We prioritized a self-transmissible conjugative plasmid for in-gut delivery because of its broad host range across *Enterobacteriaceae* (e.g., *Escherichia*, *Citrobacter*, *Klebsiella*), taxa frequently implicated in genotoxin-mediated pathology. In contrast, many phages exhibit narrow host specificity that can limit dissemination without extensive engineering approaches to optimize the target specificity ^17^. Moreover, we found that CRISPR-dCas9 achieved durable, on-target silencing of biosynthetic genes for colibactin and effectively abolished genotoxicity, while avoiding the mutational escape that undermined bactericidal wild-type Cas9 approaches. This combination, promiscuous DNA transfer with guide-directed transcriptional repression, enables dissemination to diverse *pks*⁺ strains while maintaining sequence-level specificity via the sgRNA–dCas9 module.

Nonetheless, several questions remain. While our assays focused on γH2AX staining, transcript silencing, and tumor burden, future studies should incorporate whole-genome sequencing of tumors or organoids to assess whether conjugative CRISPRi reduces colibactin-specific mutational signatures. Moreover, the ecological consequences of CRISPRi-mediated repression of *pks*⁺ bacteria within polymicrobial communities remain incompletely understood. Germ-free or synthetic community models may help dissect how CRISPRi alters competition between *pks*⁺ bacteria and commensals, and whether horizontal transfer propagates silencing across broader taxonomic niches.

Looking forward, conjugative CRISPRi represents a next-generation live biotherapeutic paradigm enabled by synthetic biology. Unlike conventional probiotics that rely on metabolic competition or small-molecule secretion, our system programs commensal bacteria to deliver precision genetic silencing directly to pathogenic strains in situ. Such a strategy could be extended beyond colibactin to a wide range of microbial metabolites implicated in cancer, inflammatory bowel disease, and neuroimmune disorders. Importantly, the modularity of dCas9 and guide RNAs allows for rapid reprogramming against new microbial targets, positioning conjugative CRISPRi as a versatile chassis for therapeutic microbiome engineering. As synthetic biology moves toward programmable, self-propagating interventions, our study provides a translational framework for building live biotherapeutics that move beyond correlation into causal modulation of disease-driving microbial functions.

## Acknowledgments

This work was supported by CDMRP PRCRP Idea Award (W81XWH-21-1-0324) (JL), Swim Across America (SAA) Young Investigator Award (JL), the NIH Office of the Director (1DP2GM154019-01) (JL), and National Cancer Institute (R01CA299955 and R01CA303150) (JL). We want to thank Dr. Christian Jobin for providing NC101 and NC101Δ*pks* strains, and Drs. Jean-Philippe Nougayrède and Eric Oswald for sending DH10B pBAC and DH10B pBAC-*pks* strains. We would like to thank the ULAM Pathology Core and the Rogel Cancer Center at the University of Michigan, Ann Arbor. We would like to express our gratitude to other members of the Li lab, Drs. James Fox and Zhongming Ge from MIT, and Dr. Yingzhong Li from SunVax mRNA Therapeutics, Inc., for helpful and substantial suggestions.

## Author contributions

J.L. and B.H. conceptualized the study. J.L. and B.H. wrote manuscripts. B.H., H.T., C.F., A.H., M.J., C.C., Z.W., and N.H. designed the experiments, analysis, and data interpretation. B.H., H.T., A.H., M.J., C.C., Z.W., C.F., Z.D., and S.M. performed the experiments. S.K., Y.S. and O.Y. guided during the optimization and assisted during the experimental procedures and analysis. All the authors contributed to data interpretation and the final version of the manuscript.

## Competing interests

J.L. received sponsored research agreements from Eco Animal Health, Ningbo Menovo Pharmaceutical, and Qingdao Saiding. The University of Michigan has filed a provisional patent related to this work. The remaining authors declare no competing interests.

Correspondence and requests for materials should be addressed to Jiahe Li, PhD.

## Materials and Methods

### Chemicals

All chemicals and cell culture broth were purchased from Fisher Scientific International Inc. (Cambridge, MA, USA) unless otherwise noted, and were of the highest purity or analytical grade commercially available. DNA and RNA oligos were ordered from Sigma Aldrich (St. Louis, MO, USA) and Integrated DNA Technologies (Coralville, IA, USA). All molecular cloning reagents, including restriction enzymes, competent cells, and the Gibson assembly kit, were purchased from New England Biolabs (Ipswich, MA, USA).

### Bacterial strains and growth conditions

Unless specified elsewhere for cocultures, all bacterial strains listed in **Table S2** were routinely maintained in lysogeny broth (LB) media and incubated at 37 °C under aerobic conditions. NC101 and NC101Δpks strains were provided by Dr. Christian Jobin. DH10B pBAC and DH10B pBAC-pks strains were a gift from Drs. Jean-Philippe Nougayrède and Eric Oswald. E. coli Nissle was a gift from Rudolf von Bünau. MIT A2 and MIT A21 were mouse pks^+^ isolates from Dr. James Fox’s lab. Kanamycin (50 μg/ml) and chloramphenicol (35 μg/ml) were included in LB media for strains with corresponding resistance genes in the plasmid. Conjugative plasmids, including eB-TP114, eB-TP114::tetB (eB-TP114Δaph-III::tetB), pKill1, pE-FLP, and pBXB1, were provided by Sébastien Rodrigue (Université de Sherbrooke). All studies using these constructs adhered to the material transfer restrictions stipulated by the providers.

### Construction of the CRISPR-Cas9 and CRISPR-dCas9 modules

The pKill1-Cas9 vectors were constructed by NEBuilder HiFi DNA Assembly (New England Biolabs) using pKill1 (a gift from Sébastien Rodrigue) as the backbone and replacing the 20-nt chloramphenicol (cat) spacer within the sgRNA cassette with target-specific spacers (primer sequences in **Table S4**). sgRNA sequences were designed using the CRISPR gRNA Design tool from ATUM (https://www.atum.bio/eCommerce/cas9/input) and sgRNAs targeting the non-template coding region with high scoring were selected for cloning. For constitutive expression, Cas9 was maintained under its native promoter on pKill1; for arabinose-inducible expression, primers were used to amplify the pBAD promoter and replace the native constitutive promoter sequence (“pNative”) with pBAD, using the pNuc-cis_TevSpCas9 template (a gift from David Edgell) as the promoter source. All assemblies were transformed into E. coli NEB5a and verified by whole-plasmid nanopore sequencing prior to use.

### Construction of CRISPR-dCas9 into eB-TP114

CRISPR–Cas9 modules were integrated into the eB-TP114 conjugative plasmid using the previously described double recombinase-operated insertion of DNA (DROID) workflow ^38^, which employs Bxb1 integrase–mediated recombination between attB/attP sites followed by Flp/FRT excision to remove auxiliary sequences and stabilize the insert.

Briefly, a TP114 loading dock containing attB/bxb1 and an FRT site was used to accept the CRISPR–Cas9 payload carrying the cognate attP^bxb1 and an FRT site; after Bxb1-mediated integration, Flp recombinase resolved the flanking FRT sites to yield the final construct. All resulting TP114::CRISPR–Cas9 plasmids were confirmed by whole-plasmid sequencing.

### Conjugation

Donor and recipient strains were inoculated in 2 ml LB and placed in a shaker (30 or 37°C, 250rpm) overnight. The next day, 10-fold diluted overnight bacterial cultures continued to expand in fresh media at 30 or 37 °C, 250rpm for about 1.5 h until bacteria reached the exponential phase (1 volume of donor for every recipient strain being used). Bacterial cultures were spun down and washed twice in sterile PBS and then resuspended in sterile LB without antibiotics. An equal volume of donor and recipient bacteria was mixed, and half of the volume was added as a mating spot on antibiotic-free LB agar plates. After the spot dried, plates were incubated for at least 4 h at 30 or 37 °C and then bacteria were tri-streaked onto LB agar plates with the necessary antibiotic resistances (donor-only and recipient-only plates were also streaked to confirm transconjugates were the only growth).

### Conjugation efficiency

Conjugation frequencies were determined following the previous protocol ^25^. Briefly, overnight cultures of the donor (K-12 MG1655^NxR^ carrying eB-TP114 with the indicated CRISPR–dCas9 module and a kanamycin resistance cassette) and the recipient (pks⁺ NC101^CmR^ with a chromosomal chloramphenicol-resistance cassette) were adjusted to OD₆₀₀ ≈ 1.0, pelleted (12,000×g, 1 min), washed once in sterile LB, and resuspended for mating. Donor and recipient were mixed 1:1 and incubated at 37 °C for 2 h either on LB agar or in LB broth (on a rotating mixer at 20 rpm). Mating mixtures were resuspended in PBS, serially diluted 1:10, and 5 µl spots were plated in duplicate on selective media to enumerate donors, recipients, and transconjugates. Conjugation efficiency was calculated as transconjugates per total recipients (Conjugation Efficiency = transconjugates (Cm and Kan) /total recipients (Cm)). All assays were performed with ≥ three biological replicates.

### Cell culture

HeLa (CRM-CCL-2™) and HCT116 (CCL-247™) were purchased from American Type Culture Collection (ATCC; Rockville, MD). HeLa and HCT116 were maintained in complete Dulbecco’s modified Eagle’s medium (DMEM; Corning, NY) supplemented with 10% fetal bovine serum (FBS; Corning, NY) and 100 U/ml penicillin-streptomycin (P/S; Corning, NY). HeLa and HCT116 cell lines were maintained at 37 °C in a humidified incubator with 5% carbon dioxide (CO_2_). Cells at passages 5-20 were used for the experiments.

### Western Blot

Bacteria strains were inoculated in 2 ml LB and placed in a shaker (30 or 37 °C, 250rpm) overnight. The next day, 10-fold diluted overnight bacterial cultures continued to expand in fresh media at 30 or 37 °C, 250rpm for about 2 h until bacteria reached the exponential phase. 200 μL aliquot of bacterial culture was pelleted by centrifugation and lysed by 100 μL 1×SDS loading buffer followed by boiling for 10 min. The samples were first separated with 8% SDS-PAGE gel and then transferred to a nitrocellulose membrane (Fisher Scientific). The membranes were incubated with the anti-Cas9 (clone 7A9, Biolegend, San Diego, CA, catalog no. 844301637301) with 1:2000 dilution overnight in a cold room, and the secondary antibody anti-mouse IgG HRP (Invitrogen, Waltham, MA, catalog no. 626520) at room temperature for 1 h. Thermo Scientific™ Pierce™ CN/DAB Substrate Kit (colorimetric Western) Premixed Pierce 3,3′-diaminobenzidine substrate (Fisher Scientific) was used to detect the target proteins.

### In vitro infection assay

NC101, NC101 Δpks, and NC101 transformed or conjugated with the dCas9 module were inoculated in 2 ml LB and placed in a shaker (30 or 37 °C, 250rpm) overnight. The next day, 10-fold diluted overnight bacterial cultures continued to expand in fresh media at 30 or 37 °C, 250rpm for about 1.5 h until bacteria reached the exponential phase. The OD600 of the bacterial culture was measured, and the density of bacteria was calculated by OD600 times 8*10^8^ /ml. Bacteria were resuspended in cell culture media without any antibiotics, and the concentrations were adjusted to the desired multiplicity of infection (MOI). To prepare monolayer host cells, different epithelial cell lines were seeded in a 24-well plate (200,000 cells per well) or a 96-well plate (40,000 cells per well) the day before the bacterial infection. On the day of infection, the media in the plates were removed and cells were washed with Dulbecco’s Phosphate Buffered Saline (DPBS; Corning, catalog#: 21-031-CV) once. The cell culture media containing bacteria was added into the wells, and this mixture of bacteria and cells was incubated in a humidified incubator at 37 °C, 5% CO_2_. After a 4 h incubation, cells were washed once with DPBS, and fresh media containing 10% FBS and 200 μg/ml gentamicin were added to each well to suppress further bacterial growth. One hour later, a single cell suspension was prepared by trypsin digestion and subject to immunocytochemistry staining for γH2AX Phosphorylation (Ser139). For the in vitro co-culture assay involving D-serine, after diluting overnight pks bacteria ten times with fresh LB broth, 10 mM D-serine was added, and the pks bacteria were expanded at 37 °C for about 1 h until reaching the exponential phase. OD600 and MOI were calculated for these pks bacteria as described above. 10 mM D-serine was further added to the cell culture medium (DMEM, 10% FBS) during the 4-h bacteria-HeLa coculture, followed by DMEM, 10% FBS, and gentamicin for another 1 h before trypsin digestion and staining with anti-gamma H2A. For cell cycle arrest detection, cells were trypsinized after a 24 h incubation with gentamicin-containing media following the 4 h co-culture.

### Immunostaining and flow cytometry

For γH2A.X Phosphorylation (Ser139) staining, detached cells were transferred to a 96-well v-bottom plate, washed with 1x FACS buffer (1x PBS, 5% FBS, 2mM EDTA, 0.05% Sodium Azide) once and fixed with 4% Paraformaldehyde (PFA) at 37 °C for 10 min. After fixation, cells were washed with 1x FACS buffer twice and permeabilized by 0.5% Triton X-100 at 4 °C for 15 min. The antibody staining buffer (1% Bovine Serum Albumin, 0.05% Tween-20 dissolved in 1x PBS) containing 250 ng/ml Alexa Fluor® 647 anti-H2A.X Phospho (Ser139) Antibody (catalog# 613408) was added to cells collected in the 96 v-bottom plate after two washes by FACS buffer. After overnight incubation at 4 °C, the antibody staining buffer was removed from wells, and cells were washed with PBST (0.05% Tween-20 dissolved in 1x PBS) twice. During each wash, the cells were shaken for 10 min at room temperature (RT) at a speed of 400 rpm. Afterward, cells resuspended in 1x FACS buffer were analyzed by an Attune NxT Flow Cytometer (Thermo Fisher).

Experiment-to-experiment variation of MFIs is relatively common, and thus genotoxic indices from different experiments should not necessarily be directly compared.

For cell cycle arrest analysis, detached cells are fixed by 90% ethanol for 20 min at 4 °C after one wash by 1x PBS. 90% ethanol is removed after fixation and the cells are washed 2 times by 1x PBS. Before analyzing in the flow cytometer, cells are incubated with 0.5mL FxCycle™ PI/RNase Staining Solution (Thermo Fisher, Catalog#: F10797) at room temperature for 30 min in the dark.

### Animal experiments

C57BL/6J (Strain #:000664) and C57BL/6J-Apc^Min/J^ (Strain#:002020) mice were initially purchased from the Jackson Laboratory, and C57BL/6J-Apc^Min/J^ were further bred in a specific-pathogen-free (SPF) facility. The offspring were genotyped via the primers provided by the Jackson Laboratory using polymerase chain reaction (PCR) analysis on tail biopsy. All animal experiments were under the IACUC protocol PRO00011467, approved by the committees at the University of Michigan.

For colon adenoma forming experiment, Apc^Min/J^ mice (8 – 10 weeks old, all genders) were pre-treated with antibiotics (2g/L streptomycin) in the drinking water for 3 days (starting from day -10 as indicated in Fig. 4a). Subsequently, drinking water was switched to the one with 1.5% (w/v) dextran sulfate sodium (DSS) for a week. Afterwards, mice were gavaged with 10^9^ CFU NC101 and treatment or control groups (10^9^ CFU Ec^eB-TP1^^14^, 10^9^ CFU Ec^dCas9-sgCtl^, 10^9^ CFU Ec^dCas9-sgclbB_4387^, 10^9^ CFU Ec^dCas9-sgclbC_2313^, 5*10^8^ Ec^dCas9-sgclbB_4387^ + 5*10^8^ Ec^dCas9-sgclbC_2313^, or PBS) in 100ul sterile PBS. NC101 was given weekly for 3 times at days 0, 7, and 14. Treatment and control groups were gavaged twice a week for 6 times in total at days 0, 3, 7, 10, 14, and 17; on days when both gavages occurred, NC101 was gavaged first and then the treatment or control groups 4 h later. Fecal samples were collected two days after oral gavage, snap frozen in liquid nitrogen, and transferred to -80℃ for long-term storage. Body weight was tracked to make sure no mice fell under humane endpoint limits. Mice were sacrificed at day 30 (± 5 days) and colons were harvested, rinsed with PBS, and cut open longitudinally. Tumors were counted and images were captured. Colons were then Swiss rolled in cassettes, fixed in 4% paraformaldehyde (PFA) overnight at 4 °C, and transferred to 70% ethanol. Swiss-rolled colon sections were sent to the University of Michigan Tissue and Molecular Pathology Core for processing, including paraffin-embedding, sectioning, and immunohistochemistry (IHC), hematoxylin and eosin (H&E) staining.

For the safety experiment, C57BL/6 mice (10 – 12 weeks old) were orally gavaged with 10^8^ NC101 and control or treatment groups (either PBS, 10^9^ CFU Ec^dCas9-sgCtl^, 10^9^ CFU Ec^dCas9-sgclbB_4387^, or 10^9^ CFU Ec^dCas9-sgclbC_2313^) in 100 μL PBS. NC101 and treatment and control groups were gavaged 5 times total on days 0, 3, 7, 10, and 14; NC101 was gavaged first and then the treatment or control groups 4 h later. Fecal samples were collected two days after oral gavage, snap frozen in liquid nitrogen, and transferred to - 80℃ for long-term storage. Body weight was tracked to make sure no mice fell under humane endpoint limits. The serum of mice was collected on day 21. Mice were sacrificed at day 21, and colons, kidneys, livers, and spleens were harvested; colons were rinsed with PBS and cut open longitudinally. Colons were then Swiss rolled in cassettes, and the kidney, liver, and spleen samples were also put into cassettes. The cassettes were fixed in 4% paraformaldehyde (PFA) overnight at 4 °C, and transferred to 70% ethanol. Tissue samples were sent to the University of Michigan Tissue and Molecular Pathology Core for processing, including paraffin-embedding, sectioning, and immunohistochemistry (IHC), hematoxylin and eosin (H&E) staining.

### qPCR Analysis

Bacterial strains of interest were inoculated in 2 ml LB and placed in a shaker (30 or 37°C, 250rpm) overnight. The next day, 10-fold diluted overnight bacterial cultures continued to expand in fresh media at 30 or 37 °C, 250rpm for about 1.5 h until bacteria reached the exponential phase. RNA was extracted using the Quick-RNA MiniPrep kit from Zymo Research (Irvine, CA, USA) and treated with the TURBO DNA-free kit (Thermo Fisher Scientific, Waltham, MA, USA). cDNA was synthesized with the EasyQuick RT Master Mix from CWBio (Taizhou, China). qPCR samples were prepared using 10-20ng of cDNA per well and either PowerUp SYBR Green Master Mix (Thermo Fisher Scientific) or FastSYBR Mixture (CWBio) along with the necessary qPCR primers. qPCR Primers specific to clbB, clbC, and 16S genes were designed by NCBI Primer BLAST, and the specificity of qPCR products was verified by 2% agarose gel and Sanger sequencing.

The clbP and 16S primers for fecal gDNA were published previously ^52^. Samples were run and analyzed on a 7500 Fast Dx Real-Time PCR System (Thermo Fisher) using a threshold value of 0.1. The detailed qPCR program is: 50 °C for 2 min, 95 °C for 2 min, 40 cycles of 95 °C for 5 s and 60°C for 30 s, and a final cycle of 95°C for 15 s, 60 °C for 1 min, and 95 °C for 15 s. Relative gene expression was calculated by the 2^−ΔΔCt^ method using 16S rRNA as the reference gene. Primers used for RT-PCR are shown in Tables S3.

### RNA sequencing and data analysis

RNA sample quality was assessed using a Nanodrop (Thermo Fisher) and Agilent 5400 (Agilent Technologies, Santa Clara, CA, USA). Prokaryotic mRNA sequencing was performed using the NovaSeq PE150 platform (Illumina, San Diego, CA, USA) at the Novogene facility (Sacramento, CA, USA). The library was prepared by a Ribo-Zero protocol (250−300 bp insert strand-specific library with rRNA removal using NEB Ribo-Zero Magnetic Kit). Paired-end sequencing produced 150 bp reads to a depth of ∼2G output per sample. Sequences were mapped to a reference genome, NC101 (GenBank accession: CP072787.1), using a Bowtie2 pipeline adjusted for paired-end sequencing. Differential gene expression was analyzed using the DESeq2 pipeline in RStudio as previously described ^57^. Due to the presence of the eB-TP114 conjugative plasmid, the total mapping rates with respect to the annotated genome for CP072787.1 were ∼ 90%. The false discovery rate (FDR) was set to 5% and genes with a log_2_FoldChange of >0.5 or <-0.5 and a p-value < 0.01 were considered significant. All reported data are representative of three biological replicates.

### 16S rRNA gene sequencing and microbiome analysis

16s rRNA analyses were performed by Novogene, China, as we previously described ^58^. Briefly, samples are checked for quality control using PCR amplification of the 16S V3/V4 regions. For library construction PCR amplification of targeted regions was performed by using specific primers connecting with barcodes. The PCR products of proper size were selected through agarose gel electrophoresis. The same amount of PCR products from each sample was pooled, end-repaired, A-tailed, and further ligated with illumina adapters. Libraries were sequenced on a paired-end Illumina platform. The library was checked with Qubit and real-time PCR for quantification, while a bioanalyzer was used for size distribution detection. Quantified libraries were pooled and sequenced on Illumina platforms according to the effective library concentration and data amount required. The amplicon was sequenced on an Illumina platform to generate paired-end raw reads (Raw PE), and then merged and pre-treated to obtain Clean Tags. The chimeric sequences in Clean Tags were detected and removed to obtain the Effective Tags, which can be used for subsequent analysis.

### Bacterial culture and extraction of yersiniabactin (ybt)

*E. coli* NC101 wild type (WT) and *irp2* knockout (KO) strains were grown in nutrient-rich conditions (Luria-Bertani), nutrient-limited conditions (M9 minimal medium (per liter: 6.8 g Na_2_HPO_4_; 3 g KH_2_PO_4_; 0.5 g NaCl; 1 g NH_4_Cl, 0.1 mM CaCl2,1 mM MgSO_4_, 0.2% glucose and supplemented with 1 mg/ml biotin and 1 mg/ml thiamine)) for 24 hours at 37 °C. Overnight cultures were spun down at 5,000 rpm for 10 minutes, and the supernatant was stored at -80°C for further analysis. Supernatants from WT and *irp2* KO *E. coli* NC101 cultures were extracted onto pre-washed solid phase extraction (SPE) cartridges. SPE cartridges were activated three times with methanol (MeOH, 3 × 3 mL), then washed twice with water and 0.1% formic acid (FA, 3 × 3 mL). The sample was loaded dropwise (steady single dripping) onto SPE cartridges, and then the cartridges were washed with water and 0.1% FA (3 × 3 mL). The sample was eluted into 2 mL of MeOH, and then it was dried under nitrogen gas at room temperature. Samples were weighed and reconstituted with 80% MeOH/20% water + 0.1% FA. 5 µl of the sample was injected per run.

### LC-MS detection of ybt

For LC-MS/MS analysis, 5 µl were injected into an Agilent Vanquish UHPLC system coupled to a Q6545A mass spectrometer (Agilent, Santa Clara, United States). For the chromatographic separation, a C18 porous core column (Infinity Lab poroshell 120 EC-C18, 50 × 2.1 mm, 1.9 μm particle size, 120 Angstrom pore size, Agilent, Santa Clara, United States) was used. For gradient elution, a high-pressure binary gradient system was used. The mobile phase consisted of solvent A (H2O+ 0.1% FA) and solvent B (100% acetonitrile). The flow rate was set to 0.5 ml/min. After injection, the samples were eluted with the following linear gradient: 0–0.5 min 5% B, 0.5–5 min 5–100% B, followed by a 2 min washout phase at 100% B and a 3 min re-equilibration phase at 5% B. Data-dependent acquisition (DDA) of MS/MS spectra was performed in positive mode. ESI parameters were set to 11 L/min sheath gas flow, 11 L/min sheath gas flow, and 350 °C gas temperature. The capillary voltage was set to 3.5 kV, and the inlet capillary temperature was set to 320 °C.

### Statistical Analysis

Statistical analysis was performed using GraphPad PRISM (San Diego, CA, USA). Specific tests are indicated in figure captions. P values < 0.05 were considered statistically significant. In the figure legends, n = sample number per group in one biological replicate, N = number of biological replicates.

## Data Availability

All data generated during this study are available within the paper.

## References

1 Metwaly, A. et al. A Consensus Statement on establishing causality, therapeutic applications and the use of preclinical models in microbiome research. Nat Rev Gastroenterol Hepatol 22, 343–356 (2025). 10.1038/s41575-025-01041-3

2 Corander, J., Hanage, W. P. & Pensar, J. Causal discovery for the microbiome. Lancet Microbe 3, e881–e887 (2022). 10.1016/S2666-5247(22)00186-0

3 Pleguezuelos-Manzano, C. et al. Mutational signature in colorectal cancer caused by genotoxic pks(+) E. coli. Nature 580, 269–273 (2020). 10.1038/s41586-020-2080-8

4 Zhang, J. et al. Coculture of primary human colon monolayer with human gut bacteria. Nat Protoc 16, 3874–3900 (2021). 10.1038/s41596-021-00562-w

5 Puschhof, J. et al. Intestinal organoid cocultures with microbes. Nat Protoc 16, 4633–4649 (2021). 10.1038/s41596-021-00589-z

6 Diaz-Gay, M. et al. Geographic and age variations in mutational processes in colorectal cancer. Nature (2025). 10.1038/s41586-025-09025-8

7 Jans, M. et al. Colibactin-driven colon cancer requires adhesin-mediated epithelial binding. Nature 635, 472–480 (2024). 10.1038/s41586-024-08135-z

8 Silpe, J. E., Wong, J. W. H., Owen, S. V., Baym, M. & Balskus, E. P. The bacterial toxin colibactin triggers prophage induction. Nature 603, 315–320 (2022). 10.1038/s41586-022-04444-3

9 Dejea, C. M. et al. Patients with familial adenomatous polyposis harbor colonic biofilms containing tumorigenic bacteria. Science 359, 592–597 (2018). 10.1126/science.aah3648

10 Nougayrede, J. P. et al. Escherichia coli induces DNA double-strand breaks in eukaryotic cells. Science 313, 848–851 (2006). 10.1126/science.1127059

11 Fais, T., Delmas, J., Barnich, N., Bonnet, R. & Dalmasso, G. Colibactin: More Than a New Bacterial Toxin. Toxins 10 (2018). ARTN 15110.3390/toxins10040151

12 Arima, K. et al. Western-Style Diet, Polyketide Synthase (pks) Island-Carrying Escherichia coli, and Colorectal Cancer: Analyses From Two Large Prospective Cohort Studies. Gastroenterology (2022). 10.1053/j.gastro.2022.06.054

13 Jinek, M. et al. A programmable dual-RNA-guided DNA endonuclease in adaptive bacterial immunity. Science 337, 816–821 (2012). 10.1126/science.1225829

14 Cong, L. et al. Multiplex genome engineering using CRISPR/Cas systems. Science 339, 819–823 (2013). 10.1126/science.1231143

15 Komor, A. C., Kim, Y. B., Packer, M. S., Zuris, J. A. & Liu, D. R. Programmable editing of a target base in genomic DNA without double-stranded DNA cleavage. Nature 533, 420–424 (2016). 10.1038/nature17946

16 Richardson, C. D., Ray, G. J., DeWitt, M. A., Curie, G. L. & Corn, J. E. Enhancing homology-directed genome editing by catalytically active and inactive CRISPR-Cas9 using asymmetric donor DNA. Nat Biotechnol 34, 339–344 (2016). 10.1038/nbt.3481

17 Brodel, A. K. et al. In situ targeted base editing of bacteria in the mouse gut. Nature 632, 877–884 (2024). 10.1038/s41586-024-07681-w

18 Lam, K. N. et al. Phage-delivered CRISPR-Cas9 for strain-specific depletion and genomic deletions in the gut microbiome. Cell Rep 37, 109930 (2021). 10.1016/j.celrep.2021.109930

19 Dougherty, M. W. & Jobin, C. Shining a Light on Colibactin Biology. Toxins (Basel) 13 (2021). 10.3390/toxins13050346

20 Baldelli, V., Scaldaferri, F., Putignani, L. & Del Chierico, F. The Role of Enterobacteriaceae in Gut Microbiota Dysbiosis in Inflammatory Bowel Diseases. Microorganisms 9 (2021). ARTN 69710.3390/microorganisms9040697

21 Behnsen, J. et al. Siderophore-mediated zinc acquisition enhances enterobacterial colonization of the inflamed gut. Nature Communications 12 (2021). ARTN 701610.1038/s41467-021-27297-2

22 Hamilton, T. A. et al. Efficient inter-species conjugative transfer of a CRISPR nuclease for targeted bacterial killing. Nat Commun 10, 4544 (2019). 10.1038/s41467-019-12448-3

23 Rottinghaus, A. G., Ferreiro, A., Fishbein, S. R. S., Dantas, G. & Moon, T. S. Genetically stable CRISPR-based kill switches for engineered microbes. Nat Commun 13, 672 (2022). 10.1038/s41467-022-28163-5

24 Auvray, F. et al. Insights into the acquisition of the pks island and production of colibactin in the Escherichia coli population. Microb Genom 7 (2021). 10.1099/mgen.0.000579

25 Neil, K. et al. High-efficiency delivery of CRISPR-Cas9 by engineered probiotics enables precise microbiome editing. Mol Syst Biol 17, e10335 (2021). 10.15252/msb.202110335

26 Oliero, M. et al. Inulin impacts tumorigenesis promotion by colibactin-producing Escherichia coli in Apc(Min/+) mice. Front Microbiol 14, 1067505 (2023). 10.3389/fmicb.2023.1067505

27 Arthur, J. C. et al. Intestinal inflammation targets cancer-inducing activity of the microbiota. Science 338, 120–123 (2012). 10.1126/science.1224820

28 Kim, S. C. et al. Variable phenotypes of enterocolitis in interleukin 10-deficient mice monoassociated with two different commensal bacteria. Gastroenterology 128, 891–906 (2005). 10.1053/j.gastro.2005.02.009

29 Dubois, D. et al. ClbP is a prototype of a peptidase subgroup involved in biosynthesis of nonribosomal peptides. J Biol Chem 286, 35562–35570 (2011). 10.1074/jbc.M111.221960

30 Dziubanska-Kusibab, P. J. et al. Colibactin DNA-damage signature indicates mutational impact in colorectal cancer. Nat Med 26, 1063–1069 (2020). 10.1038/s41591-020-0908-2

31 Iftekhar, A. et al. Genomic aberrations after short-term exposure to colibactin-producing E. coli transform primary colon epithelial cells. Nat Commun 12, 1003 (2021). 10.1038/s41467-021-21162-y

32 Mah, L. J., El-Osta, A. & Karagiannis, T. C. gammaH2AX: a sensitive molecular marker of DNA damage and repair. Leukemia 24, 679–686 (2010). 10.1038/leu.2010.6

33 Sharma, A., Singh, K. & Almasan, A. Histone H2AX phosphorylation: a marker for DNA damage. Methods Mol Biol 920, 613–626 (2012). 10.1007/978-1-61779-998-3_40

34 Bonner, W. M. et al. GammaH2AX and cancer. Nat Rev Cancer 8, 957–967 (2008). 10.1038/nrc2523

35 Sheng, Y. et al. Insertion sequence transposition inactivates CRISPR-Cas immunity. Nat Commun 14, 4366 (2023). 10.1038/s41467-023-39964-7

36 Qi, L. S. et al. Repurposing CRISPR as an RNA-guided platform for sequence-specific control of gene expression. Cell 152, 1173–1183 (2013). 10.1016/j.cell.2013.02.022

37 Homburg, S., Oswald, E., Hacker, J. & Dobrindt, U. Expression analysis of the colibactin gene cluster coding for a novel polyketide in Escherichia coli. FEMS Microbiol Lett 275, 255–262 (2007). 10.1111/j.1574-6968.2007.00889.x

38 Neil, K., Allard, N., Jordan, D. & Rodrigue, S. Assembly of large mobilizable genetic cargo by double recombinase operated insertion of DNA (DROID). Plasmid 104, 102419 (2019). 10.1016/j.plasmid.2019.102419

39 Depardieu, F. & Bikard, D. Gene silencing with CRISPRi in bacteria and optimization of dCas9 expression levels. Methods 172, 61–75 (2020). 10.1016/j.ymeth.2019.07.024

40 Cui, L. et al. A CRISPRi screen in E. coli reveals sequence-specific toxicity of dCas9. Nat Commun 9, 1912 (2018). 10.1038/s41467-018-04209-5

41 Cho, S. et al. High-Level dCas9 Expression Induces Abnormal Cell Morphology in Escherichia coli. ACS Synth Biol 7, 1085–1094 (2018). 10.1021/acssynbio.7b00462

42 Bossuet, N. et al. Oxygen concentration modulates colibactin production. Gut Microbes 15, 2222437 (2023). 10.1080/19490976.2023.2222437

43 Massip, C. et al. Deciphering the interplay between the genotoxic and probiotic activities of Escherichia coli Nissle 1917. PLoS Pathog 15, e1008029 (2019). 10.1371/journal.ppat.1008029

44 Pérez-Berezo, T. et al. Identification of an analgesic lipopeptide produced by the probiotic strain Nissle 1917. Nature Communications 8 (2017). ARTN 131410.1038/s41467-017-01403-9

45 Rosendahl Huber, A., et al. Improved detection of colibactin-induced mutations by genotoxic E. coli in organoids and colorectal cancer. Cancer Cell 42, 487–496 e486 (2024). 10.1016/j.ccell.2024.02.009

46 Bossuet-Greif, N. et al. Escherichia coli ClbS is a colibactin resistance protein. Mol Microbiol 99, 897–908 (2016). 10.1111/mmi.13272

47 Hallam, J. C. et al. D-Serine reduces the expression of the cytopathic genotoxin colibactin. Microb Cell 10, 63–77 (2023). 10.15698/mic2023.03.793

48 Volpe, M. R. et al. A small molecule inhibitor prevents gut bacterial genotoxin production. Nat Chem Biol 19, 159–167 (2023). 10.1038/s41589-022-01147-8

49 Putze, J. et al. Genetic structure and distribution of the colibactin genomic island among members of the family Enterobacteriaceae. Infect Immun 77, 4696–4703 (2009). 10.1128/IAI.00522-09

50 Carniel, E. The Yersinia high-pathogenicity island: an iron-uptake island. Microbes Infect 3, 561–569 (2001). 10.1016/s1286-4579(01)01412-5

51 Wami, H. et al. Insights into evolution and coexistence of the colibactin- and yersiniabactin secondary metabolite determinants in enterobacterial populations. Microb Genomics 7 (2021). ARTN 00057710.1099/mgen.0.000577

52 Yang, Y., Gharaibeh, R. Z., Newsome, R. C. & Jobin, C. Amending microbiota by targeting intestinal inflammation with TNF blockade attenuates development of colorectal cancer. Nat Cancer 1, 723–734 (2020). 10.1038/s43018-020-0078-7

53 Lucas, C. et al. Autophagy of Intestinal Epithelial Cells Inhibits Colorectal Carcinogenesis Induced by Colibactin-Producing Escherichia coli in Apc(Min/+) Mice. Gastroenterology 158, 1373–1388 (2020). 10.1053/j.gastro.2019.12.026

54 Beesley, S. et al. D-serine mitigates cell loss associated with temporal lobe epilepsy. Nat Commun 11, 4966 (2020). 10.1038/s41467-020-18757-2

55 Zhang, X. Q. et al. D-serine reconstitutes synaptic and intrinsic inhibitory control of pyramidal neurons in a neurodevelopmental mouse model for schizophrenia. Nat Commun 14, 8255 (2023). 10.1038/s41467-023-43930-8

56. Tronnet, S., et al. The Genotoxin Colibactin Shapes Gut Microbiota in Mice. mSphere 5 (2020). 10.1128/mSphere.00589-20

57 Yang, M. et al. Targeting Fusobacterium nucleatum through chemical modifications of host-derived transfer RNA fragments. ISME J 17, 880–890 (2023). 10.1038/s41396-023-01398-w

58 Yang, M. et al. Engineered Bacillus subtilis as Oral Probiotics To Enhance Clearance of Blood Lactate. ACS Synth Biol 14, 101–112 (2025). 10.1021/acssynbio.4c00399

