## Extended Figure for "Programmable conjugative CRISPR interference targeting genotoxin in the gut"

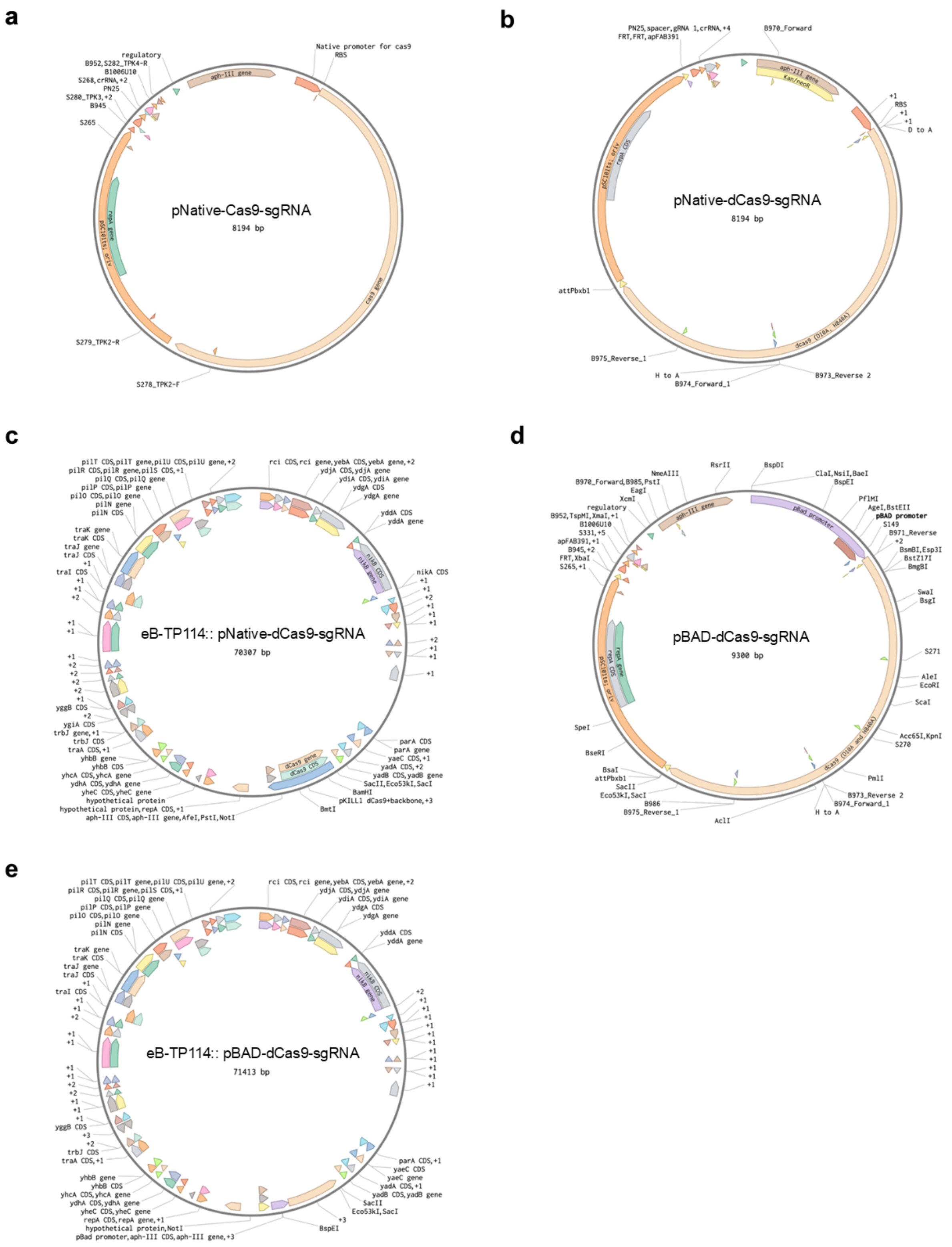


**Extended Figure 1,** Maps of replicative and conjugative plasmids.

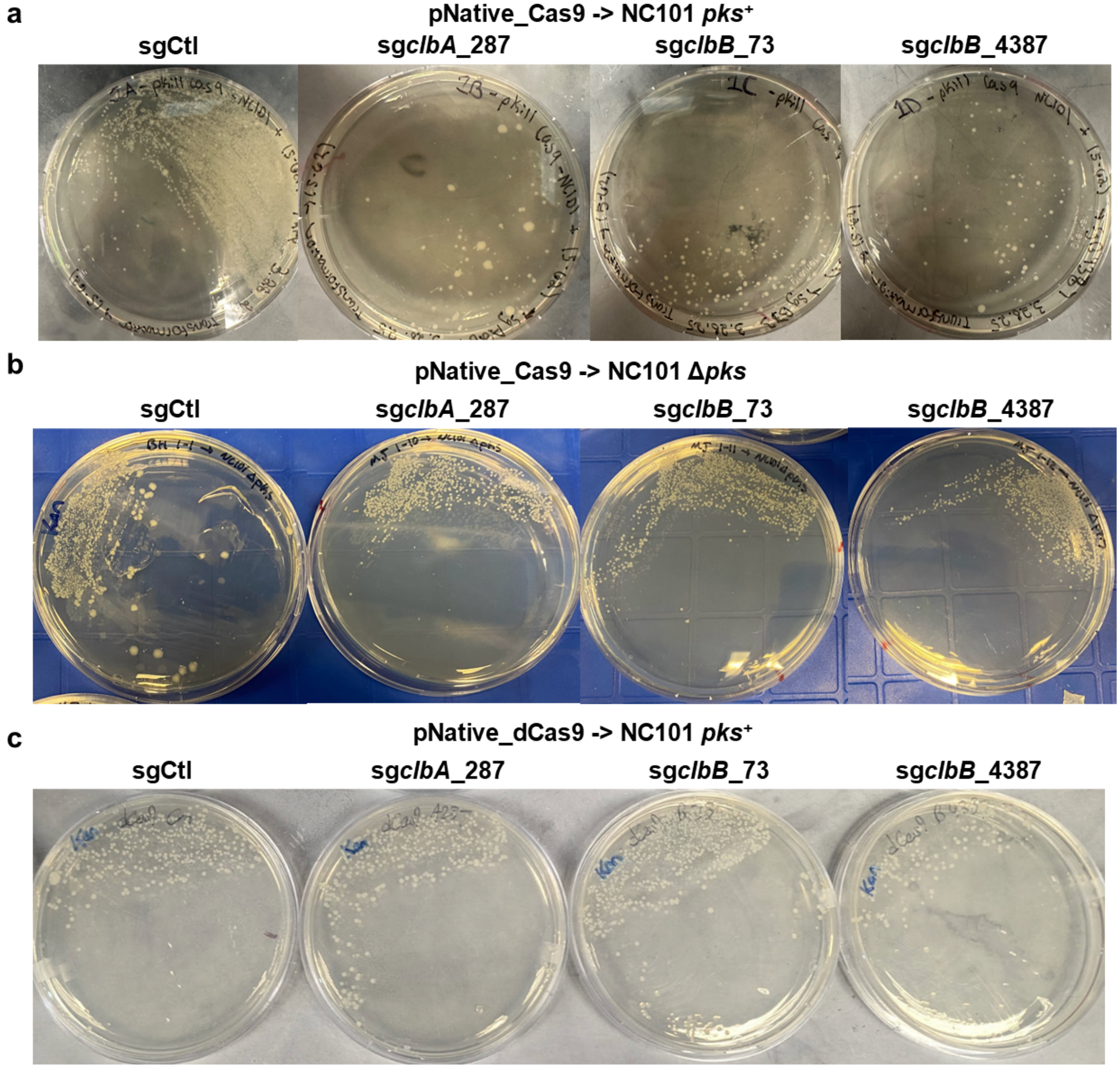


**Extended Figure 2,** Colony growth on lysogeny broth (LB) agar plates with kanamycin antibiotics shows the antimicrobial effects of pNative_Cas9 plasmids with *pks*-targeting sgRNA (**a**). This antimicrobial effect is *pks*-dependent, as transformation of NC101 Δ*pks E. coli* with pNative_Cas9 plasmids shows no effect on colony growth on LB agar plates with kanamycin (**b**). pNative_dCas9 shows no such antimicrobial effects (**c**). Results are presentative of three independent replicates.

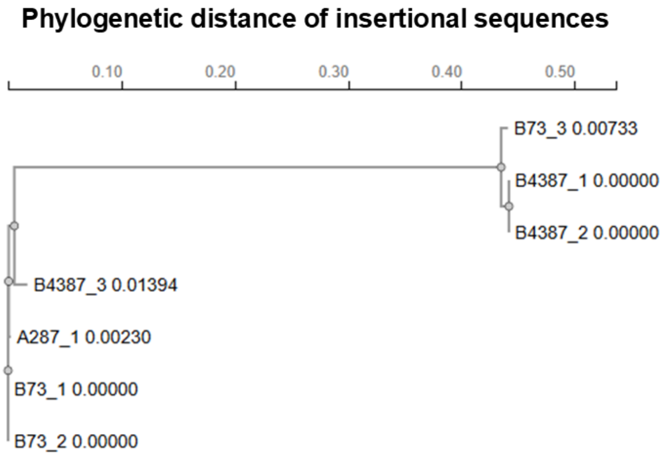


**Extended Figure 3,** Phylogenetic distance after alignment of insertional sequences discovered within the open reading frame of the Cas9 gene. The sequence alignment and phylogenetic tree were generated by Clustal Omega (multiple sequence alignment).

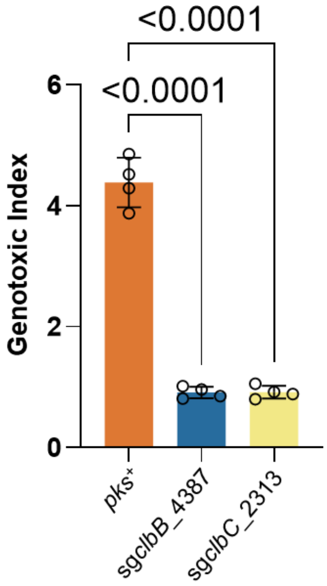


**Extended Figure 4,** NC101 *pks*$^{+}$ cells conjugated with eB-TP114-pBAD_dCas9 constructs cultured in anaerobic conditions prior to the HeLa co-culture can effectively lower the genotoxic index as compared to positive controls. MOI = 100, n = four technical repeats per independent biological replicate, N = two independent biological replicates. Data are means ± SD. Significance values calculated using one-way ANOVA with Dunnett's multiple comparisons test.

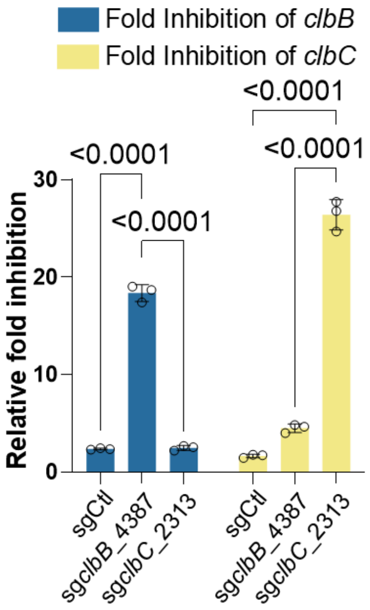


**Extended Figure 5,** $2^{-\Delta\Delta Ct}$ qPCR analysis of *pks*$^{+}$ *E. coli* Nissile conjugated with eB-TP114-pBAD_dCas9 plasmids show highly specific targeting of the *pks* island. n = three technical repeats per independent biological replicate, N = two independent biological replicates. Data are means ± SD. Significance values calculated using two-way ANOVA with Tukey's multiple comparison test.

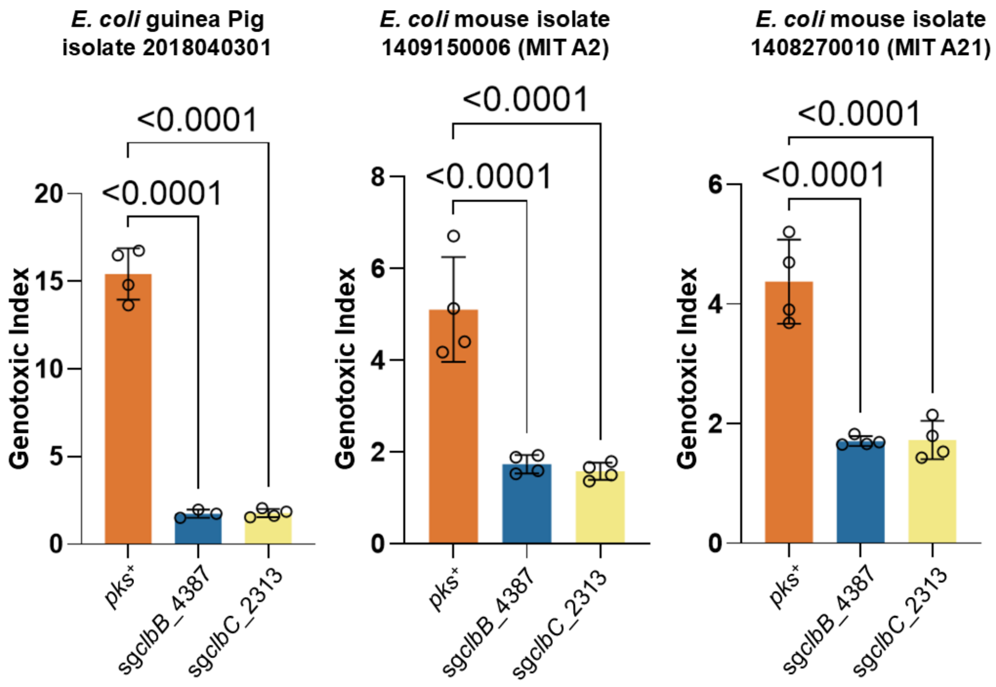


**Extended Figure 6,** eB-TP114-pBAD_dCas9 strains transformed into *pks*+ bacteria extracted from different mammalian hosts significantly lower genotoxicity in HeLa cells after 4 h co-culture as compared to positive controls. MOI = 100, n = four technical repeats per independent biological replicate, N = two independent biological replicates. Data are means ± SD. Significance values calculated using one-way ANOVA with Dunnett's multiple comparisons test.

1063

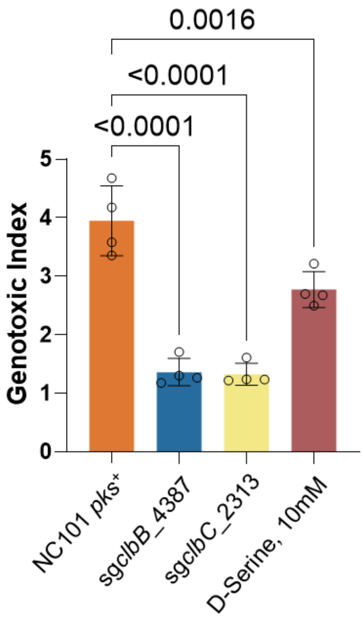


**Extended Figure 7,** NC101 *pks*$^{+}$ conjugated with eB-TP114-pBAD_dCas9 constructs can more effectively lower the genotoxic index as compared to positive controls and the small molecule inhibitor D-Serine. MOI = 100, n = four technical repeats per independent biological replicate, N = two independent biological replicates. Data are means ± SD. Significance values calculated using one-way ANOVA with Dunnett's multiple comparisons test.

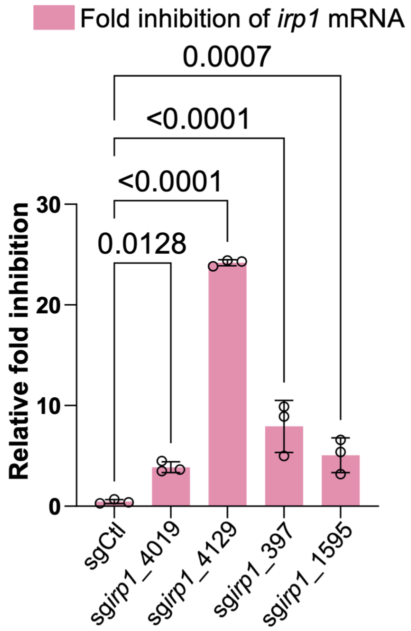


**Extended Figure 8**, *pks*$^{+}$ NC101 *E. coli* are conjugated with eB-TP114-pBAD_dCas9 plasmids encoding four different *irp1*-specific sgRNAs or the non-targeting sgCtl. $2^{-\Delta\Delta Ct}$ qPCR analysis of *irp1* mRNA shows specific targeting of the *irp1* gene, which is necessary for the biosynthesis of yersiniabactin. n = three technical repeats per independent biological replicate, N = two independent biological replicates. Data are means ± SD. Significance values calculated using two-way ANOVA with Dunnett's multiple comparison test.

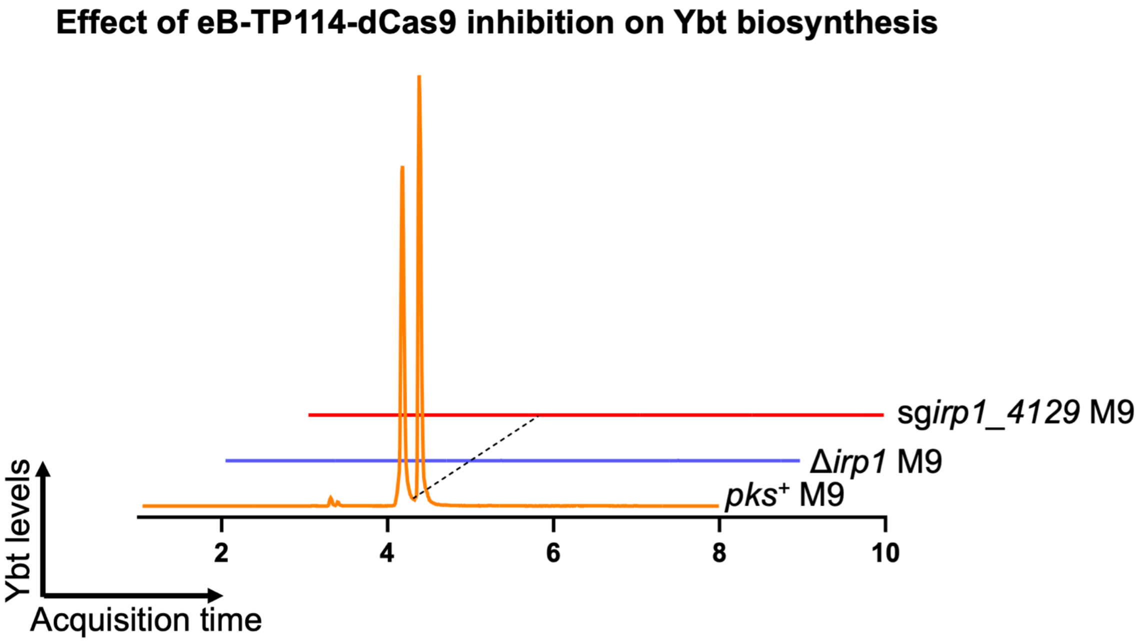


**Extended Figure 9,** Liquid chromatography-mass spectrometry-quadrupole time of flight (LC-MS-QTOF) analysis of *pks*$^+$ NC101 *E. coli* conjugated with eB-TP114-pBAD_dCas9-sg*irp1*_4129 show complete disruption of the biosynthesis of yersiniabactin (ybt) through the silencing of the *irp1* gene as compared to positive and negative controls grown in minimal M9 broth (M9). Results are representative of two independent experiments.

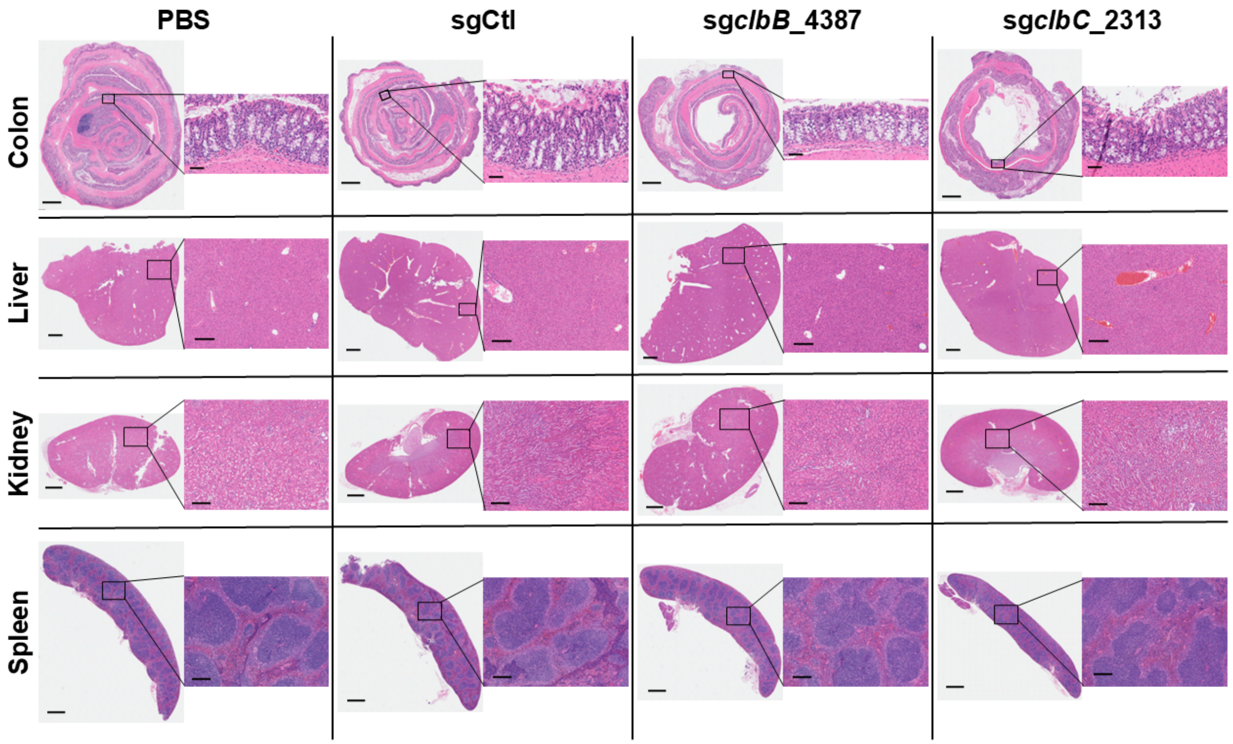


**Extended Figure 10,** H&E staining of colon Swiss rolls, livers, kidneys, and spleens show similar phenotypes with no obvious signs of inflammation. Images are representative samples from each treatment group. Scale bars on colon full zoom images are 800µm; scale bars on colon insets are 50µm. Scale bars on liver, kidney, and spleen full zoom images are 1000µm; scale bars on liver, kidney, and spleen insets are 200µm.

1070

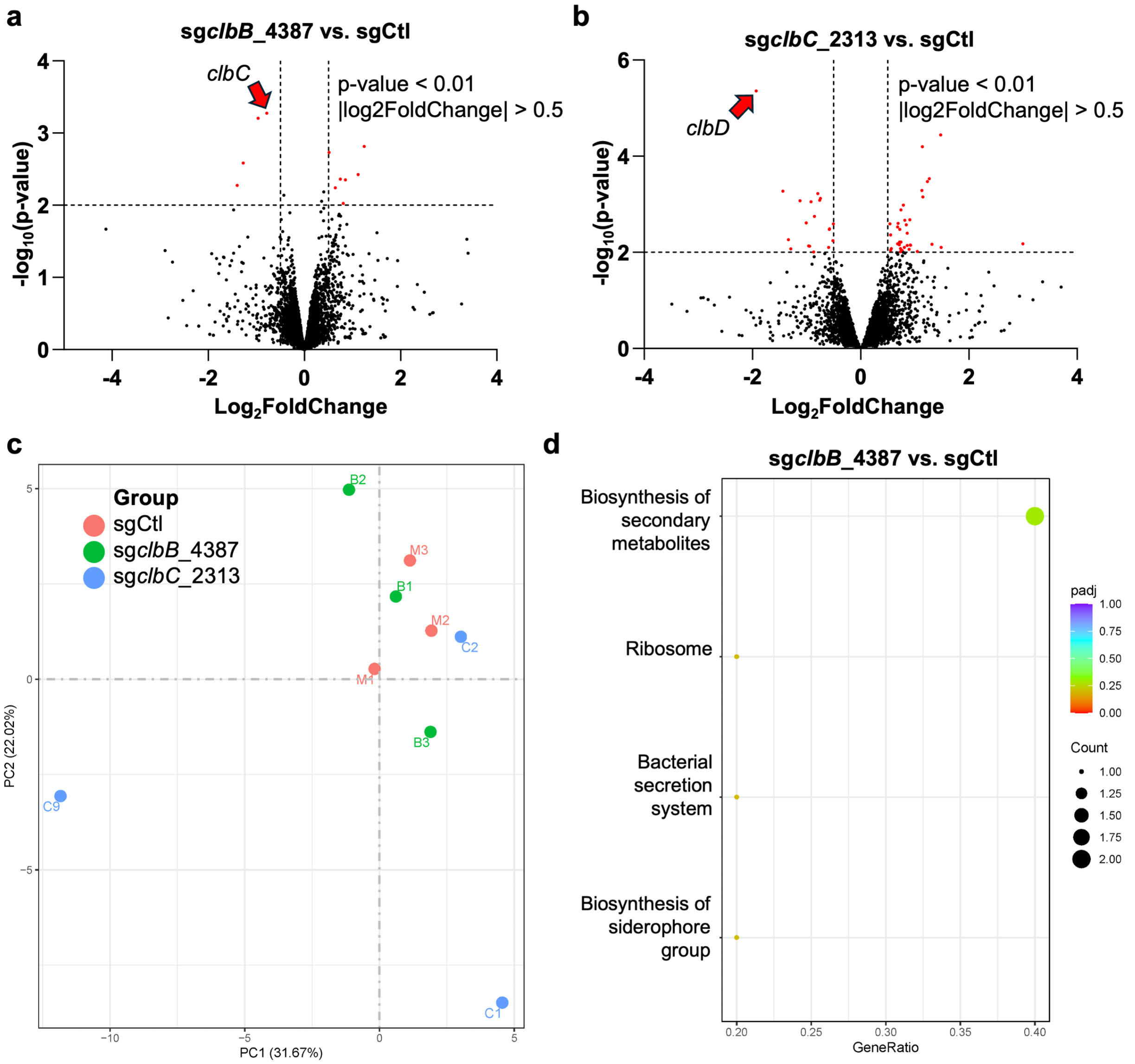


**Extended Figure 11,** RNAseq analysis of sg*clbB*_4387 and sg*clbC*_2313 vs. sgCtl show that *clbC* and *clbD* are the two genes most statistically significantly downregulated (**a** and **b**, respectively), but sg*clbC*_2313 significantly affects the expression of many more genes than sg*clbB*_4387 (19 downregulations and 35 upregulations vs. 4 downregulations and 7 upregulations, respectively). Principal component analysis reinforces this observation, as sgCtl and sg*clbB*_4387 are much more closely clustered than sg*clbC*_2313 (**c**). The associated KEGG plots show that sg*clbB*_4387 downregulates the biosynthesis of secondary metabolites (including, in this case, colibactin) (**d**).

1071

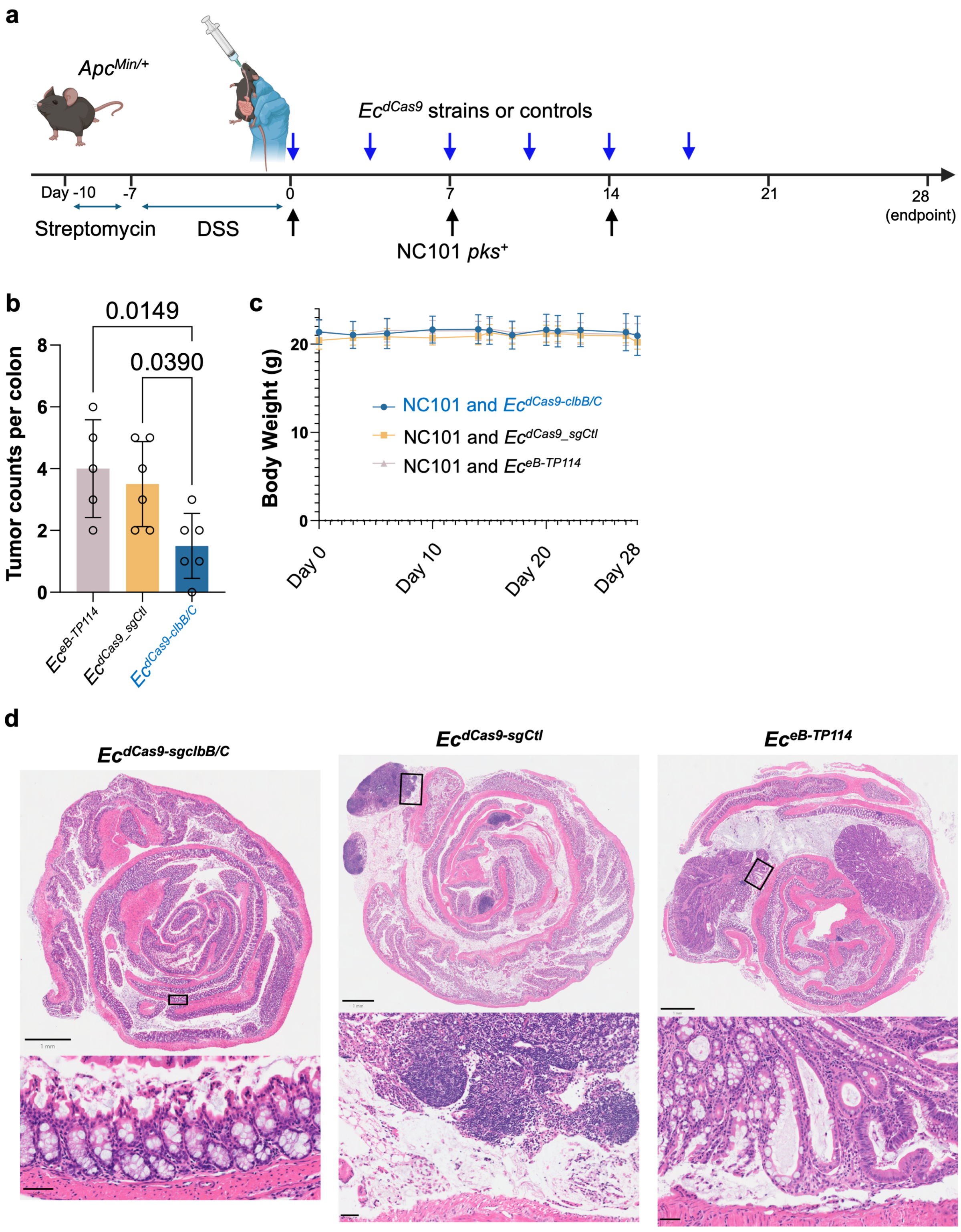


**Extended Figure 12, a,** the cocktail strategy combining equal mixture of $Ec^{dCas9\text{-}sgclbB\_4387}$ and $Ec^{dCas9\text{-}sgclbC\_2313}$ in DSS/$Apc^{Min/+}$ mice. Mice received one of three donor strains by oral gavage: (1) $Ec^{eB\text{-}TP114}$ (eB-TP114 lacking dCas9 and sgRNA), (2) $Ec^{dCas9\text{-}sgCtl}$, and 3) an equal mixture of $Ec^{dCas9\text{-}sgclbB\_4387}$ and $Ec^{dCas9\text{-}sgclbC\_2313}$, denoted as '$Ec^{dCas9\text{-}clbB/C}$'). **b**, $Ec^{dCas9\text{-}clbB/C}$ significantly lowers tumor counts at time of sacrifice compared to $Ec^{eB\text{-}TP114}$ or $Ec^{dCas9\text{-}sgCtl}$. Data are means ± SD. Significance values calculated using one-way ANOVA with Dunnett's multiple comparisons test. **c**, Body weight changes over the course of treatment. **d**, representative H&E staining of colon Swiss rolls. Full image scale bars are 1mm, inset zoom bars are 50 µm.

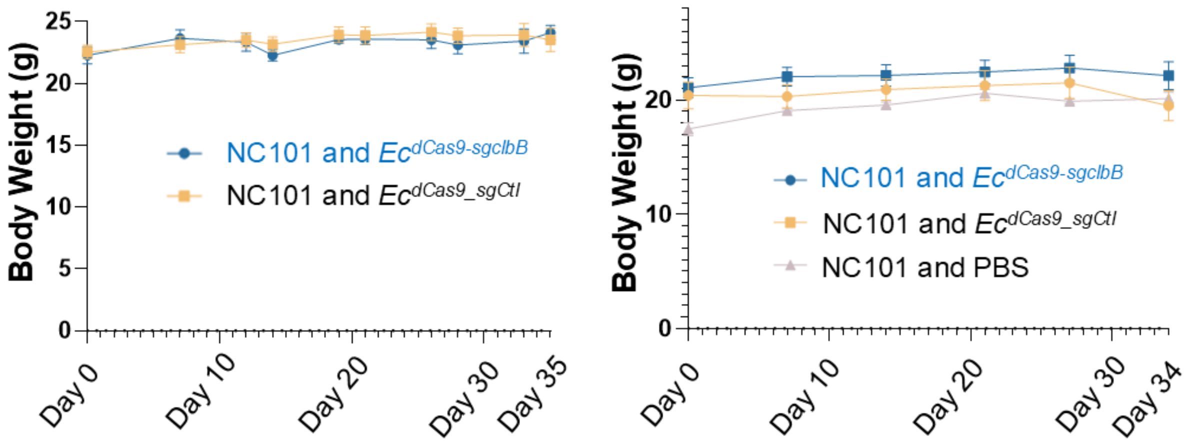


**Extended Figure 13,** Body weight records from the two *in vivo* efficacy studies comparing $Ec^{dCas9\text{-}sgclbB\_4387}$ and $Ec^{dCas9\text{-}sgCtl}$ show stable weights during the experimental timeframes.

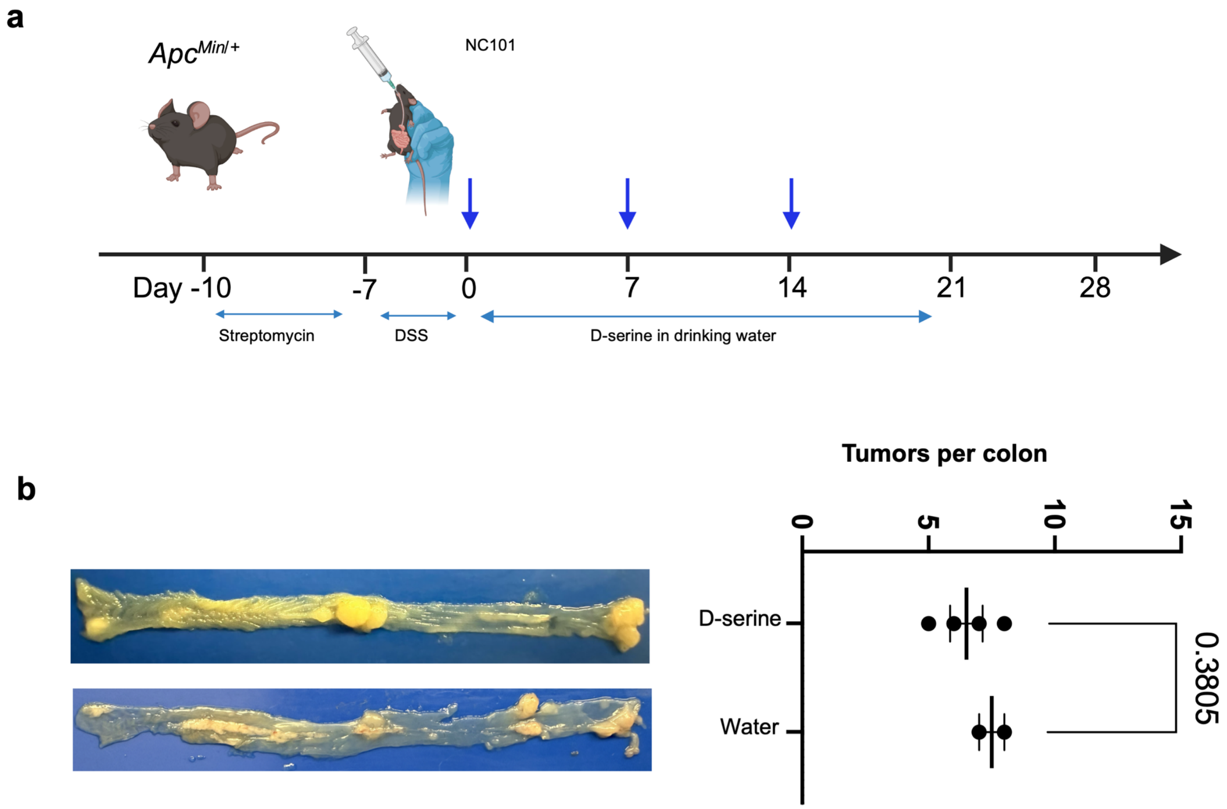


**Extended Figure 14,** D-serine supplementation does not suppress NC101-induced tumorigenesis compared with water controls. **a**, Timeline of DSS/*Apc*$^{\mathrm{Min}/+}$ mice treated with NC101 in combination with D-serine in drinking water. *Apc*$^{\mathrm{Min}/+}$ mice were pretreated with 2 g/L streptomycin in drinking water for three days for bacterial colonization, followed by 2% DSS in water for one week to induce colitis. Next, the mice were infected with NC101 weekly for three weeks, with concurrent 600 mg/L D-serine or water treatment. **b**, Representative tumors in the colon on day 28 and quantification of colonic tumor counts. Data are means ± SD using two-sided unpaired t test.

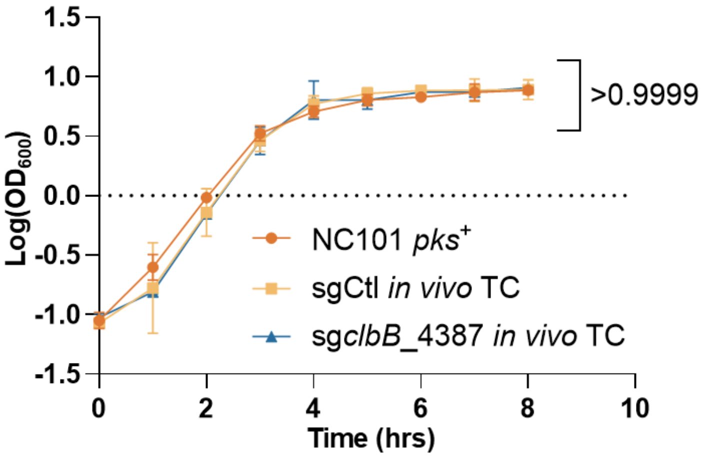


**Extended Figure 15,** Growth curve analysis of *pks*+ NC101 *E. coli* and *in vivo* transconjugates (TC) show no statistically significant difference between cultures. n = 2 technical repeats, N = 2 biological replicates. Data are means ± SD. Significance values calculated using nonlinear regression analysis.

1078

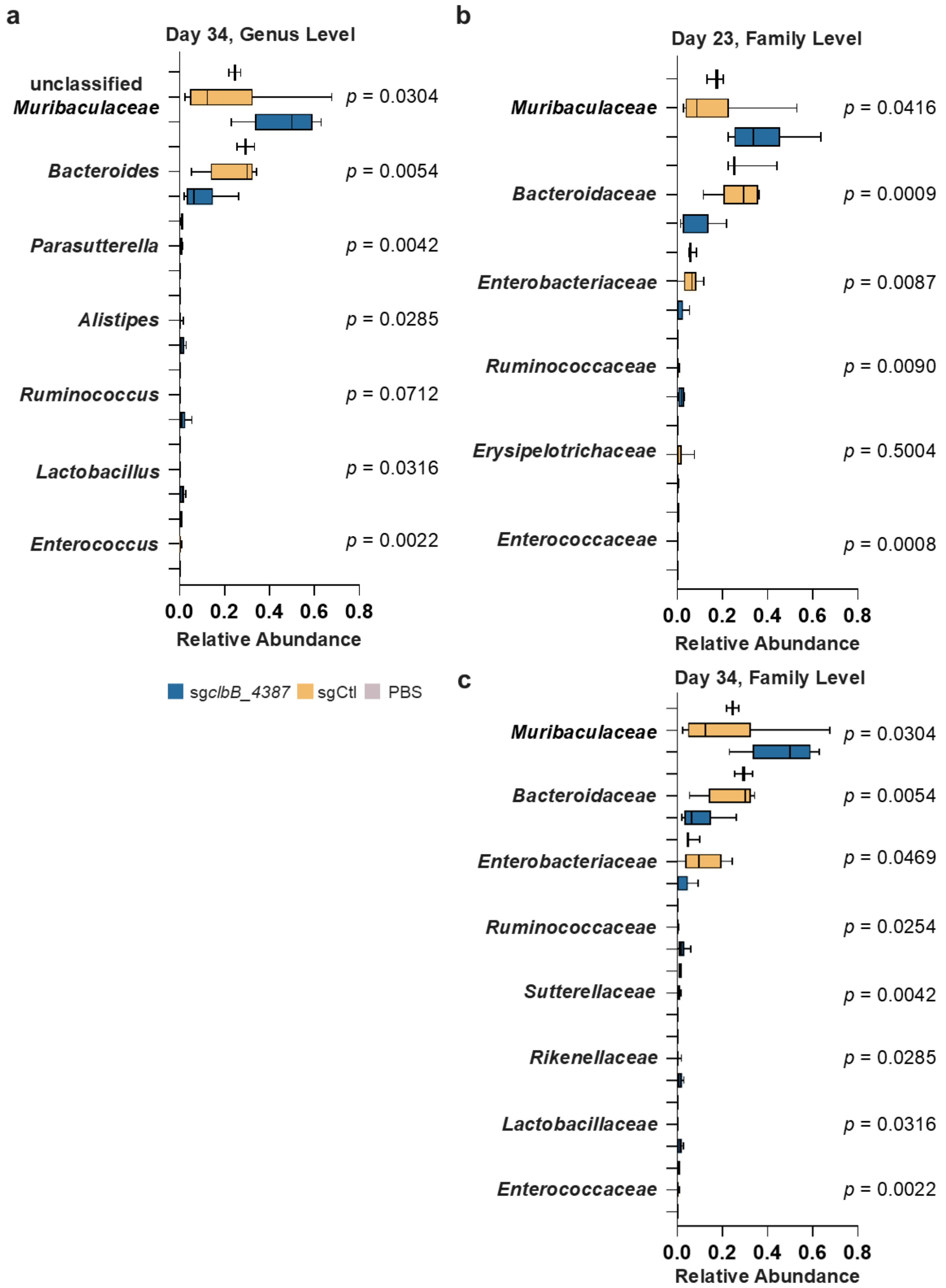


**Extended Figure 16**, Top differential abundance (%) at the genus level (**a**) and family level (**b** and **c**) from fecal microbiota in different groups at Day 23 (**b**) and Day 34 (**a** and **c**). The family level shows significant reduction of *Enterobacteriaceae* by $Ec^{dCas9\text{-}sgclbB\_4387}$ compared to the vehicle and $Ec^{dCas9\text{-}sgCtl}$ control at both mid and endpoints. In box plots, the bounds of the box indicate the 25th–75th percentiles, the center line indicates the median, and whiskers extend from the minimum to the maximum values. Statistical values calculated using one-way ANOVA.

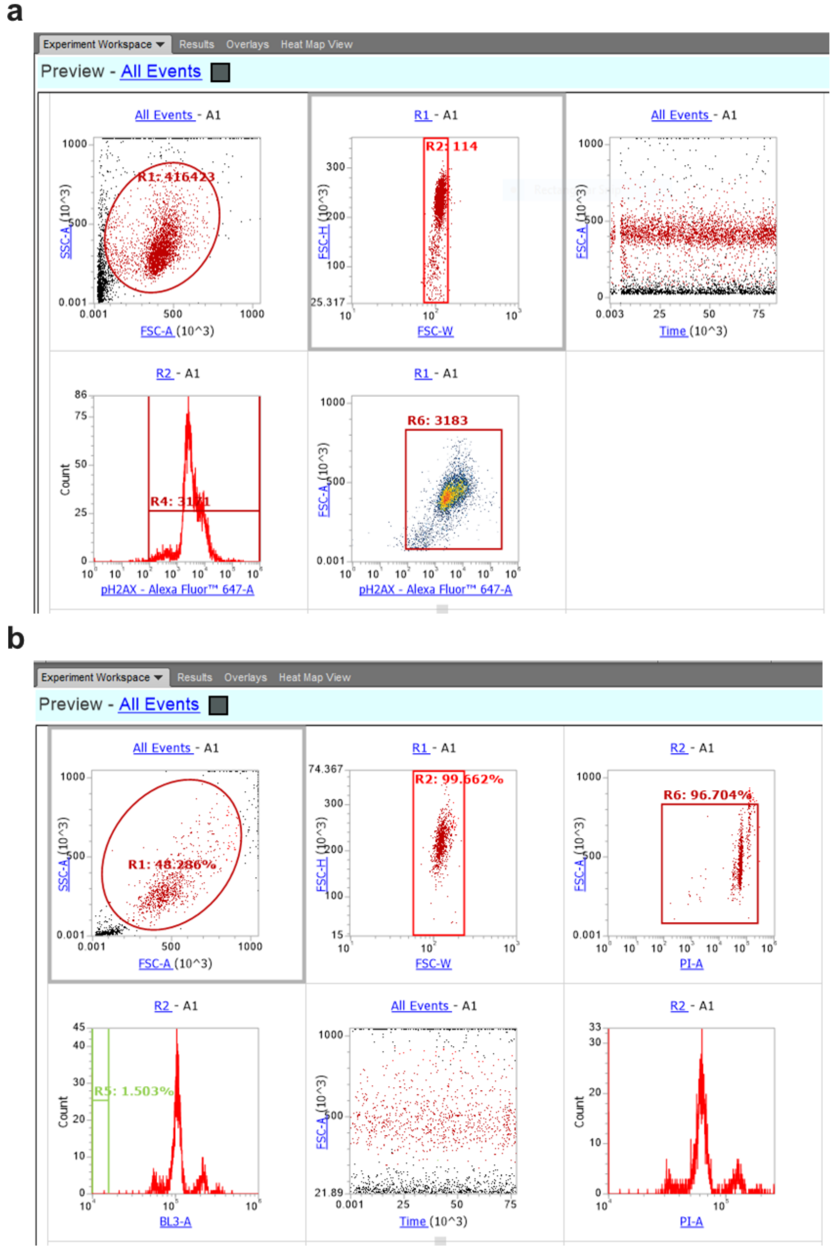


**Extended Figure 17,** Flow cytometry gating for γH2AX analysis (**a**) and cell cycle analysis (**b**).

**Table S1,** Alignment of insertional sequences discovered within the open reading frame of the Cas9 gene.

```
CLUSTAL O(1.2.4) multiple sequence alignment


B73_3           GGCAAAGCACTGAGAGATCCCCTCATAATTTCCCCAAAGCGTA---ACCATGTGT----G   53
B4387_1         TGCTAAGTCCTGAGAGATCCCCTCATAATTTCCCCAAAACGTA---ACCATGTGT----G   53
B4387_2         TGCTAAGTCCTGAGAGATCCCCTCATAATTTCCCCAAAACGTA---ACCATGTGT----G   53
A287_1          TACTCAATACTGATGAATCCCCTAATGATTTTTATCAAAATCATTAAGTTAAGGTAGATA   60
B73_1           TGCTAAGTCCTGATGAATCCCCTAATGATTTTTATCAAAATCATTAAGTTAAGGTAGATA   60
B73_2           TGCTAAGTCCTGATGAATCCCCTAATGATTTTTATCAAAATCATTAAGTTAAGGTAGATA   60
B4387_3         -CAAATAGCGCTACTAATCCCCTAATGATTTTGGTAAAAATCATTAAGTTAAGGTGGATA   59
                            *   ******* ** ****     **    *   *      **

B73_3           AATAAATTTTGAGCTAGT---AGGGTT-----------------GCAGCCACGAGTAAGT   93
B4387_1         AATAGATTTTGAGTAAGC---AGGGTT-----------------GCAGCCACGAGTGAGT   93
B4387_2         AATAGATTTTGAGTAAGC---AGGGTT-----------------GCAGCCACGAGTGAGT   93
A287_1          CACATCTTGTCATATGATCAAATGGTTTCGCCAAAAATCAATAATCAGACAACAAAATGT   120
B73_1           CACATCTTGTCATATGATCAAATGGTTTCGCCAAAAATCAATAATCAGACAACAAAATGT   120
B73_2           CACATCTTGTCATATGATCAAATGGTTTCGCCAAAAATCAATAATCAGACAACAAAATGT   120
B4387_3         CACATCTTGTCATATGATCAAATGGTTTCGCGAAAAATCAATAATCAGACAACAAGATGT   119
                 * *  ** * *         * ****                  *** **  *    **

B73_3           CTTCCCTTGTTATTG------------TGTAGCCAGAATGCCGCAAAACTTCCATGCCTA   141
B4387_1         CTTCCCTTGTTATTG------------TGTAGCCAGAATGCCGCAAAACTTCCATGCCTA   141
B4387_2         CTTCCCTTGTTATTG------------TGTAGCCAGAATGCCGCAAAACTTCCATGCCTA   141
A287_1          GCGAACTCGATATTTTACACGACTCTCTTTACCAATTCTGCCCCGAATTA--C--ACTTA   176
B73_1           GCGAACTCGATATTTTACACGACTCTCTTTACCAATTCTGCCCCGAATTA--C--ACTTA   176
B73_2           GCGAACTCGATATTTTACACGACTCTCTTTACCAATTCTGCCCCGAATTA--C--ACTTA   176
B4387_3         GCGAACTCGATATTTTACACGACTCTCTTTACCAATTCTGCCCCGAATTA--C--ACTTA   175
                     ** * ****             * ** * *   **** * **     *   * **

B73_3           AGCGAACTGTTGAGAGTACGTTTCGATTTC-TGACTGTGTTAGCCTGGAAGTGCTTGTCC   200
B4387_1         AGCGAACTGTTGAGAGTACGTTTCGATTTC-TGACTGTGTTAGCCTGGAAGTGCTTGTCC   200
B4387_2         AGCGAACTGTTGAGAGTACGTTTCGATTTC-TGACTGTGTTAGCCTGGAAGTGCTTGTCC   200
A287_1          AAACGACTCAACAGCTTAACGTTGGCTTGCCACGCCTTACTTGACTGTAAAA----CTCT   232
B73_1           AAACGACTCAACAGCTTAACGTTGGCTTGCCACGCCTTACTTGACTGTAAAA----CTCT   232
B73_2           AAACGACTCAACAGCTTAACGTTGGCTTGCCACGCCTTACTTGACTGTAAAA----CTCT   232
B4387_3         AAACGACTCAACAGCTTAACGTTGGCTTGCCACGCATTACTTGACTGTAAAA----CTCT   231
                *    ***    **  **   ** * ** *    *  *  * * *** **       **

B73_3           CAACCTTGTTTCTGAGCATGAACGCCCGCAAGCCAACATGTTAGTTGAAGCATCAGGGCG   260
B4387_1         CAACCTTGTTTCTGAGCATGAACGCCCGCAAGCCAACATGTTAGTTGAAGCATCAGGGCG   260
B4387_2         CAACCTTGTTTCTGAGCATGAACGCCCGCAAGCCAACATGTTAGTTGAAGCATCAGGGCG   260
A287_1          CAC---TCTTACCGAACTTGGCC----GTAACCTGCCA-------------ACCAAAGCG   272
B73_1           CAC---TCTTACCGAACTTGGCC----GTAACCTGCCA-------------ACCAAAGCG   272
B73_2           CAC---TCTTACCGAACTTGGCC----GTAACCTGCCA-------------ACCAAAGCG   272
B4387_3         CAC---TCTTACCGAACTTGGCC----GTAACCTGCCA-------------ACCAAAGCG   271
                **    * ** * ** * **  *    * ** *   **             * **  ***
```

```
B73_3          ATT-AGCAGCATGATATCAAAACGCTCTGAGCTGCTCGTTCGGCTATGGCGTAGGCCTAG   319
B4387_1        ATT-AGCAGCATGATATCAAAACGCTCTGAGCTGCTCGTTCGGCTATGGCGTAGGCCTAG   319
B4387_2        ATT-AGCAGCATGATATCAAAACGCTCTGAGCTGCTCGTTCGGCTATGGCGTAGGCCTAG   319
A287_1         AGAACAAAACATAACATCAAACGA--ATCGACCGATTGTTAGGTAATCGTCACCTCCACA   330
B73_1          AGAACAAAACATAACATCAAACGA--ATCGACCGATTGTTAGGTAATCGTCACCTCCACA   330
B73_2          AGAACAAAACATAACATCAAACGA--ATCGACCGATTGTTAGGTAATCGTCACCTCCACA   330
B4387_3        AGAACAAAACATAACATCAAACGA--ATCGACCGATTGTTAGGTAATCGTCACCTCCACA   329
               *      * *** * ******      *   * * * *** **  ** *      **

B73_3          TCCGTAGGCAGGACTTTTCAAGTCTCGGAAGGTTTCTTCAATCTGC--ATTCGCTTCGAA   377
B4387_1        TCCGTAGGCAGGACTTTTCAAGTCTCGGAAGGTTTCTTCAATCTGC--ATTCGCTTCGAA   377
B4387_2        TCCGTAGGCAGGACTTTTCAAGTCTCGGAAGGTTTCTTCAATCTGC--ATTCGCTTCGAA   377
A287_1         AAGAGCGACT-CGC-TGTATACCGTTGGCATGCTAGCTTTATCTGTTCGGGCAATACGAT   388
B73_1          AAGAGCGACT-CGC-TGTATACCGTTGGCATGCTAGCTTTATCTGTTCGGGCAATACGAT   388
B73_2          AAGAGCGACT-CGC-TGTATACCGTTGGCATGCTAGCTTTATCTGTTCGGGCAATACGAT   388
B4387_3        AAGAGCGACT-CGC-TGTATACCGTTGGCATGCTAGCTTTATCTGTTCGGGCAATACGAT   387
                     * *    * * *  *   * ** * * *   *  *****      *  * ***

B73_3          TAGATATTAACAAGTTGTTTGGGTGTTCGAATTTCAACAGGTAAGTTAGTTGCTAGAACC   437
B4387_1        TAGATATTAACAAGTTGTTTGGGTGTTCGAATTTCAACAGGTAAGTTAGTTGCTAGAATC   437
B4387_2        TAGATATTAACAAGTTGTTTGGGTGTTCGAATTTCAACAGGTAAGTTAGTTGCTAGAATC   437
A287_1         GCCCATTGTACTTGTTGAC---TGGTCTGATATCCGTGAGCAAAAACGGCTTATGGTATT   445
B73_1          GCCCATTGTACTTGTTGAC---TGGTCTGATATCCGTGAGCAAAAACGGCTTATGGTATT   445
B73_2          GCCCATTGTACTTGTTGAC---TGGTCTGATATCCGTGAGCAAAAACGGCTTATGGTATT   445
B4387_3        GCCCATTGTACTTGTTGAC---TGGTCTGATATTCGTGAGCAAAAACGACTTATGGTATT   444
                     *  **  ****       **  **  * *   **  **      *  * * *

B73_3          CATGGCTCCTTTGCCGACGCTGAGTAGATTTTAGGTGACGGGTGGTGACAATGAGTCCGT   497
B4387_1        CATGGCTCCTTTGCCGACGCTGAGTAGATTTTAGGTGACGGGTGGTGACAATGAGTCCGT   497
B4387_2        CATGGCTCCTTTGCCGACGCTGAGTAGATTTTAGGTGACGGGTGGTGACAATGAGTCCGT   497
A287_1         GCGAGCTTCAG---TCGCACTACACGGTCGTTCTGTTACTCTTTATGAGAAAGCGTTCCC   502
B73_1          GCGAGCTTCAG---TCGCACTACACGGTCGTTCTGTTACTCTTTATGAGAAAGCGTTCCC   502
B73_2          GCGAGCTTCAG---TCGCACTACACGGTCGTTCTGTTACTCTTTATGAGAAAGCGTTCCC   502
B4387_3        GCGAGCTTCAG---TCGCACTACACGGTCGTTCTGTTACTCTTTATGAGAAAGCGTTCCC   501
                   *** *        * **     *   **  ** **   *  *** ** * ** *

B73_3          G--TCGAGCGCTGATTTTT--------------------TCGGCCTTTAGAGCGAGATT   534
B4387_1        G--TCGAGCGCTGATTTTT--------------------TCGGCCTTTAGAGCGAGATT   534
B4387_2        G--TCGAGCGCTGATTTTT--------------------TCGGCCTTTAGAGCGAGATT   534
A287_1         GCTTTCAGAGCAATGTTCAAAGAAAGCTCATGACCAATTTCTAGCCGACCTTGCGAGCAT   562
B73_1          GCTTTCAGAGCAATGTTCAAAGAAAGCTCATGACCAATTTCTAGCCGACCTTGCGAGCAT   562
B73_2          GCTTTCAGAGCAATGTTCAAAGAAAGCTCATGACCAATTTCTAGCCGACCTTGCGAGCAT   562
B4387_3        GCTTTCAGAGCAATGTTCAAAGAAAGCTCATGACCAATTTCTAGCCGACCTTGCGAGCAT   561
               *  *  ** **    **                          ***      *****  *
```

```
B73_3           TATACAATAGA----------ATTTGGCATGAGATTGGATTGCTTTTAGTCAGCCTCTTA   584
B4387_1         TATACAATAGA----------ATTTGGCATGAGATTGGGTTGCTTTTAGTCAGCCTCTTA   584
B4387_2         TATACAATAGA----------ATTTGGCATGAGATTGGGTTGCTTTTAGTCAGCCTCTTA   584
A287_1          TCTACCGAGTAACACCACACCGCTCATTGTCAGTGATGCTGGCTTTAAAGT-----GCCA   617
B73_1           TCTACCGAGTAACACCACACCGCTCATTGTCAGTGATGCTGGCTTTAAAGT-----GCCA   617
B73_2           TCTACCGAGTAACACCACACCGCTCATTGTCAGTGATGCTGGCTTTAAAGT-----GCCA   617
B4387_3         TCTACCGAGTAACACCACACCGCTCATTGTCAGTGATGCTGGCTTTAAAGT-----GCCA   616
                * ***     *            *     * **    * * ***** *           *

B73_3           TAGCCTAAAGTCTTTGAGTGACTAGATGACATATCATGTAAGT---TGCTGATAGGTTTC   641
B4387_1         TAGCCTAAAGTCTTTGAGTGACTAGATGACATATCATGTAAGT---TGCTGATAGGTTTC   641
B4387_2         TAGCCTAAAGTCTTTGAGTGACTAGATGACATATCATGTAAGT---TGCTGATAGGTTTC   641
A287_1          TGGTATAAATCCGTTGAGAAGCTGGGTTGGTACTGGTTAAGTCGAGTAAGAGGAAAAGTA   677
B73_1           TGGTATAAATCCGTTGAGAAGCTGGGTTGGTACTGGTTAAGTCGAGTAAGAGGAAAAGTA   677
B73_2           TGGTATAAATCCGTTGAGAAGCTGGGTTGGTACTGGTTAAGTCGAGTAAGAGGAAAAGTA   677
B4387_3         TGGTATAAATCCGTTGAGAAGCTGGGTTGGTACTGGTTAAGTCGAGTAAGAGGAAAAGTA   676
                * *  ****  * *****   ** * *      *  *  *      *      *    *

B73_3           CAGTTTTCCGCTCCTAGGTCTGCATATTGTACTT-----TTCCTCTTACTCGACTTAACC   696
B4387_1         CAGTTTTCCGCTCCTAGGTCTGCATATTGTACTT-----TTCCTCTTACTCGACTTAACC   696
B4387_2         CAGTTTTCCGCTCCTAGGTCTGCATATTGTACTT-----TTCCTCTTACTCGACTTAACC   696
A287_1          CAATATGCAGACCTAGGAGCGGAAAACTGGAAACCTATCAGCAACTTACATGATATGTCA   737
B73_1           CAATATGCAGACCTAGGAGCGGAAAACTGGAAACCTATCAGCAACTTACATGATATGTCA   737
B73_2           CAATATGCAGACCTAGGAGCGGAAAACTGGAAACCTATCAGCAACTTACATGATATGTCA   737
B4387_3         CAATATGCAGACCTAGGAGCGGAAAACTGGAAACCTATCAGCAACTTACATGATATGTCA   736
                ** * * * *  *   *  * * * * ** *          *  *****  **  *  *

B73_3           AGTACCAACCCAGCTTCTCAACGGATTTATACCATGGCACTTTAAAGCCAGCATCACTGA   756
B4387_1         AGTACCAACCCAGCTTCTCAACGGATTTATACCATGGCACTTTAAAGCCAGCATCACTGA   756
B4387_2         AGTACCAACCCAGCTTCTCAACGGATTTATACCATGGCACTTTAAAGCCAGCATCACTGA   756
A287_1          --------TCTAGTCACTCAAAGACTTTAGGCTATAAGAGGC---------------TGA   774
B73_1           --------TCTAGTCACTCAAAGACTTTAGGCTATAAGAGGC---------------TGA   774
B73_2           --------TCTAGTCACTCAAAGACTTTAGGCTATAAGAGGC---------------TGA   774
B4387_3         --------TCTAGTCACTCAAAGACTTTAGGCTATAAGAGGC---------------TGA   773
                         * **   ***** *  ****  * **   *                  ***

B73_3           CAATGAGCGGTGTGGTGTTACTCGGTAGAATGCTCGCAAGGTCGGCTAGAAATTGGTCAT   816
B4387_1         CAATGAGCGGTGTGGTGTTACTCGGTAGAATGCTCGCAAGGTCGGCTAGAAATTGGTCAT   816
B4387_2         CAATGAGCGGTGTGGTGTTACTCGGTAGAATGCTCGCAAGGTCGGCTAGAAATTGGTCAT   816
A287_1          CTAAAAG------------------------CAACCCAATCTC---------ATGCCAAA   801
B73_1           CTAAAAG------------------------CAACCCAATCTC---------ATGCCAAA   801
B73_2           CTAAAAG------------------------CAACCCAATCTC---------ATGCCAAA   801
B4387_3         CTAAAAG------------------------CAATCCAATCTC---------ATGCCAAA   800
                * *  **                             ***  **          **   *
```

```
B73_3        GAGCTTTCTTTGAACATTGCTCTGAAAGCGGGAAC-------------------GCTTT   856
B4387_1      GAGCTTTCTTTGAACATTGCTCTGAAAGCGGGAAC-------------------GCTTT   856
B4387_2      GAGCTTTCTTTGAACATTGCTCTGAAAGCGGGAAC-------------------GCTTT   856
A287_1       TTCTATTGTATAAATCTCGCTCTAAAGGCCGAAAAAATCAGCGCTCGACACGGACTCATT   861
B73_1        TTCTATTGTATAAATCTCGCTCTAAAGGCCGAAAAAATCAGCGCTCGACACGGACTCATT   861
B73_2        TTCTATTGTATAAATCTCGCTCTAAAGGCCGAAAAAATCAGCGCTCGACACGGACTCATT   861
B4387_3      TTCTATTGTATAAATCTCGCTCTAAAGGCCGAAAAAATCAGCGCTCGACACGGACTCATT   860
                  ** * * **  * ***** ** ** * **                      * **

B73_3        CTCATAAAGAGTAACAGAACGACCGTGTAGT---GCGACTGAAGCTCGCAATACCATAAG   913
B4387_1      CTCATAAAGAGTAACAGAACGACCGTGTAGT---GCGACTGAAGCTCGCAATACCATAAG   913
B4387_2      CTCATAAAGAGTAACAGAACGACCGTGTAGT---GCGACTGAAGCTCGCAATACCATAAG   913
A287_1       GTCACCACCCGTCACCTAAAATCTACTCAGCGTCGGCAAAGGAGCCATGGATTCTAGCAA   921
B73_1        GTCACCACCCGTCACCTAAAATCTACTCAGCGTCGGCAAAGGAGCCATGGATTCTAGCAA   921
B73_2        GTCACCACCCGTCACCTAAAATCTACTCAGCGTCGGCAAAGGAGCCATGGATTCTAGCAA   921
B4387_3      GTCACCACCCGTCACCTAAAATCTACTCAGCGTCGGCAAAGGAGCCATGGGTTCTAGCAA   920
              ***  *   ** **  **   *     **    *  *  * ***      * * *  *

B73_3        TCGTTTTTGCTCACGAATATCAGACCAGTCAAC---AAGTACAATGGGCATCGTATTGCC   970
B4387_1      CCGTTTTTGCTCACGGATATCAGACCAGTCAAC---AAGTACAATGGGCATCGTATTGCC   970
B4387_2      CCGTTTTTGCTCACGGATATCAGACCAGTCAAC---AAGTACAATGGGCATCGTATTGCC   970
A287_1       CTAACTTACCTGTTGAAATTCGAACACCCAAACAACTTGTTAATATCTATTCGAAGC--G   979
B73_1        CTAACTTACCTGTTGAAATTCGAACACCCAAACAACTTGTTAATATCTATTCGAAGC--G   979
B73_2        CTAACTTACCTGTTGAAATTCGAACACCCAAACAACTTGTTAATATCTATTCGAAGC--G   979
B4387_3      CTAACTTACCTGTTGAAATTCGAACACCCAAACAACTTGTTAATATCTATTCGAAGC--G   978
                  **  **   * *  **  **     ***     **  *       *** *

B73_3        CGAACAGATAAAGCTAGCATGCCAACGGTATACAGCGAGTCGCT-CTTTGTG-GAGGTGA   1028
B4387_1      CGAACAGATAAAGCTAGCATGCCAACGGTATACAGCGAGTCGCT-CTTTGTG-GAGGTGA   1028
B4387_2      CGAACAGATAAAGCTAGCATGCCAACGGTATACAGCGAGTCGCT-CTTTGTG-GAGGTGA   1028
A287_1       AATGCAGATTGAAGAAACCTTCCGAGACTTGAAAAGTCCTGCCTACGGACTAGGCCTACG   1039
B73_1        AATGCAGATTGAAGAAACCTTCCGAGACTTGAAAAGTCCTGCCTACGGACTAGGCCTACG   1039
B73_2        AATGCAGATTGAAGAAACCTTCCGAGACTTGAAAAGTCCTGCCTACGGACTAGGCCTACG   1039
B4387_3      AATGCAGATTGAAGAAACCTTCCGAGACTTGAAAAGTCCTGCCTACGGACTAGGCCTACG   1038
                 *****  *   * * * ** *   *  * *     *  ** *    *  *

B73_3        CGATTACCTAACAATCG-GTCGATTCGT-TTGATGTTATGTTTTGTTCTCGCTTTGG---   1083
B4387_1      CGATTACCTAACAATCG-GTCGATTCGT-TTGATGTTATGTTTTGTTCTCGCTTTGG---   1083
B4387_2      CGATTACCTAACAATCG-GTCGATTCGT-TTGATGTTATGTTTTGTTCTCGCTTTGG---   1083
A287_1       CCATAGCCGAACGAGCAGCTCAGAGCGTTTTGATATCATGC-TGCTAATCGCCCTGATGC   1098
B73_1        CCATAGCCGAACGAGCAGCTCAGAGCGTTTTGATATCATGC-TGCTAATCGCCCTGATGC   1098
B73_2        CCATAGCCGAACGAGCAGCTCAGAGCGTTTTGATATCATGC-TGCTAATCGCCCTGATGC   1098
B4387_3      CCATAGCCGAACGAGCAGCTCAGAGCGTTTTGATATCATGC-TGCTAATCGCCCTGATGC   1097
             * **  ** *** * *   **    *** ***** * ***  *  *  ****  **
```

```
B73_3         ----------TTGGCAGGTTACGGCCAAGTTCGGTAAGAGTGAGAGTTTTACAGTCAAGT   1133
B4387_1       ----------TTGGCAGGTTACGGCCAAGTTCGGTAAGAGTGAGAGTTTTACAGTCAAGT   1133
B4387_2       ----------TTGGCAGGTTACGGCCAAGTTCGGTAAGAGTGAGAGTTTTACAGTCAAGT   1133
A287_1        TTCAACTAACATGTTGGCTTGCGGGCGTTCATGCTCAGAAACAAGGTTG---GGACAAGC   1155
B73_1         TTCAACTAACATGTTGGCTTGCGGGCGTTCATGCTCAGAAACAAGGTTG---GGACAAGC   1155
B73_2         TTCAACTAACATGTTGGCTTGCGGGCGTTCATGCTCAGAAACAAGGTTG---GGACAAGC   1155
B4387_3       TTCAACTAACATGTTGGCTTGCGGGCGTTCATGCTCAGAAACAAGGTTG---GGACAAGC   1154
                         **   * ** *** *      * * ***   *  ***     * ****

B73_3         AATGCGTGGCAAGCCAACGTTAAGCTGTTGAGTCGTTTTAAGTGTAATTCG-----GGGC   1188
B4387_1       AAGGCGTGGCAAGCCAACGTTAAGCTGTTGAGTCGTTTTAAGTGTAATTCG-----GGGC   1188
B4387_2       AAGGCGTGGCAAGCCAACGTTAAGCTGTTGAGTCGTTTTAAGTGTAATTCG-----GGGC   1188
A287_1        ACTTCCAGGCTAACACA--GTCAGAAATCGAAACGTACTCTCAACAGTTCGCTTAGGCAT   1213
B73_1         ACTTCCAGGCTAACACA--GTCAGAAATCGAAACGTACTCTCAACAGTTCGCTTAGGCAT   1213
B73_2         ACTTCCAGGCTAACACA--GTCAGAAATCGAAACGTACTCTCAACAGTTCGCTTAGGCAT   1213
B4387_3       ACTTCCAGGCTAACACA--GTCAGAAATCGAAACGTACTCTCAACAGTTCGCTTAGGCAT   1212
              *   *  *** * *  *   * **   * **  ***  *      * ****     *

B73_3         AGAATTGGTAAAGAGAGTCG--TGTAAAATATCGAGTTCGCACATCTTGTTGTCTGATTA   1246
B4387_1       AGAATTGGTAAAGAGAGTCG--TGTAAAATATCGAGTTCGCACATTTTGTTGTCTGATTA   1246
B4387_2       AGAATTGGTAAAGAGAGTCG--TGTAAAATATCGAGTTCGCACATTTTGTTGTCTGATTA   1246
A287_1        GGAAGTTTTGCGGCATTCTGGCTACACAATAACAAGGGAAGACTCACTCGTGGCTGCAAC   1273
B73_1         GGAAGTTTTGCGGCATTCTGGCTACACAATAACAAGGGAAGACTCACTCGTGGCTGCAAC   1273
B73_2         GGAAGTTTTGCGGCATTCTGGCTACACAATAACAAGGGAAGACTCACTCGTGGCTGCAAC   1273
B4387_3       GGAAGTTTTGCGGCATTCTGGCTACACAATAACAAGGGAAGACTTACTCGTGGCTGCAAC   1272
               *** *  *   *      *  *  * **** * **     **    *  ** ***

B73_3         TTGATTTTTCGCGAAACCATTTGATCATATGACAAGATGTGTATCCACCTTAACTTAATG   1306
B4387_1       TTGATTTTTGGCGAAACCATTTGATCATATGACAAGATGTGTATCTACCTTAACTTAATG   1306
B4387_2       TTGATTTTTGGCGAAACCATTTGATCATATGACAAGATGTGTATCTACCTTAACTTAATG   1306
A287_1        CC--TGCTTACTCAA----------------------AATCTATTCACA---CATGGTTA   1306
B73_1         CC--TGCTTACTCAA----------------------AATCTATTCACA---CATGGTTA   1306
B73_2         CC--TGCTTACTCAA----------------------AATCTATTCACA---CATGGTTA   1306
B4387_3       CC--TACTAGCTCAA----------------------AATTTATTCACA---CATGGTTA   1305
                  *  *     **                        * ***  **      *   *

B73_3         ATTTTTACCAAAATCATTAGGGGATTCATCAG      1338
B4387_1       ATTTTGATAAAAATCATTAGGGGATTCATCAG      1338
B4387_2       ATTTTGATAAAAATCATTAGGGGATTCATCAG      1338
A287_1        CGTTTTGGGGAAATTATGAGGGGATCTCTCAG      1338
B73_1         CGTTTTGGGGAAATTATGAGGGGATCTCTCAG      1338
B73_2         CGTTTTGGGGAAATTATGAGGGGATCTCTCAG      1338
B4387_3       CGCTTTGGGGAAATTATGAGGGGATCTCTCAG      1337
                 **     **** ** *******   ****
```

**Table S2. Bacterial strains**

| Strain | Relevant phenotype or genotype* | Source/Reference |
| --- | --- | --- |
| NEB5a | | New England Biolab |
| MIT A2 | *pks*$^{+}$ | Mouse isolate, from the James Fox lab at MIT |
| MIT A21 | *pks*$^{+}$ | Mouse isolate, from the James Fox lab at MIT |
| MIT guinea pig isolate | *pks*$^{+}$ | Guinea pig isolate, from the James Fox lab at MIT |
| NC101 | *pks*$^{+}$ | JANELLE C. ARTHUR, et al., 2012 |
| NC101$^{CmR}$ | *pks*$^{+}$ with a Cm resistance gene | This study |
| NC101 Δ*pks* | *pks* knockout | JANELLE C. ARTHUR, et al., 2012 |
| DH10B pBAC | pBAC vector containing the *pks* island | Jean-Philippe Nougayrède, et al., 2021 |
| DH10B pBAC-*pks* | Empty pBAC vector | Jean-Philippe Nougayrède, et al., 2021 |
| EC100Dpir$^{+}$ | F– mcrA Δ(mrr-hsdRMS-mcrBC) φ80dlacZΔM15 ΔlacX74 recA1 endA1 araD139 Δ(ara, leu)7697 galU galK λ– rpsL nupG pir+ (DHFR) | #ECP09500 (Lucigen) |

| | | |
|---|---|---|
| MG1655$^{NxR}$ | K-12 F– λ– ilvG– rfb-50 rph-1 Nxr | (Carraro et al., 2014) |
| *E. coli* Nissle 1917 strepR | *pks*$^{+}$ and Streptomycin resistance | Jean-Philippe Nougayrède, et al., 2021 |
| eB-TP114 | Evolved by accelerated laboratory evolution for 5 cycles in broth | This study |
| KN01 | Smr, Spr Nissle 1917 | Neil, et al., 2020 |

**Table S3: Replicative and conjugative plasmids**

| Plasmid | Relevant phenotype or genotype* | Source/Reference |
|---|---|---|
| pBXB1 | oriVpMB1, bxb1 integrase, bla (Apr) | Genbank: MK756311 (Neil et al., 2019) |
| pE-FLP | oriVpSC101ts, flp, Apr | Addgene #45978 |
| pGRG25: CmR | Tn7 machinery to insert the aad7 and cat genes in the glmS terminator region | This study |
| pKill1 | oriVpSC101ts, attPbxb1, FRT, 1 gRNA vs cat, aph-IIIa (Kmr), cas9 | Genbank: MK756312 (Neil et al., 2019) |
| pNative-Cas9 | Adapted from pKill1 | This study |
| pNative-dCas9 | oriVpSC101ts, attPbxb1, FRT, 1 gRNA vs cat, aph-IIIa (Kmr), dcas9 | This study |
| pBAD-dCas9 | oriVpSC101ts, attPbxb1, FRT, 1 gRNA vs cat, aph-IIIa (Kmr), dcas9, pBAD proter | This study |
| eB-TP114::tetB | eB-TP114Δaph-III::tetB | (Neil et al., 2021) |
| eB-TP114::pNative-Cas9 | eB-TP114::tetB with inserted wild-type Cas9 with the original native promoter after bxb1- and flp-mediated deletion of tetB and oriVpSC101ts | This study |
| eB-TP114::pNative-dCas9 | eB-TP114::tetB with inserted catalytically inactive Cas9 (dCas9) and the original native promoter after bxb1- and flp-mediated deletion of tetB and oriVpSC101ts | This study |
| eB-TP114::pBAD-dCas9 | eB-TP114::tetB with dCas9 and the arabinose inducible promoter, pBAD, after bxb1- and flp-mediated deletion of tetB and oriVpSC101ts | This study |

**Table S4: Cloning, qPCR and sequencing primers**

Capitalized letters in the sgRNA primers are the 20-nt spacer.

| Name | Sequence | Descriptions |
|---|---|---|
| clbP_F | tatcatctcctgtgctgtatgctg | qPCR |
| clbP_R | cctgcatccgttggtgaatt | qPCR |
| Universal 16S_F | ggtgaatacgttcccgg | qPCR |
| Universal 16S_R | tacggctaccttgttacgactt | qPCR |
| irp1_F1 | cactttctccgctaaggcca | qPCR |
| irp1_R1 | gcatcattcacgatccgcac | qPCR |
| irp1_F2 | agcacgttgtccatccagta | qPCR |
| irp1_R2 | cggcgaaccctgctatgtat | qPCR |
| irp1_F3 | aacagggtttcctcaccgtc | qPCR |
| irp1_R3 | ttgccagggctaaccttcag | qPCR |
| clbA_287_fwd | GGCTACTGTATCTATAGTATgttttagagctagaaatagcaagt | sgRNA primers |
| clbA_287_rev | ATACTATAGATACAGTAGCCgaatctattatacagaaaaattttc ctga | sgRNA primers |
| clbB_73_fwd | AATAAAAACCCCTGTTGACTgttttagagctagaaatagcaagt | sgRNA primers |
| clbB_73_rev | AGTCAACAGGGGTTTTTATTgaatctattatacagaaaaattttcc tga | sgRNA primers |
| clbB_1667_fwd | TGGCAACCACGGTGTTTACCgttttagagctagaaatagcaagt | sgRNA primers |
| clbB_1667_rev | GGTAAACACCGTGGTTGCCAgaatctattatacagaaaaattttc ctga | sgRNA primers |
| clbB_5400_fwd | TGTGCAAAATCGTGCAGCATgttttagagctagaaatagcaagt | sgRNA primers |
| clbB_5400_rev | ATGCTGCACGATTTTGCACAgaatctattatacagaaaaattttcc tga | sgRNA primers |
| clbB_3673_fwd | GGAATGGATATTCAGGATACgttttagagctagaaatagcaagt | sgRNA primers |
| clbB_3673_rev | GTATCCTGAATATCCATTCCgaatctattatacagaaaaattttcct ga | sgRNA primers |
| clbB_4387_fwd | GAACGCGATAGATCTATAGCgttttagagctagaaatagcaagt | sgRNA primers |
| clbB_4387_rev | GCTATAGATCTATCGCGTTCgaatctattatacagaaaaattttcc tga | sgRNA primers |
| clbB_5610_fwd | AAGGTAGAGAGCGTATTACCgttttagagctagaaatagcaagt | sgRNA primers |
| clbB_5610_rev | GGTAATACGCTCTCTACCTTgaatctattatacagaaaaattttcct ga | sgRNA primers |
| clbB_4129_fwd | GATACTGTCGCTATCAAAACgttttagagctagaaatagcaagt | sgRNA primers |
| clbB_4129_rev | GTTTTGATAGCGACAGTATCgaatctattatacagaaaaattttcc tga | sgRNA primers |

| | | |
|---|---|---|
| clbC_516_fwd | CTGGTGTAGAGACTATACACgttttagagctagaaatagcaagt | sgRNA primers |
| clbC_516_rev | GTGTATAGTCTCTACACCAGgaatctattatacagaaaaattttcctga | sgRNA primers |
| clbC_2085_fwd | TCAACGTCTTCTCTTATTTCgttttagagctagaaatagcaagt | sgRNA primers |
| clbC_2085_rev | GAAATAAGAGAAGACGTTGAgaatctattatacagaaaaattttcctga | sgRNA primers |
| clbC_2313_fwd | ACGAAAGGTACGCTTAACACgttttagagctagaaatagcaagt | sgRNA primers |
| clbC_2313_rev | GTGTTAAGCGTACCTTTCGTgaatctattatacagaaaaattttcctga | sgRNA primers |
| clbC_1481_fwd | CGTTTTCCACTTGTATCACTgttttagagctagaaatagcaagt | sgRNA primers |
| clbC_1481_rev | AGTGATACAAGTGGAAAACGgaatctattatacagaaaaattttcctga | sgRNA primers |
| clbH_1167_fwd | CCAGTTTTATACAGCCGCTCgttttagagctagaaatagcaagt | sgRNA primers |
| clbH_1167_rev | GAGCGGCTGTATAAAACTGGgaatctattatacagaaaaattttcctga | sgRNA primers |
| clbH_1297_fwd | CCCCGGGAAACGCATCGCATgttttagagctagaaatagcaagt | sgRNA primers |
| clbH_1297_rev | ATGCGATGCGTTTCCCGGGGgaatctattatacagaaaaattttcctga | sgRNA primers |
| clbI_331_fwd | ACCCACATCCCCCGGATAATgttttagagctagaaatagcaagt | sgRNA primers |
| clbI_331_rev | ATTATCCGGGGGATGTGGGTgaatctattatacagaaaaattttcctga | sgRNA primers |
| clbI_170_fwd | CTTTGGCCTTGACGTAATCTgttttagagctagaaatagcaagt | sgRNA primers |
| clbI_170_rev | AGATTACGTCAAGGCCAAAGgaatctattatacagaaaaattttcctga | sgRNA primers |
| clbJ_5578_fwd | GCCCCGACCGTAGGGAATACgttttagagctagaaatagcaagt | sgRNA primers |
| clbJ_5578_rev | GTATTCCCTACGGTCGGGGCgaatctattatacagaaaaattttcctga | sgRNA primers |
| clbJ_3688_fwd | CGTGCGCATACTCAGAGCATgttttagagctagaaatagcaagt | sgRNA primers |
| clbJ_3688_rev | ATGCTCTGAGTATGCGCACGgaatctattatacagaaaaattttcctga | sgRNA primers |
| clbL_412_fwd | TAATAGCTCGTTATCCCCTCgttttagagctagaaatagcaagt | sgRNA primers |
| clbL_412_rev | GAGGGGATAACGAGCTATTAgaatctattatacagaaaaattttcctga | sgRNA primers |
| clbL_654_fwd | CTGTTGATCTATCTCATGGTgttttagagctagaaatagcaagt | sgRNA primers |
| clbL_654_rev | ACCATGAGATAGATCAACAGgaatctattatacagaaaaattttcctga | sgRNA primers |
| clbL_993_fwd | CACCGCCACCGCCGCTCCTCgttttagagctagaaatagcaagt | sgRNA primers |
| clbL_993_rev | GAGGAGCGGCGGTGGCGGTGgaatctattatacagaaaaattttcctga | sgRNA primers |
| clbN_759_fwd | GCGCCAGCTTTCGTGTGTGGgttttagagctagaaatagcaagt | sgRNA primers |
| clbN_759_rev | CCACACACGAAAGCTGGCGCgaatctattatacagaaaaattttcctga | sgRNA primers |
| irp1_4019_fwd | AACCTTCCCTAGCCACTCCGgttttagagctagaaatagcaagt | sgRNA primers |
| irp1_4019_rev | CGGAGTGGCTAGGGAAGGTTgaatctattatacagaaaaattttcctga | sgRNA primers |

| | | |
|---|---|---|
| irp1_4129_fwd | GGCGCAGGAAGAGGCTCGGCgttttagagctagaaatagcaagt | sgRNA primers |
| irp1_4129_rev | GCCGAGCCTCTTCCTGCGCCgaatctattatacagaaaaattttcctga | sgRNA primers |
| irp1_397_fwd | GTCTTATGGGGGACGGCGCCgttttagagctagaaatagcaagt | sgRNA primers |
| irp1_397_rev | GGCGCCGTCCCCCATAAGACgaatctattatacagaaaaattttcctga | sgRNA primers |
| irp1_1595_fwd | CCCCGATTTCTCACCGGCCCgttttagagctagaaatagcaagt | sgRNA primers |
| irp1_1595_rev | GGGCCGGTGAGAAATCGGGGgaatctattatacagaaaaattttcctga | sgRNA primers |
| irp1_ko_fwd | aacgtgttctccggttgcgttgaggttgcgcaaagccagcgggtaatgtgatgggaattagccatggtcc | irp1 knockout |
| irp1_ko_rev | aacgcatggcgttccattgactttatgaaccttaggaaatgggaccgattgtgtaggctggagctgcttc | iIrp1 knockout |
| BACT1369F | cggtgaatacgttcycgg | qPCR |
| PROK1541R | aaggaggtgatccrgccgca | qPCR |
| clbB_F1 | acacccgctgcgttagagttgc | qPCR |
| clbB_R1 | ccgatggtcaaaggctgtgcgt | qPCR |
| clbC_F1 | gctggaaatatcgccggtcggg | qPCR |
| clbC_R1 | ttgcgcggattggggttttcca | qPCR |
| 1492R | tacggytaccttgttacgactt | 16S whole sequence |
| 27F | agagtttgatcmtggctcag | 16S whole sequence |
| aph-III upstream TP114 | ctatgccacgctacgccgctac | sequencing |
| aph-III downstream TP114 | ctggtcacggcggcgaatatcc | sequencing |
| ClbA upstream | tcacgctatacacattgctaacag | *pks*$^+$ detection |
| ClbA downstream | tgtgaatggcacgattatgcggg | *pks*$^+$ detection |
| pkill1 | gctcaacagtcacacatagacagcc | sequencing |

| | | |
|---|---|---|
| Cm spacer in original pKill1 | taacacgccacatcttgcga | sequencing |
| KO of clbP verification_dwn upstream | cgaatacggagcgacctcgatgc | *pks*$^{+}$ detection |
| KO of clbP verification_dwn | gtcagcgacggcatccaccatc | *pks*$^{+}$ detection |
| sgclbB_4387 | gctatagatctatcgcgttc | sgRNA detection |
| sgclbC_2313 | gtgttaagcgtacctttcgt | sgRNA detection |
| PstI_backbone | ggtgccctgaatgaactgca | Mutation of Cas9 to dCas9 |
| D10A | tgtgccgatagctaagcctat | Mutation of Cas9 to dCas9 |
| D10A | ataggcttagctatcggcaca | Mutation of Cas9 to dCas9 |
| H840A | tggaacaatggcatcgacatc | Mutation of Cas9 to dCas9 |
| H840A | gatgtcgatgccattgttcca | Mutation of Cas9 to dCas9 |
| BamHI_backbone | actatcaaaaccaccatatttttttg | Mutation of Cas9 to dCas9 |
| pBAD_Promoter_fwd | accccagcttcaaaagcgctatcgatgcataatgtgcc | Construction of pBAD_dCas9 |
| pBAD_Promoter_rev | attgagtatttcttatccatggtatatctccttattaaagttaaac | Construction of pBAD_dCas9 |
| pKILL1-cas9_Backbone_fwd | atggataagaaatactcaataggcttag | Construction of pBAD_dCas9 |
| pKILL1-cas9_Backbone_rev | agcgcttttgaagctggg | Construction of pBAD_dCas9 |

**Table S5. Promoter and coding DNA sequences**

| Name | Length | sequences |
|---|---|---|
| pNative promoter | 240 bp | tttatcagccataaaacaatacttaatactatagaatgataacaaaataaactactttttaaaagaattttgtgttata<br>atctatttattattaagtattgggtaatatttttgaagagatattttgaaaaagaaaaattaaagcatattaaactaat<br>ttcggaggtcattaaaactattattgaaatcatcaaactcattatggatttaatttaaactttttattttaggaggcaa<br>aa |
| pBAD promoter | 1346 bp | atcgatgcataatgtgcctgtcaaatggacgaagcagggattctgcaaaccctatgctactccgtcaagccgt<br>caattgtctgattcgttaccaattatgacaacttgacggctacatcattcactttttcttcacaaccggcacggaac<br>tcgctcgggctggccccggtgcattttttaaatacccgcgagaaatagagttgatcgtcaaaaccaacattgcg<br>accgacggtggcgataggcatccgggtggtgctcaaaagcagcttcgcctggctgatacgttggtcctcgcg<br>ccagcttaagacgctaatccctaactgctggcggaaaagatgtgacagacgcgacggcgacaagcaaacat<br>gctgtgcgacgctggcgatatcaaaattgctgtctgccaggtgatcgctgatgtactgacaagcctcgcgtac<br>ccgattatccatcggtggatggagcgactcgttaatcgcttccatgcgccgcagtaacaattgctcaagcagat<br>ttatcgccagcagctccgaatagcgcccttccccttgcccggcgttaatgatttgcccaaacaggtcgctgaaa<br>tgcggctggtgcgcttcatccgggcgaaagaaccccgtattggcaaatattgacggccagttaagccattcat<br>gccagtaggcgcgcggacgaaagtaaacccactggtgataccattcgcgagcctccggatgacgaccgta<br>gtgatgaatctctcctggcgggaacagcaaaatatcacccggtcggcaaacaaattctcgtccctgatttttca<br>ccacccccctgaccgcgaatggtgagattgagaatataacctttcattcccagcggtcggtcgataaaaaaatc<br>gagataaccgttggcctcaatcggcgttaaacccgccaccagatgggcattaaacgagtatcccggcagca<br>ggggatcattttgcgcttcagccatacttttcatactcccgccattcagagaagaaaccaattgtccatattgcat<br>cagacattgccgtcactgcgtcttttactggctcttctcgctaaccaaaccggtaaccccgcttattaaaagcatt<br>ctgtaacaaagcgggaccaaagccatgacaaaaacgcgtaacaaaagtgtctataatcacggcagaaaagt<br>ccacattgattatttgcacggcgtcacactttgctatgccatagcatttttatccataagattagcggatcctacct<br>gacgctttttatcgcaactctctactgtttctccatacccgtttttttgggctagccctgtagaaataattgtttaactt<br>taataaggagatatacc |
| Cas9 | 4107 bp | atggataagaaatactcaataggcttagatatcggcacaaatagcgtcggatgggcggtgatcactgatgaat<br>ataaggttccgtctaaaaagttcaaggttctgggaaatacagaccgccacagtatcaaaaaaaatcttataggg<br>gctcttttatttgacagtggagagacagcggaagcgactcgtctcaaacggacagctcgtagaaggtatacac<br>gtcggaagaatcgtatttgttatctacaggagatttttcaaatgagatggcgaaagtagatgatagtttctttcat<br>cgacttgaagagtcttttttggtggaagaagacaagaagcatgaacgtcatcctatttttggaaatatagtagat<br>gaagttgcttatcatgagaaatatccaactatctatcatctgcgaaaaaaattggtagattctactgataaagcgg<br>atttgcgcttaatctatttggccttagcgcatatgattaagtttcgtggtcatttttttgattgagggagatttaaatcct<br>gataatagtgatgtggacaaactatttatccagttggtacaaacctacaatcaattatttgaagaaaaccctatta<br>acgcaagtggagtagatgctaaagcgattctttctgcacgattgagtaaatcaagacgattagaaaatctcattg<br>ctcagctccccggtgagaagaaaaatggcttatttgggaatctcattgctttgtcattgggtttgacccctaatttt<br>aaatcaaattttgatttggcagaagatgctaaattacagctttcaaaagatacttacgatgatgatttagataattta<br>ttggcgcaaattggagatcaatatgctgatttgtttttggcagctaagaatttatcagatgctattttactttcagata<br>tcctaagagtaaatactgaaataactaaggctcccctatcagcttcaatgattaaacgctacgatgaacatcatc<br>aagacttgactcttttaaaagctttagttcgacaacaacttccagaaaagtataaagaaatcttttttgatcaatca<br>aaaaacggatatgcaggttatattgatgggggagctagccaagaagaattttataaatttatcaaaccaattta<br>gaaaaaatggatggtactgaggaattattggtgaaactaaatcgtgaagatttgctgcgcaagcaacggacct<br>ttgacaacggctctattccccatcaaattcacttgggtgagctgcatgctattttgagaagacaagaagactttta<br>tccatttttaaaagacaatcgtgagaagattgaaaaaatcttgacttttcgaattccttattatgttggtccattggc<br>gcgtggcaatagtcgttttgcatggatgactcggaagtctgaagaaacaattaccccatggaattttgaagaag<br>ttgtcgataaaggtgcttcagctcaatcatttattgaacgcatgacaaactttgataaaaatcttccaaatgaaaa<br>agtactaccaaaacatagtttgctttatgagtattttacggtttataacgaattgacaaaggtcaaatatgttactga<br>aggaatgcgaaaaccagcatttctttcaggtgaacagaagaaagccattgttgatttactcttcaaaacaaatcg<br>aaaagtaaccgttaagcaattaaaagaagattatttcaaaaaaatagaatgttttgatagtgttgaaatttcagga |

| | | |
|---|---|---|
| | | gttgaagatagatttaatgcttcattaggtacctaccatgatttgctaaaaattattaaagataaagatttttttggata<br>atgaagaaaatgaagatatcttagaggatattgttttaacattgaccttatttgaagatagggagatgattgagga<br>aagacttaaaacatatgctcacctctttgatgataaggtgatgaaacagcttaaacgtcgccgttatactggttg<br>gggacgtttgtctcgaaaattgattaatggtattagggataagcaatctggcaaaacaatattagatttttttgaaat<br>cagatggttttgccaatcgcaattttatgcagctgatccatgatgatagtttgacatttaaagaagacattcaaaa<br>agcacaagtgtctggacaaggcgatagtttacatgaacatattgcaaatttagctggtagccctgctattaaaaa<br>aggtattttacagactgtaaaagttgttgatgaattggtcaaagtaatggggcggcataagccagaaaatatcgt<br>tattgaaatggcacgtgaaaatcagacaactcaaaagggccagaaaaattcgcgagagcgtatgaaacgaat<br>cgaagaaggtatcaaagaattaggaagtcagattcttaaagagcatcctgttgaaaatactcaattgcaaaatg<br>aaaagctctatctctattatctccaaaatggaagagacatgtatgtggaccaagaattagatattaatcgtttaagt<br>gattatgatgtcgatcacattgttccacaaagtttccttaaagacgattcaatagacaataaggtcttaacgcgttc<br>tgataaaaatcgtggtaaatcggataacgttccaagtgaagaagtagtcaaaaagatgaaaaactattggaga<br>caacttctaaacgccaagttaatcactcaacgtaagtttgataatttaacgaaagctgaacgtggaggtttgagt<br>gaacttgataaagctggttttatcaaacgccaattggttgaaactcgccaaatcactaagcatgtggcacaaatt<br>ttggatagtcgcatgaatactaaatacgatgaaaatgataaacttattcgagaggttaaagtgattaccttaaaat<br>ctaaattagtttctgacttccgaaaagatttccaattctataaagtacgtgagattaacaattaccatcatgcccat<br>gatgcgtatctaaatgccgtcgttggaactgctttgattaagaaatatccaaaacttgaatcggagtttgtctatg<br>gtgattataaagtttatgatgttcgtaaaatgattgctaagtctgagcaagaaataggcaaagcaaccgcaaaat<br>atttcttttactctaatatcatgaacttcttcaaaacagaaattacacttgcaaatggagagattcgcaaacgccct<br>ctaatcgaaactaatggggaaactggagaaattgtctgggataaagggcgagattttgccacagtgcgcaaa<br>gtattgtccatgccccaagtcaatattgtcaagaaaacagaagtacagacaggcggattctccaaggagtcaa<br>ttttaccaaaaagaaattcggacaagcttattgctcgtaaaaaagactgggatccaaaaaaatatggtggttttg<br>atagtccaacggtagcttattcagtcctagtggttgctaaggtggaaaaagggaaatcgaagaagttaaaatcc<br>gttaaagagttactagggatcacaattatggaaagaagttcctttgaaaaaaatccgattgactttttagaagcta<br>aaggatataaggaagttaaaaaagacttaatcattaaactacctaaatatagtctttttgagttagaaaacggtcg<br>taaacggatgctggctagtgccggagaattacaaaaaggaaatgagctggctctgccaagcaaatatgtgaa<br>tttttatatttagctagtcattatgaaaagttgaagggtagtccagaagataacgaacaaaaacaattgtttgtgg<br>agcagcataagcattatttagatgagattattgagcaaatcagtgaattttctaagcgtgttattttagcagatgcc<br>aatttagataaagttcttagtgcatataacaaacatagagacaaaccaatacgtgaacaagcagaaaatattatt<br>catttatttacgttgacgaatcttggagctcccgctgcttttaaatattttgatacaacaattgatcgtaaacgatata<br>cgtctacaaaagaagttttagatgccactcttatccatcaatccatcactggtctttatgaaacacgcattgatttg<br>agtcagctaggaggtgactga |
| dCas9 | 4107 bp | atggataagaaatactcaataggcttaGCTatcggcacaaatagcgtcggatgggcggtgatcactgatga<br>atataaggttccgtctaaaaagttcaaggttctgggaaatacagaccgccacagtatcaaaaaaaatcttatag<br>gggctcttttatttgacagtggagagacagcggaagcgactcgtctcaaacggacagctcgtagaaggtata<br>cacgtcggaagaatcgtatttgttatctacaggagatttttttcaaatgagatggcgaaagtagatgatagtttcttt<br>catcgacttgaagagtcttttttggtggaagaagacaagaagcatgaacgtcatcctatttttggaaatatagtag<br>atgaagttgcttatcatgagaaatatccaactatctatcatctgcgaaaaaaattggtagattctactgataaagc<br>ggatttgcgcttaatctatttggccttagcgcatatgattaagtttcgtggtcatttttttgattgagggagatttaaat<br>cctgataatagtgatgtggacaaactatttatccagttggtacaaacctacaatcaattatttgaagaaaaccta<br>ttaacgcaagtggagtagatgctaaagcgattctttctgcacgattgagtaaatcaagacgattagaaaatctca<br>ttgctcagctccccggtgagaagaaaaatggcttatttgggaatctcattgctttgtcattgggtttgacccctaat<br>tttaaatcaaattttgatttggcagaagatgctaaattacagctttcaaaagatacttacgatgatgatttagataatt<br>tattggcgcaaattggagatcaatatgctgatttgtttttggcagctaagaatttatcagatgctattttactttcaga<br>tatcctaagagtaaatactgaaataactaaggctcccctatcagcttcaatgattaaacgctacgatgaacatca<br>tcaagacttgactcttttaaaagctttagttcgacaacaacttccagaaaagtataaagaaatctttttttgatcaatc<br>aaaaaacggatatgcaggttatattgatgggggagctagccaagaagaattttataaatttatcaaaccaatttta<br>gaaaaaatggatggtactgaggaattattggtgaaactaaatcgtgaagatttgctgcgcaagcaacggacct<br>ttgacaacggctctattccccatcaaattcacttgggtgagctgcatgctattttgagaagacaagaagactttta<br>tccatttttaaaagacaatcgtgagaagattgaaaaaatcttgacttttcgaattccttattatgttggtccattggc<br>gcgtggcaatagtcgttttgcatggatgactcggaagtctgaagaaacaattaccccatggaattttgaagaag |

| | | |
|---|---|---|
| | | ttgtcgataaaggtgcttcagctcaatcatttattgaacgcatgacaaactttgataaaaatcttccaaatgaaaa<br>agtactaccaaaacatagtttgctttatgagtattttacggtttataacgaattgacaaaggtcaaatatgttactga<br>aggaatgcgaaaaccagcatttctttcaggtgaacagaagaaagccattgttgatttactcttcaaaacaaatcg<br>aaaagtaaccgttaagcaattaaaagaagattatttcaaaaaaatagaatgttttgatagtgttgaaatttcagga<br>gttgaagatagatttaatgcttcattaggtacctaccatgatttgctaaaaattattaaagataaagatttttggata<br>atgaagaaaatgaagatatcttagaggatattgttttaacattgaccttatttgaagatagggagatgattgagga<br>aagacttaaaacatatgctcacctctttgatgataaggtgatgaaacagcttaaacgtcgccgttatactggttg<br>gggacgtttgtctcgaaaattgattaatggtattagggataagcaatctggcaaaacaatattagatttttgaaat<br>cagatggttttgccaatcgcaattttatgcagctgatccatgatgatagtttgacatttaaagaagacattcaaaa<br>agcacaagtgtctggacaaggcgatagtttacatgaacatattgcaaatttagctggtagccctgctattaaaaa<br>aggtattttacagactgtaaaagttgttgatgaattggtcaaagtaatggggcggcataagccagaaaatatcgt<br>tattgaaatggcacgtgaaaatcagacaactcaaaagggccagaaaaattcgcgagagcgtatgaaacgaat<br>cgaagaaggtatcaaagaattaggaagtcagattcttaaagagcatcctgttgaaaatactcaattgcaaaatg<br>aaaagctctatctctattatctccaaaatggaagagacatgtatgtggaccaagaattagatattaatcgtttaagt<br>gattatgatgtcgatGCCattgttccacaaagtttccttaaagacgattcaatagacaataaggtcttaacgcgt<br>tctgataaaaatcgtggtaaatcggataacgttccaagtgaagaagtagtcaaaaagatgaaaaactattgga<br>gacaacttctaaacgccaagttaatcactcaacgtaagtttgataatttaacgaaagctgaacgtggaggtttga<br>gtgaacttgataaagctggttttatcaaacgccaattggttgaaactcgccaaatcactaagcatgtggcacaa<br>attttggatagtcgcatgaatactaaatacgatgaaaatgataaacttattcgagaggttaaagtgattaccttaa<br>aatctaaattagtttctgacttccgaaaagatttccaattctataaagtacgtgagattaacaattaccatcatgcc<br>catgatgcgtatctaaatgccgtcgttggaactgctttgattaagaaatatccaaaacttgaatcggagtttgtcta<br>tggtgattataaagtttatgatgttcgtaaaatgattgctaagtctgagcaagaaataggcaaagcaaccgcaaa<br>atatttcttttactctaatatcatgaacttcttcaaaacagaaattacacttgcaaatggagagattcgcaaacgcc<br>ctctaatcgaaactaatggggaaactggagaaattgtctgggataaagggcgagattttgccacagtgcgcaa<br>agtattgtccatgccccaagtcaatattgtcaagaaaacagaagtacagacaggcggattctccaaggagtca<br>attttaccaaaaagaaattcggacaagcttattgctcgtaaaaaagactgggatccaaaaaaatatggtggtttt<br>gatagtccaacggtagcttattcagtcctagtggttgctaaggtggaaaaagggaaatcgaagaagttaaaat<br>ccgttaaagagttactagggatcacaattatggaaagaagttcctttgaaaaaaatccgattgactttttagaagc<br>taaaggatataaggaagttaaaaaagacttaatcattaaactacctaaatatagtctttttgagttagaaaacggt<br>cgtaaacggatgctggctagtgccggagaattacaaaaaggaaatgagctggctctgccaagcaaatatgtg<br>aatttttatatttagctagtcattatgaaaagttgaagggtagtccagaagataacgaacaaaaacaattgtttgt<br>ggagcagcataagcattatttagatgagattattgagcaaatcagtgaattttctaagcgtgttattttagcagatg<br>ccaatttagataaagttcttagtgcatataacaaacatagagacaaaccaatacgtgaacaagcagaaaatatt<br>attcatttatttacgttgacgaatcttggagctcccgctgcttttaaatattttgatacaacaattgatcgtaaacgat<br>atacgtctacaaaagaagttttagatgccactcttatccatcaatccatcactggtctttatgaaacacgcattgat<br>ttgagtcagctaggaggtgactga |
